# Exploring a diverse world of effector domains and amyloid signaling motifs in fungal NLR proteins

**DOI:** 10.1101/2022.03.16.484565

**Authors:** Jakub W. Wojciechowski, Emirhan Tekoglu, Marlena Gąsior-Głogowska, Virginie Coustou, Natalia Szulc, Monika Szefczyk, Marta Kopaczyńska, Sven J. Saupe, Witold Dyrka

## Abstract

NLR proteins are intracellular receptors constituting a conserved component of the innate immune system of multicellular organisms. In fungi, NLRs are characterized by high diversity of architectures and presence of amyloid signaling. Here, we explore the diverse world of effector and signaling domains of fungal NLRs using state-of-the-art bioinformatic methods including MMseqs2 for fast clustering, probabilistic context-free grammars for sequence analysis, and AlphaFold2 deep neural networks for structure prediction. In addition to substantially improving the overall annotation, especially in basidiomycetes, the study identifies novel domains and reveals the structural similarity of MLKL-related HeLo- and Goodbye-like domains forming the most abundant superfamily of fungal NLR effectors. Moreover, compared to previous studies, we found several times more amyloid motifs, including novel families, and validated aggregating and prion-forming properties of the most abundant of them *in vitro* and *in vivo*. Also, through an extensive *in silico* search, the NLR-associated amyloid signaling is for the first time identified in basidiomycetes. The emerging picture highlights similarities and differences in the NLR architectures and amyloid signaling in ascomycetes, basidiomycetes and other branches of life.

## Background

### NLR proteins

All living organisms possess an immune system allowing them to cope with viral or cellular pathogens. Among the central and conserved components of the innate immune system in animals and plants are the NLR proteins. NLRs are intracellular immune receptors that induces various host responses including regulated cell death upon the detection of non-self cues [1, 2, 3]. A typical NLR protein functions following a ligand-induced oligomerisation and activation process. Its tripartite domain architecture displays 1) a central Nucleotide-binding and Oligomerization Domain (NOD), 2) a C-terminal domain composed of superstructure forming repeats that is typically involved in detection of non-self cues in the form of DAMPs or MAMPs (Damage- or Microbe-Associated Molecular Patterns) and 3) a N-terminal effector domain whose activation induces various downstream host responses including regulation of the infected cell death [4, 5, 6, 7, 8]. While historically, NLRs were mostly studied within the animal and plant kingdoms (as Nod-Like Receptors and NBS-LRR Receptors respectively) [9, 10], their homologs were identified in bacteria and fungi [4].

In fungi, homologs of NLR proteins were initially identified in the context of the study of a non-self recognition process termed heterokaryon incompatibility (HI) [11]. This reaction occurs in filamentous fungi in the event of the fusion (anastasmosis) of the hyphæof genetically incompatible individuals, resulting in the death of mixed fusion cells [12, 13]. Incompatibility prevents in particular the transmission of mycoviruses between isolates during the anastomosis events. In *Podospora anserina*, HET-E, one of the proteins controlling heterokaryon incompatibility is a homolog of NLR proteins (although its N- and C-terminal domains differ from those known in animals and plants, a situation typical for NLR architecture proteins outside of the plant and animal kingdom [4, 14, 15]). Its central NOD domain is one of the original funding members used to define the NACHT domain (Pfam PF05729) common in animal NLRs (the H in the NACHT acronym stands for HET-E) [10, 16]. The C-terminal domain of HET-E protein, built of hypervariable WD40 repeats recognizes a non-self cue, here polymorphic variants of a host protein termed HET-C, a glycolipid transfer protein universally conserved in eukaryotes that could represent a pathogen effector target [17]. In such event, the N-terminal HET domain is activated which eventually leads to regulated cell death [17]. The HET domain (PF06985) is a cell death inducing domain with a remote homology to TIR domains [14]. Several other fungal cell death inducing incompatibility pathways in *Podospora* and other species are controlled by NLR proteins [18, 5]. Yet, apparently only a small fraction of the existing fungal NLRs appear involved in heterokaryon incompatibility and it is proposed that these proteins have more general function in immune defense and establishment of symbiotic interactions in fungi [19, 5]. Indeed, NLR proteins are abundant in multicellular filamentous fungi (no NLR protein was found in unicellular yeasts). In a recent study, a total of about 36 000 NLR proteins have been found in around 880 strains of over 560 species of fungi with on average 57 NLRs per genome and numerous species displaying hundreds of NLR genes [14, 5].

In terms of domain annotation fungal NLRs differ from their typical animal and plant counterparts. Unlike more homogenous NLR proteins in animals and plants, the central domain of fungal NLRs can be either of the NACHT [10] or the NB-ARC type (PF00931) [9]. Then fungal NLRs display ankyrin repeats (ANK, Pfam CL0465), tetratricopeptide repeats (TPR, CL0020) and beta-propellers of the WD40 meta-family (CL0186) in place of the LRR repeats found in most animal and plant NLRs. The NBS-TPR architecture was proposed to correspond to the ancestral architecture whilst NLR proteins in multicellular bacteria also typically display TPR, ANK or WD repeats [4, 14, 15, 20]. Consistent with a role in immune defense C-terminal repeated domains of fungal NLRs display marks of positive selection and are highly variable [21, 14, 18]. In addition, the C-terminal domains show original modes of functional diversification. First, about 1/6 of these C-terminal repeat domains consist of highly similar repeats with only a few highly variable positions under positive selection [22, 14]. These repeats arrays with high internal similarity are hypervariable loci in which individual repeats are exchanged and reshuffled resulting in functional diversification [21, 22]. High internal similarity of repeats is both a cause and a result of an unequal crossing over mechanism, a process which is 5-6 orders of magnitude faster than the point mutation [23]. Then, in the truffle *Tuber melanosporum* a superfamily of NACHT-ANK NLR encoding genes displays dozens of 3 bp miniexons whose alternative splicing can considerably diversify the repertoire of potential C-terminal recognition domain [24]. These strinking modes of recognition domain diversification are consistent with the proposed role of NLR proteins in the immune response, as capability of quickly adapting to evolving pathogens is a condition of success in the constant arms race against them [21].

For about 50% of fungal NLR proteins, N-terminal domain annotations could be determined with the Pfam [25] and similar HMM profiles [14], which make up for 12-13 major meta-families [14, 5]. Functionally, the characterized N-terminal domains belong to three basic types: enzymatic, signaling, and regulated cell death induction [26]. Out of 72 possible architectures made with the most common domain families (12 types of N-terminal domains, 2 types of central domains and 3 clans of C-terminal domains), as many as 32 were identified in fungal proteomes [14]. Interestingly, in about 20 cases, the closest orthologs of the central domain sequences were bound to different N-terminal domains (including in two different strains of the same species). Moreover, the maximum-likelihood phylogenetic trees generated separately for the N-terminal and central domains were mutually incompatible, and distribution of the N-terminal domains over the branches of central domains trees generated for selected species was scattered. Together with a relatively high number of NLRs without ortholog in other strains of the same species, these findings indicate high plasticity of the architecture of NLR proteins and the occurrence of the *death-and-birth evolution* process [14, 5].

### Amyloid signaling motifs

Another notable feature of fungal NLRs is the occurrence of amyloid-forming motifs at their N-termini [26]. A series of studies derived from the characterization of *Podospora anserina* [Het-s] prion protein, which controls regulated cell death in the context of heterokaryon incompatibility, has revealed that a fraction of the fungal NLR employ amyloid signaling to activate downstream cell death effector domains [27, 26]. The paradigmatic example of such amyloid NLR signalosomes is the HET-S/NWD2 two-component system of *P. anserina*. HET-S encodes a cell death execution protein with a globular N-terminal HeLo domain (PF14479) and a C-terminal amyloid forming prion domain composed of two elementary repeats r1 and r2 which are able adopt a specific *β*-solenoid amyloid fold [28, 29, 30, 31]. Amyloid transconformation of the C-terminal domain induces activation of the HeLo domain which turns into a pore-forming toxin. NWD2 is a NLR, encoded by the gene immediately adjacent to *het-S*, and displays at its N-terminus a motif termed r0 which is homologous to the elementary r1 and r2 repeats [32, 27]. When activated by their cognate ligand, engineered variants of NWD2 are capable of triggering transconformation of HET-S and to induce its toxicity. In this system, activation of the NLR leads to amyloid folding of its N-terminus which then serves as template to activate a cognate cell death execution protein [26].

The r0, r1, and r2 motifs, collectively referred to as the HET-s *motif*, represent one of the best studied examples of an amyloid signaling motif (ASM). Homologs of the HET-s motif can be grouped in 5 subclasses (collectively denoted as HET-s Related Amyloid Motifs or HRAM) [33], which co-occur in N-termini of fungal NLR proteins and in C-termini of HeLo [34, 35, 29] and HeLo-like (PF17111) proteins [32, 14] encoded by genes adjacent to NLR-encoding genes in the genome. In some organisms, two or three subclasses of HRAMs exist simultaneously, which allows for maintaining distinct signaling pathways [33, 36].

There are two other families of Fungal Amyloid Signaling Sequences (FASS) with similar functionality in the NLR protein system, namely sigma (named after the *σ* prion, which contains this motif [37]) and PP (pseudopalindromic due to the amino acid pattern NxGxQxGxN at its core) [32]. The PP motif bears significant resemblance to the mammalian RHIM motif [38, 39, 40] with remote homologs also in multicellular bacteria [20]. In addition, a recent *in silico* analysis of over 100,000 available bacterial genomes in search of sequence motifs repeated in adjacent genes encoding the Bell (bacterial homolog of fungal HeLo) and NLR proteins revealed ten families of Bacterial Amyloid Signal Sequences (BASS) widespread in multicellular Actinomycetes, Cyanobacteria and in Archaea [20]. Despite their sequence-level diversity, at least some if not all known BASS and FASS motifs are believed to share the beta-arch fold [41, 42, 43].

When compared to the NLR proteins in plant and animal kingdoms, the fungal NLR proteins display larger diversity of architectures. In addition, NLR-associated amyloid signaling appears specific to fungal and bacterial kingdoms although amyloid-motifs also occur in immune pathways in animals [44, 45]. The dominant view, until recently, was that the architecture and immunological function of NLR proteins in plants and animals resulted from the convergent evolution [15]. However, higher diversity of NLRs in fungi than in animals and plants, as well as presence of NLRs in multicellular prokaryotes [4, 20] suggest the early evolutionary origins of the architecture and the immune function of NLR proteins [5, 26]. Exploration of the diversity of fungal NLRs is an important asset for deciphering of the potential roles of these immune receptors in fungal biology in addition to their documented role in cell death related to incompatibility. In addition, comparative studies of NLRs in the different kingdoms can provide a more global view of the long term evolution of these central components of immunity in both microbes and macro-organisms. The aim of the current study is to improve the annotation and characterization of the vast ensemble of N-terminal domain of fungal NLRs with special focus on short domains (shorter than 150 amino acids) and amyloid-like motifs.

## Results

### Overview of N-terminal domains of fungal NLRs

In a previous study, we identified 36 141 NLR proteins in 487 fungal strains out of a total of 882 strains of 561 species for which the genome sequencing data was then available [20]. The ratio between the number of sequences from Ascomycota and Basidiomycota was 3.12:1 (27 152 to 8708). The N-terminus, as delimited according to NACHT or NB-ARC query match, was at least 20 amino-acids long in 32 962 proteins. Among these N-termini, 18 674 (57%) were annotated using either the Pfam [25] or our inhouse profiles [14]. In order to improve the annotation coverage, we clustered the set of N-termini with MMseqs2 [46] and then, for each cluster with more than 20 members (21 758 N-termini in 127 clusters, 15 105 already annotated), searched for homologs in UniRef30 [47, 48] and subsequently in Pfam using HHblits [49]. The clustering required mutual coverage of at least 80% of sequence length, and annotations were only assigned to sequences which overlapped at least 50% of the match to the Pfam profile. Moreover, only the matches including at least 50% of the Pfam profile length were retained to avoid excessive number of false positive hits. The procedure led to assigning the Pfam-based annotations to 3003 additional N-termini. This increased the annotation coverage from 69% to 82% (17 902 sequences) in the 20+ clusters, and from 57% to 66% (21 677 sequences) in the entire N-termini dataset.

The corpus of fungal NLR N-termini can be broadly divided into four main super-architectures (Fig. 1a). The first of them consists of the shortest N-termini, up to roughly 50 amino acid in length, which mainly comprise a direct N-terminal extension of the nucleotide binding domain. The second architecture adds the amyloid signaling motifs (ASM), typically made of 20–30 residues, which makes the entire N-terminus 70–120 amino-acids long. The third and the most frequent architecture consists of a single effector domain, while the fourth comprises with multiple domains. The length distribution of N-terminal sequences varies significantly with regard to the fungal phylum (Fig. 1b): while Basidiomycota are over-represented among short N-termini (below 100 amino acids), Ascomycota make up for 85% of domains longer than 200 amino acids.

**Figure 1:**
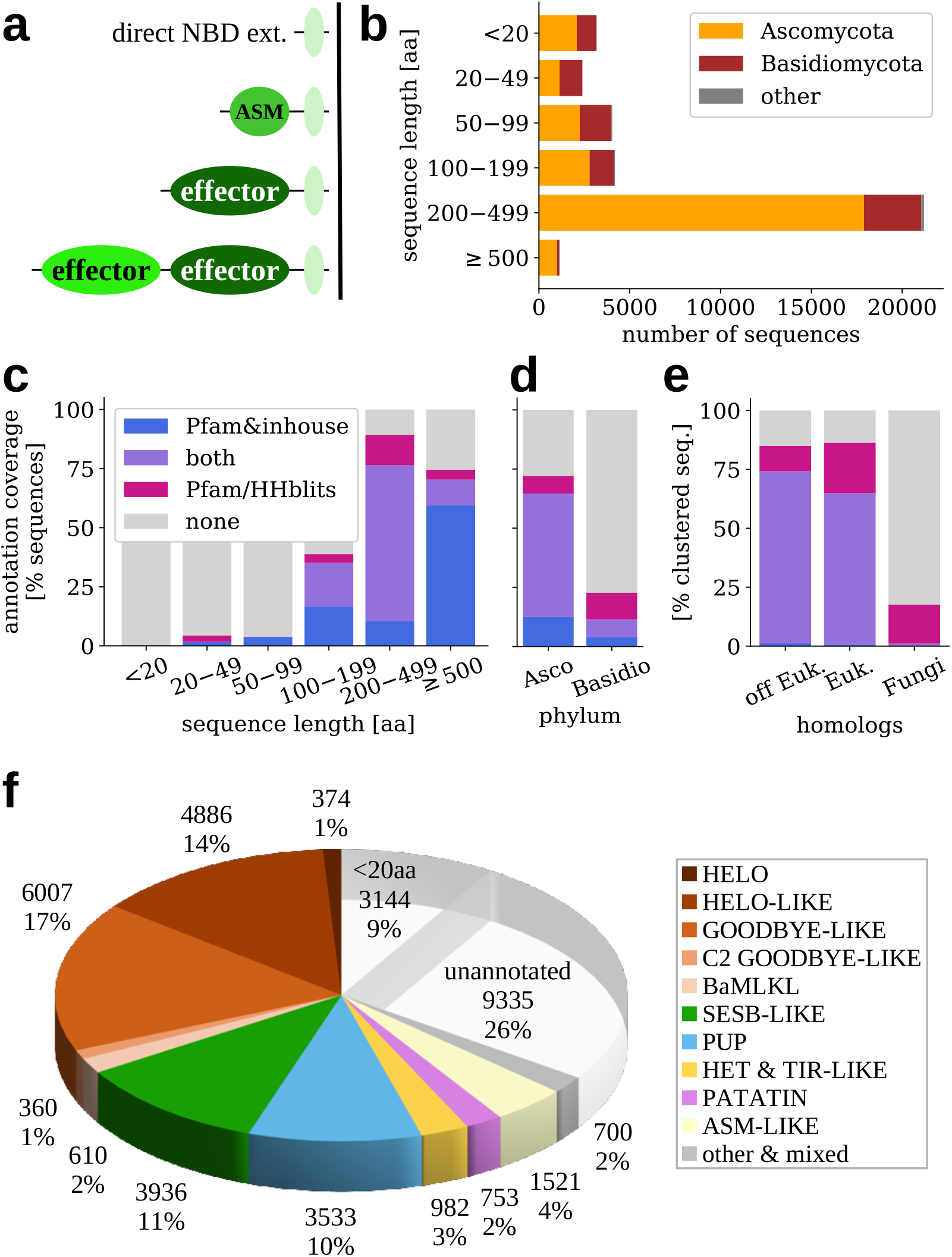
Fungal NLR N-termini. **a**) Major general archit ectures; **b**) Sequence length distribution; Annotation coverage with regard to **c**) sequence length, **d**) taxonomic division; **e**) Annotation coverage of clustered sequences (the 127 clusters, see main text) with regard to presence of taxonomically distant homologs in UniRef100 top hits. Euk. denotes *Eukaryota*. Colored bars indicate fraction of Pfam & inhouse annotated sequences (blue: only direct Pfam hits, violet: direct *and* with clustering & HHblits, rose: only with clustering & HHblits). Inhouse profiles were used only for direct Pfam searches. **f**) Distribution of domain families. Additional non-Pfam annotations included, see Results and Methods

The Pfam annotation coverage is not evenly distributed. While almost 90% of longer N-terminal domains (200 amino acids or more) are at least partially annotated, the figure is below 40% for the 100–200 amino acids range, and — not surprisingly — a few percent for domains shorter than 100 amino acids, which constitute 1/4 of all NLR N-termini (Fig. 1c). Also, sequences from Ascomycota are more completely annotated (72%) than Basidiomycota (23%) even though our the clustering-based annotation scheme increased coverage of the latter branch roughly twice (Fig. 1d). This inequality holds as well when N-termini in the same length ranges (above 100aa) are compared in both branches. When considering only 127 clusters from MMseqs2, Pfam annotations were found in more than 80% sequences with UniRef100 homologs outside the Fungi kingdom, but only in about 20% sequences with fungal-only homologs (Fig. 1e).

#### Major differences

Pfam annotations for fungal NLR N-termini are listed in Tab. 1 and summarized in Fig. 1f, and in Fig. S1 in Supplementary File 1. Vast majority of newly added annotations belonged to domain families already described as fungal NLR effectors (Tab. 1). In addition to domains reported in our previous surveys [14, 20], this involved also the Crinkler domain of the Ubiquitin clan only recently included in Pfam [50, 51, 52, 53, 54]. In our dataset the domain was found in two mycorrhizal species from two different phyla: *Serendipitia vermifiera* (Basidiomycota) and *Rhizophagus irregularis* (Mucormycota). The clustering-based search revealed also one new annotation (assigned to 24 sequences), the Sterile Alpha Motif family SAM_Ste50p [55]. SAM motifs are involved in homologous and heterologous protein-protein interactions [56].

**Table 1:**
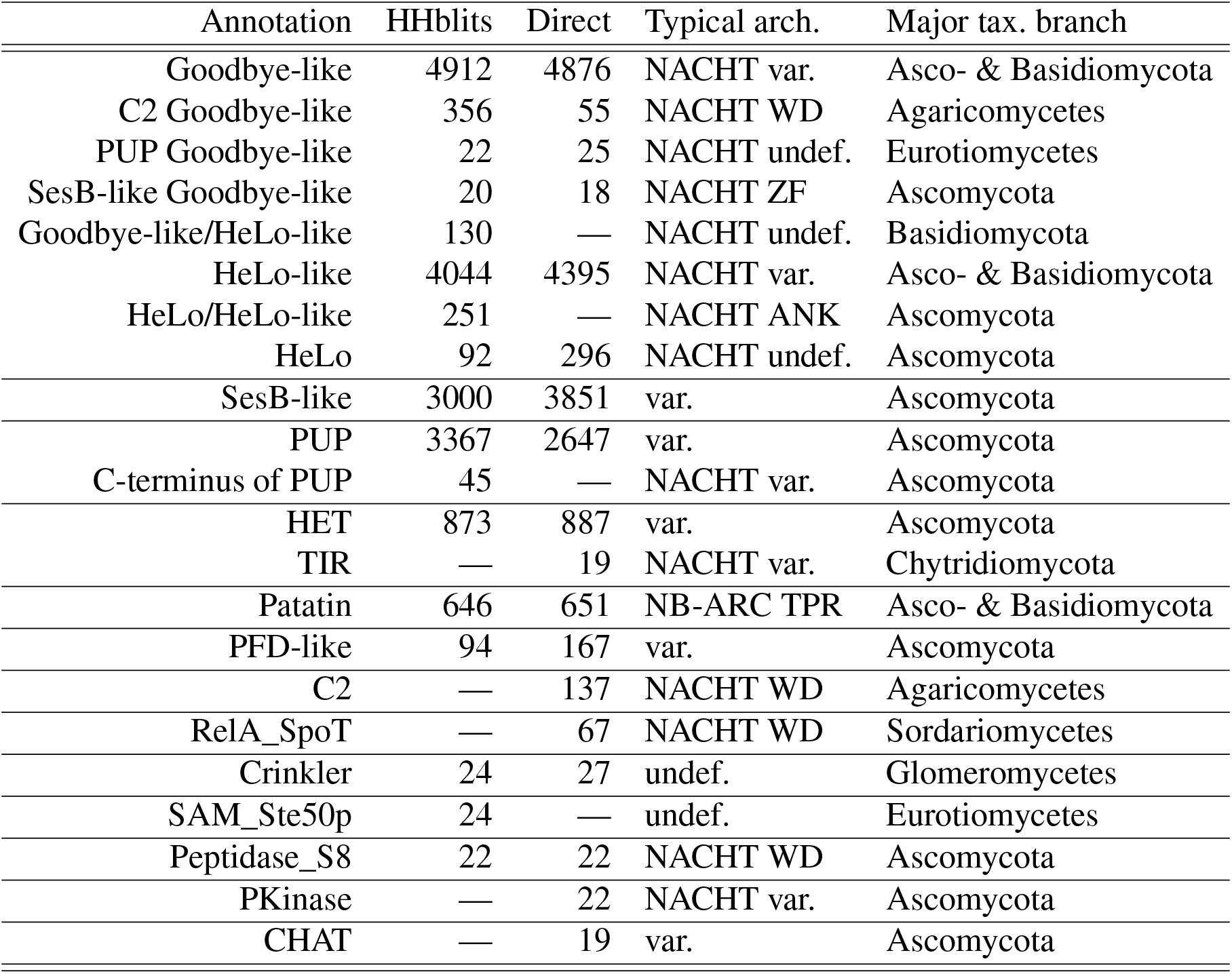
N-terminal annotations of fungal NLRs. Annotations based on Pfam [25] and inhouse [14] families and superfamilies. HHblits hits based on Pfam only. Direct hits include also inhouse annotations from [5]. Annotations are generally sorted by decreasing number of sequences. Homologous families are grouped together. Given are also the typical architecture of central and C-terminal domains if there is one clearly dominating, and the major taxonomic branch including vast <u>majority of the hits.</u>

On the other hand, no cluster matched the previously reported RelA_SpoT [57], PKinase [58], CHAT [59] and TIR [60, 61] profiles with sufficiently high hit coverage. TIR is a remote homologue of HET [14]. Interestingly, in the profile-sequence search PKinase was found to be N-terminally attached to another N-terminal domain in 29 cases, therefore it is not unlikely that the remaining single PKinase hits are also partial annotations. While no cluster was annotated as C2 only [62], a huge number of sequences fell into the C2 Goodbye-like class [32, 14] (356 hits), the architecture which is specific to Agaricomycetes. The Goodbye-like domain was found also in other double domain architectures of NLR N-termini (Tab. 1).

#### Major families

While the clustering-based approach found less SesB-like annotations than the profile-sequence search, this is likely due to the fact that the former procedure used only Pfam AB_Hydrolase clan [63] entries, while the latter method also relied on a sensitive inhouse SesB-like profile [14, 5]. On the contrary, the new approach increased the number of the Purine and Uridine Phosphorylase (PUP) superfamily annotations, mostly due to the matches to the purine NUcleoside Permease (NUP) profile [64]. In addition, 45 sequences from 27 various Pezizomycotina species comprised of about 50 amino-acid long C-terminus of PUP. The fragment forms an intrinsic part of the entire PNP_UDP_1 fold (cf. pdb:6po4B, residues 176–234). Both domain families are common in ascomycetal NLRs (13–14% each) but are virtually (SesB-like) or completely (PUP) missing from basidiomycotal NLRs (Fig. S1). The same is true of the HET domain present as N-terminal domain in ascomycetes but not in basidiomycetes.

Notably, we found clusters with apparently overlapping HeLo/HeLo-like and HeLo-like/Goodbye-like domain annotations. The latter situation was found in Basidiomycota and mostly involved sequences annotated as MLKL_NTD according to Conserved Domain Database (CDD) [65]. Human MLKL is an executioner domain homologous to fungal HeLo and bacterial Bell domains [32, 40, 20]. Moreover, there were additional basidiomycotal clusters with CDD MLKL_NTD annotation and/or with Pfam HeLo- or Goodbye-like annotations just below the assignment threshold, surmounting to a total of 600 basidiomycotal MLKL-like (BaMLKL) sequences. This makes the superfamily of Goodbye/HeLo/MLKL_NTD-like domains the most frequent in Basidiomycota (nearly 2000 sequences, around 1*/*4 of all), similarly to Ascomycota (Fig. S1).

Overall, the three most abundant domain classes, foremost the Goodbye/HeLo/MLKL-like followed by the SesB-like and PUP families, account for 60% of fungal NLR N-termini (Fig. 1f).

### Relation between HeLo-, Goodbye- and basidiomycotal MLKL-likes

To gain more insight in fungal MLKL-likes, we analyzed the largest cluster (OBZ65626, 106 sequences) with the overlapping Goodbye-like and HeLo-like annotations assigned through the HHblits-based procedure. Several sequences in the cluster received also hits from various MLKL-related Pfam profiles when sequences were searched individually (sequence and domain E-values of 1*e* − 3, Fig. 2a). Not surprisingly, the multiple sequence alignment of the cluster closely matched (HHpred [66, 67] probability above 98%) the sequence of human MLKL executioner domain with an experimentally solved spatial structure (pdb:6vzo [68], Fig. 2b). In fact the MLKL domain is virtually perfectly aligned with the Helo_like_N profile match, while the related SesA profile match is slightly shorter. At the same time, the matches to the two Goodbye-like profiles, Goodbye and NACHT_N [14], are both shifted N-terminally with regard to the MLKL-like domain resulting in a partial overlap, significantly longer for NACHT_N. Importantly, the multiple sequence alignment is well conserved for the combined stretch of Goodbye- and HeLo-like matches regardless of Pfam annotations of individual sequences (Fig. 2a).

**Figure 2:**
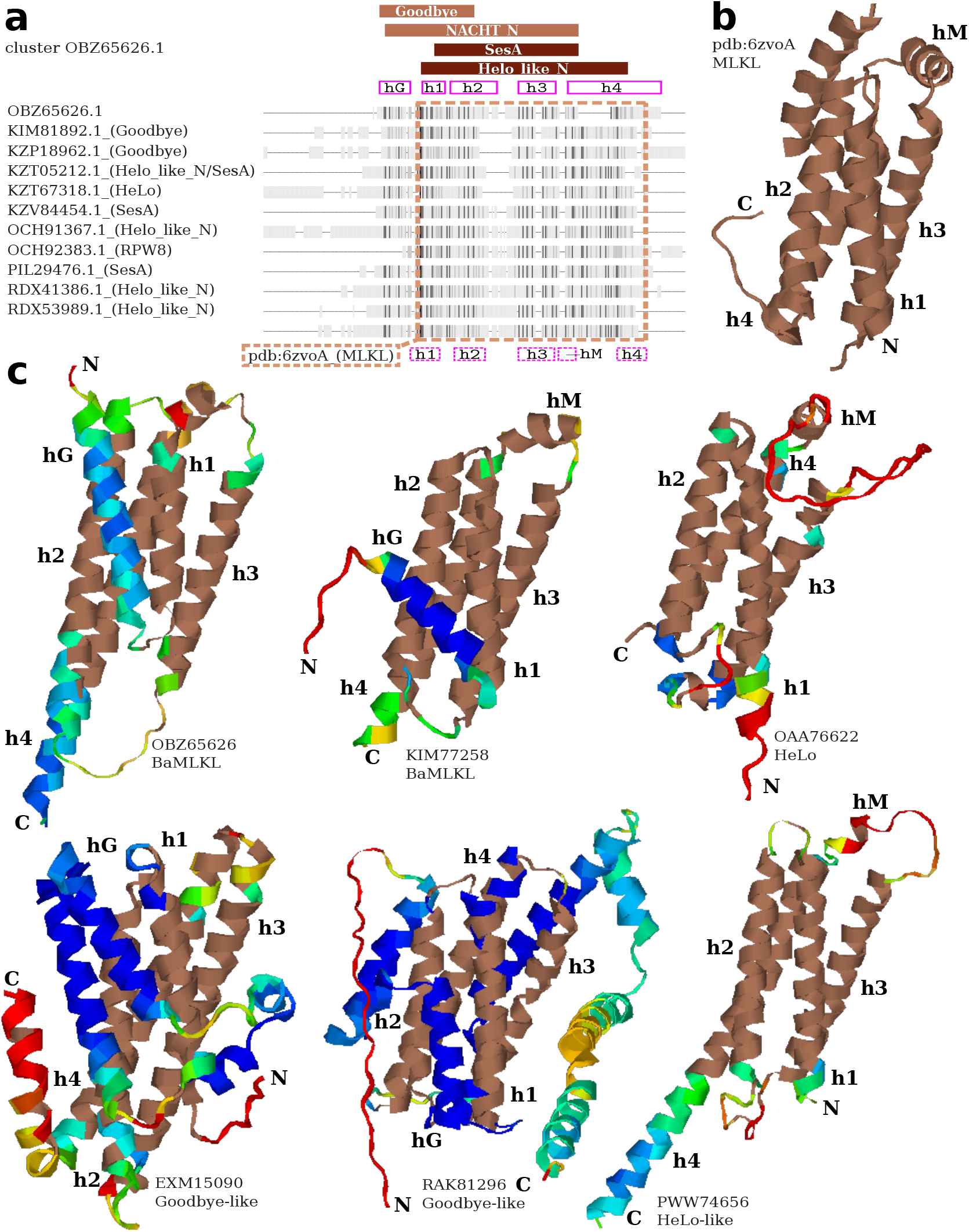
MLKL-like N-termini. (**a**) Fingerprint alignment of the doubly (Goodbye-like & Helo-like) annotated OBZ65626 cluster including non-redundant sequences with direct Pfam annotations. The alignment was truncated C-terminally. Darker shade implies higher conservation, while gaps are represented as lines. Columns matched with Pfam profiles of MLKL-like domains are indicated with brown bars. Columns corresponding to helices in a predicted OBZ65626 model are indicated with solid magenta boxes. Columns alignable to the human MLKL structure are framed with a brown dashed line. Columns corresponding to helices in the aligned MLKL structure are indicated with dashed magenta boxes. (**b**) The human MLKL structure (pdb:6zvoA). (**c**) Structural models of various MLKL-like domains predicted with AlphaFold2. Regions aligned to the human MLKL structure with TMalign are shown in brown. Rainbow colors indicate model quality in terms of lDDT (below or 50: red, 60: yellow, 70: green_3_,_9_80: cyan, above 90: blue).

Then, we attempted structure prediction for the largest MLKL-like clusters using AlphaFold2 [69] through the ColabFold advanced notebook [70]. The predictions were carried out solely using multiple sequences alignments of each cluster. Except for the largest HeLo-like cluster, all other predictions resulted in very good quality models (lDDT around 0.80) sharing a four-helix core (Fig. 2c), which is characteristic to the solved MLKL structure. When aligned to the latter using TMalign [71], the predicted models achieved TM-scores between 0.51 to 0.64. The most notable difference between structural models obtained for various clusters is an additional N-terminal helix in basidomycotal MLKL_NTD homologs and Goodbye-likes (hG in Fig. 2), not found in MLKL and HeLo-likes. However, Goodbye-like models present longer and more complex N-terminal extension than BaMLKLs. Also, Goodbye-likes lack a short perpendicular helix (hM) between helices h3 and h4, which seems to be a common feature of human and basidiomycotal MLKLs and HeLo-likes (Fig. 2c).

Taken together, these analyses indicate that although Goodbye-like profiles share a core region with the MLKL bundle and HeLo and Helo-like profiles, they also differ by the presence of an N-terminal extension ahead of the region corresponding to the first helix in MLKL/RPW8/HeLo proteins. Considering the critical role of this region in the membrane targeting activity of these animal, plant and fungal proteins, further experimental investigation are needed before a potential cell death inducing activity can be firmly attributed to Goodbye-like profiles [31, 72, 73].

### Unannotated longer N-termini

Roughly half of the 127 clusters with at least 20 member sequences did not get any Pfam annotation through the HHblits procedure. We carefully examined clusters with at least 10 non-redundant sequences (identity threshold of 70% or nr70) and median length above 100 amino acids. Apart from the MLKL-related clusters described above, there were five such clusters with a total of 195 sequences (127 at nr70) corresponding to four candidate domain families (Tab. 2). All five clusters were associated with NACHT NLRs.

**Table 2:** Unannotated N-terminal domains of fungal NLRs. For each cluster given are: an accession of its representative protein is given (Rep. acc.), proposed annotation label, number of sequences in cluster in total (#seq) and non-redundant at identity threshold of 70% (nr70), median sequence length (Len.), typical architecture of central and C-terminal domains (undef.—unannotated, var.—variable), and taxonomic distribution. NLR_PRDR stands for NLR effector domain with a Proline-Rich Disordered Region. See main text for details and Fig. S2 for structural models of <u>KEY84097, KFH66451 and PQE30996.</u>

| Rep. acc. | Prop. annot. | #seq (nr70) | Len. | Typical arch. | Tax. distrib. |
| --- | --- | --- | --- | --- | --- |
| KEY84097 | NLR_Helical | 67 (42) | 515 | NACHT TPR | Pezizomycotina |
| KFH66451 | NLR_Helical | 27 (19) | 617 | NACHT WD | Mortierellomycetes |
| XP_007333708 | NLR_PRDR | 41 (32) | 246 | NACHT undef. | Agaricaceae |
| KIL58680 | NLR_Koide | 31 (15) | 210 | NACHT WD | *Amanita muscaria* |
| PQE30996 | TIR-like | 29 (19) | 389 | NACHT var. | Pezizomycotina |

Two clusters (with a total of 94 sequences) consisted of homologous (HHalign [49] E-value of 6*e−* 44) relatively long domains (N-terminal length above 500 amino acids) from Pezizomycotina (e.g. KEY84097 protein from *Aspergillus fumigatus*) and Mortierellomycetes (e.g KFH66451 from *Podila verticillata*). Structure prediction with AlphaFold2 [69, 70] revealed the domain is made of multiple alpha-helices forming two stretches of the alpha solenoid-like structure (Fig. S2ab). NLRs with this N-terminal domain usually display C-terminal repeats of type HEAT (from the TPR clan) in Ascomycota, and WD40 in Mucormycota. Homologous proteins were also found (through the web-based profile HMM search) in bacteria, mainly in *Mycoavidus cysteinexigens*. As this betaproteobacteria is an endosymbiont of *Linnemania (Mortierella) elongata* AG-77, a fungus with the largest number of these proteins [2], this may suggest possibility of the horizontal gene transfer.

The second most abundant unnannoted cluster included 41 sequences (32 non-redundant at nr70) from *Agaricus bisporus* and *Leucoagaricus sp. SymC.cos* (representative protein XP_007333708 from *A. bisporus*), yet the profile HMM search revealed also single homologs in *Coprinopsis marcescibilis*, yeast *Saprochaete ingens* and protozoan parasite *Eimeria burnetti*. Interestingly, the latter species is a host of *Totiviridae* RNA virus similar to fungal viruses [74] reported also in Agaricaceae [75, 76]. These NLR N-termini (median length of 246 amino acids) consist of proline-rich disordered region and stretches of amyloid-like composition, including a conserved motif in C-terminus. The yeast homolog (non-NLR) is partially annotated as the glycogen recognition site of AMP-activated protein kinase AMPK1_CBM (PF16561) [77]. Due to the central disordered part, poorly alignable, no reliable structure prediction was possible.

Third most populated unannotated cluster consisted of 31 sequences (15 at nr70, representative protein KIL58680) with the median length of 210 amino acids, which are specific to the *Amanita muscaria* strain Koide BX008. In NLRs the domain is typically associated with C-terminal WD repeats. Again, no reliable structure was predicted.

The final candidate effector domain identified in this study was found in 29 sequences (19 at nr70) from various Pezizomycotina species (representative protein PQE30996 from *Rutstroemia sp.* NJR-2017a WRK4). The N-terminus has the median length of 389 amino acids and partially resembles the SEFIR family ([78, 79]) of TIR clan (HHblits hit probability of 90%). The TIR domain was reported in NLRs from plants, bacteria and Chytridiomycota [80, 20, 5]. The NLR proteins in this cluster are often associated with C-terminal repeats of Ankyrin, HEAT and WD40 types. In addition to the NACHT-based architectures present in the cluster, the web-based profile HMM search revealed several additional homologs in NLRs with the NB-ARC TPR domains. Interestingly, homologous domains are also present as separate proteins in Mucormycota *Rhizophagus irregularis*, a species related to *Mortierella*, and in association with NACHT WD40 and NACHT HEAT in *Mycoavidus cysteinexigens*, in accordance with the possibility of horizontal gene transfer [81]. A good quality structural model supports homology to TIR and HET domains (Fig. S2c).

### Amyloid-like motifs in short N-termini

The largest deficiency in the annotation coverage concerns short N-terminal sequences (length below 150 amino acids). Only a few percent of them (242 out of 3441) was annotated as so called prion-forming domains (PFD) [32, 14], consisting of the known fungal amyloid signaling sequences (FASS). However, many other fungal NLRs include short N-termini with a stretch of amino acids similar to already known signaling amyloids, which are characterized by a hydrophobic pattern typical to beta-sheets and frequent asparagines, glutamines and glycines. Moreover, the repertoire of already described FASS, merely three, is significantly smaller in comparison to bacterial amyloid signaling sequences (BASS) of which 10 families were recently identified in association to NLR proteins and the Bell domain [20]. Thus, in search for potential additional fungal amyloid signaling motifs we decided to thoroughly scan the short N-termini of NLRs with a probabilistic grammatical model inferred from ten families of bacterial amyloid signaling motifs (BASS) and shown to be sensitive to fungal amyloid signaling motifs [43].

In total, N-termini of 3441 sequences from 54 clusters with average sequence length up to around 150 amino acids were analyzed. Very high scoring fragments (see Methods) were found in 18 clusters with 1456 sequences. This included all 8 clusters (including 592 sequences) with at least one PFD-like annotation. The N-terminal sequences were made non-redundant at the identity level of 90% and submitted to motif extraction with MEME [82]. A total of 51 motifs were found for the E-value threshold of 1, for which profile HMMs were built in a two-stage procedure as in [20] (see Methods). Sequences of the 16 motifs were consistent with the grammatical model of BASS. These candidate amyloid-like motifs are presented in Fig. 3a. Eventually, their HMM profiles were used to scan all NLR N-termini at least 10 amino-acids long, comprising also sequences not included in the 127 clusters with 20 or more members (Tab. 3).

**Figure 3:**
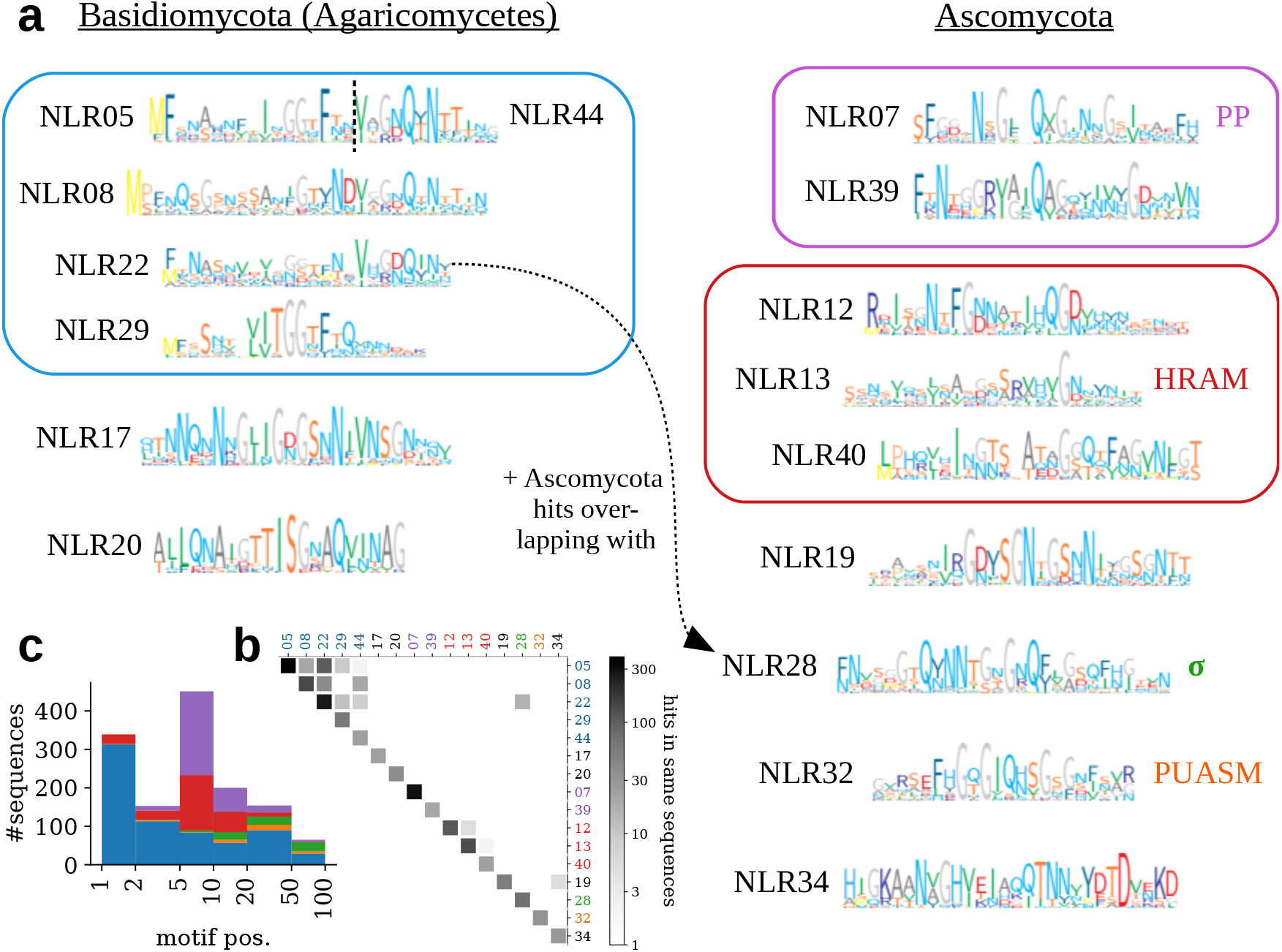
Amyloid-like motifs in short N-termini of NLRs. (**a**) Motif logos — grouped according to panel (**b**) Overlapping hits. See main text for details. (**c**) Stacked histogram of motif positions in NLR N-termini (the five main families of motifs color-coded as in panel (a)).

**Table 3:** Amyloid-like motifs in short N-termini of NLRs. Motif id indicates ranks in the MEME output. Motifs are grouped based on overlapping hits in NLRs and similar sequence patterns. Established and proposed motif annotation labels are given where applicable. #NLR and #nei. indicate number of sequences with a given motif in short N-termini of NLRs and C-termini of their genomic neighbors, respectively. AC+ indicates a proportion of motif instances for which ArchCandy score is 0.56 or above. Major taxonomic branch including vast majority of NLRs with the motif is given. #eff. indicates total number of single domain effector proteins (with established association to NLRs) with a given motifs in C-termini [5]. #cooc. indicates number of sequences with a given motif in short N-termini of NLRs / C-termini of effector proteins cooccurring in the same strains <u>(genome assemblies). #str. is a number of such strains with cooccurrence.</u>

| Mot. id | Annot. | Len. | #NLR | AC+ | Major tax. | #nei. | #eff. | #cooc. | #str. |
| --- | --- | --- | --- | --- | --- | --- | --- | --- | --- |
| NLR05 | — | 17 | 387 | 0.28 | Agaricomycetes | 0 | 0 | — | — |
| NLR08 | — | 30 | 138 | 0.55 | Agaricomycetes | 0 | 0 | — | — |
| NLR22 | — | 24 | 263 | 0.53 | Agaricomycetes | 0 | 0 | — | — |
| NLR29 | — | 20 | 60 | 0.53 | Agaricales | 0 | 0 | — | — |
| NLR44 | — | 12 | 24 | 0.00 | Agaricomycetes | 0 | 0 | — | — |
| NLR07 | PP | 23 | 296 | 0.64 | Ascomycota | 37 | 112 | 128/58 | 45 |
| NLR39 | PP | 24 | 20 | 0.80 | Ascomycota | 0 | 1 | 0/0 | 0 |
| NLR12 | HRAM3 | 26 | 110 | 0.48 | Sordariomycetes | 4 | 24 | 80/14 | 13 |
| NLR13 | HRAM | 24 | 131 | 0.57 | Ascomycota | 16 | 50 | 37/21 | 18 |
| NLR40 | HRAM | 26 | 24 | 0.92 | Ascomycota | 0 | 0 | — | — |
| NLR17 | — | 26 | 24 | 0.88 | *A. muscaria* | 0 | 0 | — | — |
| NLR19 | — | 27 | 53 | 0.60 | *Tuber* | 0 | 0 | — | — |
| NLR20 | — | 21 | 37 | 0.68 | Agaricales | 0 | 0 | — | — |
| NLR28 | $\sigma$ | 28 | 71 | 0.80 | Ascomycota | 42 | 90 | 42/40 | 37 |
| NLR32 | *PUASM* | 22 | 33 | 0.45 | Sordariomycetes | 11 | 55 | 25/21 | 18 |
| NLR34 | — | 28 | 29 | 0.52 | Ascomycota | 0 | 0 | — | — |

Not surprisingly, some of the 16 motifs clearly corresponded to the three FASS families: HRAM (NLR13, found in 131 sequences), PP (NLR07, 296), *σ* (NLR28, 71). The overall recall of 498 hits is twice higher in comparison to the combined Pfam-based approaches (242). Several hits of another two motifs, NLR12 and NLR40, overlapped with the NLR13 (HRAM) matches (Fig. 3b). Moreover, the HMM scan with a generalized HRAM profile based on HRAM dataset from [33] recognized 1/3 of NLR12 and 2/3 of NLR40 motifs at E-value of 0.01, thus indicating that these two classes are related to HRAM. Indeed, the NLR12 motif (Fig. 3a) is apparently similar to HRAM3 [33]. In addition, the G-hydrophobic-Q-hydrophobic-G pattern of NLR39 motif resembles NLR07 (PP). Five other motifs (NLR17, NLR19, NLR20, NLR32 and NLR34, in 138 sequences altogether) were difficult to assign to the known families. The final and the largest subgroup (689 sequences) consisted of five motifs (NLR05, NLR08, NLR22, NLR29 and NLR44) with hits substantially overlapping NLR22 hits. This large group is specific to basidiomycetes except of a dozen of NLR22 hits overlapping ascomycotal NLR28 (*σ*) (Fig. 3b). While most motifs are distributed in larger taxonomic branches, two motifs are more restricted: NLR17 is specific to *Amanita muscaria* (strain Koide) and NLR19 to genus *Tuber*. A combined NLR19 + NLR34 configuration was found in five highly homologous sequences from *Tuber melanosporum* (Fig. 3b).

Significant numbers of similar motifs in C-termini (100aa) of genomically neighboring (20kbp) proteins were found only for motifs representing the three FASS families (NLR07 in 37 sequences, NLR12 in 4, NLR13 in 16, and NLR28 in 42) and for NLR32 (11 sequences). This suggests that NLR32 defines a new family of amyloid signaling motifs. (For further computational and experimental verification, see below.)

Notably, the amyloid-like motifs differ in their position in NLR N-termini. While motifs from the NLR05/NLR22 group are usually situated in the very terminus, most HRAMs (NLR12/13/40) and PPs (NLR07/39) are located at positions 5–9. Moreover, NLR32 and *σ* motifs (NLR28) are shifted further C-terminally with relative majority at positions 20–49 and 50–99, respectively (Fig. 3c). In addition, a couple of dozens of amyloid-like motifs of various families (including 17 NLR05 and 7 NLR07) were found located centrally or C-terminally in longer N-termini. Some of them form combined architectures with annotated domains, most notably with NLR_PRDR (NLR05 in 10 sequences from *A.bisporus*) and MLKL-likes (5 BaMLKL + NLR05 in *Laccaria bicolor*, 4 HeLo-like + NLR28 and 1 HeLo-like + NLR07 in various Ascomycota).

Regardless of the evidence of amyloid signaling, all 16 motifs are likely to assume the beta-arch fold typical to known FASS and BASS as from 45 to 95% motif instances pass the fold prediction threshold of ArchCandy (0.56). The only exceptions are two shortest motifs NLR05 (28%) and NLR44 (none), probably because they comprise only parts of the actual amyloid-like motif (Fig. 3a). For three motif profiles matches were found in the PDB database [83], namely pdb:3erb [84] for NLR05/NLR22, and pdb:4q2w:A [85] for NLR34. The both solved proteins are hydrolases and the matched fragments possess a beta-arch-like fold.

### A reverse approach: amyloid-like motifs in C-termini of effector proteins

In order to complement the search for amyloid signaling motifs in NLRs and verify discovery of the fourth NLR-related FASS family, we adapted the approach recently used for identification of 10 families of BASS in NLR-related proteins in bacteria [20].

We iteratively searched for remote homologs (using HMMER [86]) of effector domains related to NLR proteins, starting with 19 Pfam profiles of N-terminal domains of NLRs reported in [5] (see Methods). We found almost 140,000 sequences with the single domain architecture and C-terminus between 10 to 150 amino acids in length. For each domain family separately, the C-termini were made non-redundant at identity of 70% (using CD-HIT [87, 88]) and searched for short (10–30 amino acids) motifs using MEME [82]. This resulted in around 800 motifs, for which profile HMMs were built, as in the previous section (see Methods). In addition, we extracted all fungal NACHT and NB-ARC proteins (according to Pfam) with N-termini between 10 and 150 amino acids. The N-termini were then searched for MEME motifs, for which, eventually, 49 profile HMMs were generated. NLR N-termini and effector C-termini were scanned with the combined set of effector- and NLR-side motifs, and the hits in both groups of proteins (at E-value of 1*e−* 2) were matched based on genomic proximity (up to 20kbp) of genes encoding the proteins. Eventually, for 19 motifs, at least 3 non-redundant pairs of motif instances were found.

These motifs were clustered on the basis of their co-occurrence in pairs of sequences, with three clusters corresponding to the already known families PP, *σ*, and HRAM (Fig. S3). Two additional motifs with few pairs apparently resemble HRAM2 and HRAM4 [33], respectively, and one another motif resembles the C-terminal part of the *σ* family motifs (Fig. S3). The fourth largest family of motifs exhibits a distinctive conserved pattern FxGxGxQxxGxGxF, which clearly corresponds to the NLR32 motif in Fig. 3. Since in both searches the motif was found associated uniquely with the PNP_UDP domain, we term it PUASM, or the Pnp_Udp-associated Amyloid Signaling Motif. The NLRs with the PUASM motif proteins are annotated either as NACHT or NACHT WD40. All matched instances of the PUASM motif come from various Pezizomycotina species. Finally, we found one more distinct motif related to PNP_UDP, however only present in four pairs (PF01048_040 in Fig. S3).

Overall, the ASM differ in type of associated effector domain, either pore-forming (HeLo and HeLo-like for HRAM/NLR13), enzymatic (PNP_UDP for HRAM/NLR12, NLR32 and PF01048_040), or both (PP/NLR07 and *σ* /NLR28). Interestingly, while the NLR13 motif was typically found as a double in C-termini of HeLo and HeLo-like domains, for the second HRAM-like, NLR12, only single instances were found in C-termini of PNP_UDP_1 effector proteins. This suggests a different mode of operation despite their similar sequence profiles.

To check the possibility that proteins cooperating through the amyloid signaling are encoded by sequentially distant genes, we analyzed co-occurrence of particular amyloid-like motifs in N-termini of NLRs and C-termini of previously described effector domains [5] in entire genomes. Non-singular C-terminal hits and genomic co-occurrences were found only for FASS and NLR32 (Tab. 3).

### Amyloid-like motifs in Basidiomycota

With the NLR-related amyloid signaling previously described in multicellular bacteria and Ascomycota, apparent is the lack of evidence of this mechanism in Basidiomycota. On the other hand, we found numerous homologs of the pore-forming HeLo and HeLo-like domains in Basidiomycotal NLRs. Thus, we used them for searching the entire Basidiomycota genomes for homologs separate from NLR domains (web-based profile HMM search with standard parameters). We identified hundreds of such putative singular pore-forming domains, which — because of their potential to cause the cell death — can be expected to be under control of other proteins. As in Ascomycota such control is exerted by NLRs through the amyloid signaling motifs, we scanned the identified MLKL_NTD homologs against ASM motif profiles and grammars. However, motifs resembling ASMs were identified only in a few out of 500 sequences and no associated motifs were found in the neighboring NLRs. Yet in two cases the motif pairs occurred when entire genomes were considered (Fig. 4a). In *Moniliophthora roreri* (strain MCA 2997) there is a 18 amino-acid long motif apparently shared between two MLKL_NTD-like C-termini and 26 short NACHT N-termini (Fig. S4). In addition in *Fibularhizoctonia* sp. CBS 109695 there is a conserved pattern shared between two MLKL_NTD-like C-termini, eight short NLR N-termini (including KZP25847 with NLR20 instance) and additional five NLR proteins where the pattern is situated between MLKL_NTD-like and NACHT domains (including KZP30127 and KZP3012 with NLR22 instances), see alignment in Fig. S5. It would suggest a possibility that in *Fibularhizoctonia* proteins with the N-terminal and C-terminal amyloid-like motifs are pseudogenes, especially that three NLRs in this group are untypically short (less than 200 amino acids). However, NLRs with N-terminal and mid-sequence ASMs differ in domain configuration with the former belonging to NACHT, NACHT ANK and NACHT VHS architectures, while the latter are all of the NACHT TPR type (Fig. 4a). In turn, for *M. rorei*, we found only one protein with the MLKL_NTD + AAA_16 architecture (ESK90106.1) and the linker sequence between the domains does not resemble an amyloid-like motif.

**Figure 4:**
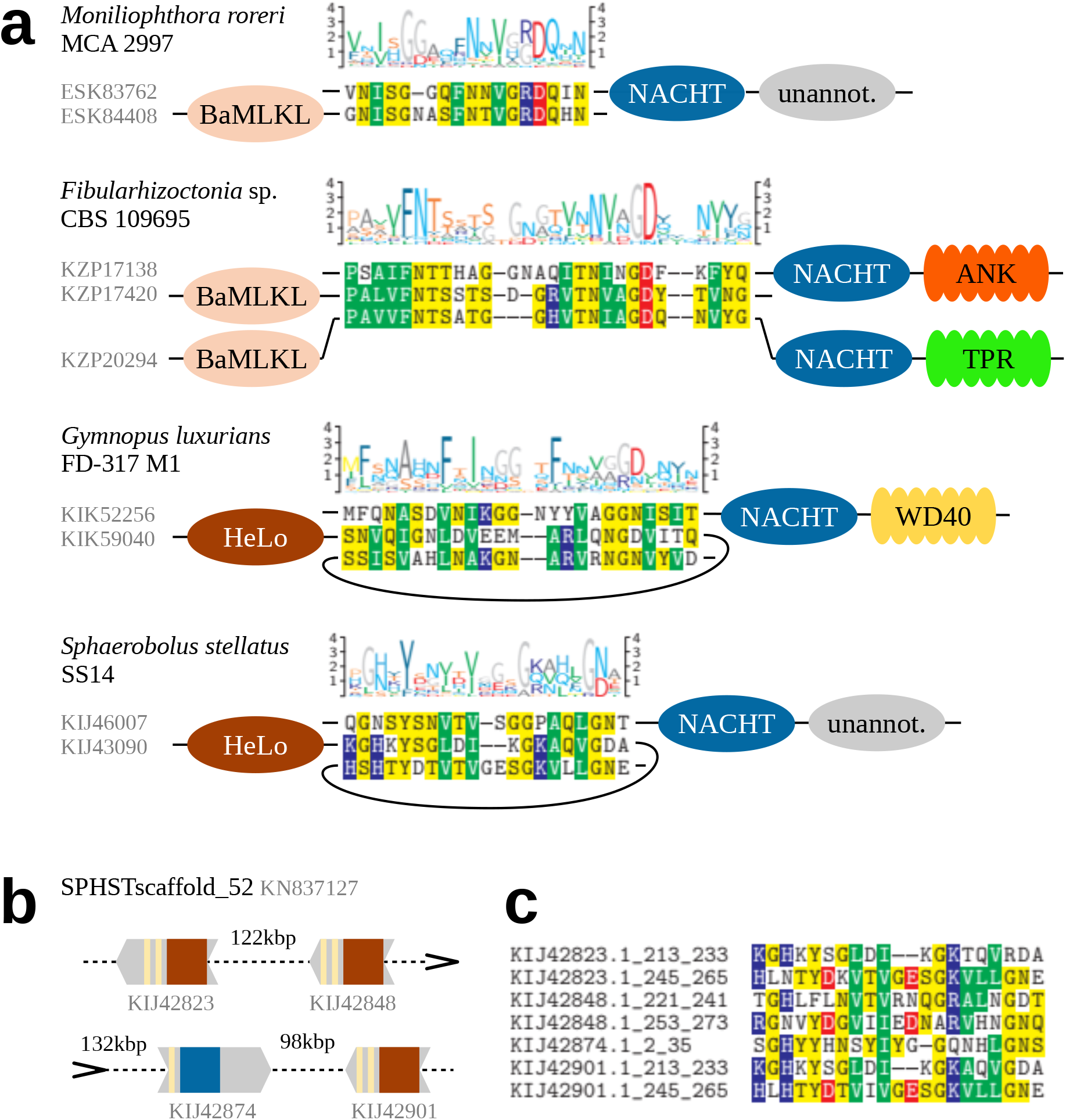
Potentially interacting amyloid-like motifs in Agaricomycetes. (**a**) Motif logos, sequence alignments and domain architectures of selected motif instances. (**b**) Schematic representation of a cluster of amyloid-like motifs in contig SPHSTscaffold_52 from genome assembly GCA_000827215.1 of *Sphaerobolus stellatus* SS14. (**c**) Multiple sequence alignment of motif instances in (b).

In addition, we investigated two Agaricomycetes species with proteins comprising of a singular HeLo domain and a C-terminal double HET-s motif. In the genome of *Sphaerobolus stellatus* (strain SS14), which includes four such C-termini, we found at least eight NACHT NLRs with N-termini comprising of single HRAM-like motifs (Supp. Fig. S6). Two instances (in KIJ28522 and KIJ30800) are recognized as the NLR13 HRAM motif at E-value around 0.01. This strain is the only case where an NLR and three HeLo proteins are situated on a single contig in genome assembly (Fig. 4bc). Interestingly the shortest distance between genes encoding NLR and HeLo is 95 kbp. The second species, *Gymnopus luxurians* (strain FD-317 M1), includes one protein with HeLo + double HET-s motif architecture. While we did not find any typical HRAMs in N-termini of 200 NLRs, several dozens included an instance of the NLR05/08/22/44 motif meta-family. Eventually we extracted N-terminal motif with highest scores according to the PCFG model and aligned them with Mafft, which revealed a 25-residue long core pattern. Interestingly, the alignment exhibited features characteristic to HRAMs: the N-terminal pattern of three hydrophobic residues and the C-terminal G[DN] bigram (Supp. Fig. S7). The 32 motifs are associated with NB-ARC, NACHT, NACHT WD and NACHT TPR domain architectures. Taken together these analyses strongly suggest that the NLR-associated amyloid signaling process also occurs in Basidiomycota.

### Experimental validation of a novel amyloid signaling motif

The alignment of the PUASM motif pairs (Fig. 5a) reveals high similarity of PNP_UDP- and NLR-side sequences in the core region covered with the NANBNtm_035 pattern. Some divergence can be seen C-terminally, with GND pattern prevailing in PNP_UDP-side motifs, while ARD pattern — in NLR-side motifs. Interestingly, these 3-mers can be found in C-termini of already known amyloid signaling motifs HRAM1 [33] and BASS2 [20], respectively. Further four residues of the C-terminal extension of the motif exhibit a hydrophobic pattern well conserved in pairwise alignments (Fig. 5a). On the other side, N-terminal extensions of the PUASM profile matches often include histidine in the PNP_UDP side and glutamic acid in the NLR-side. This, together with the overall composition of the N-terminal extensions, suggests some role of the charge complementarity.

**Figure 5:**
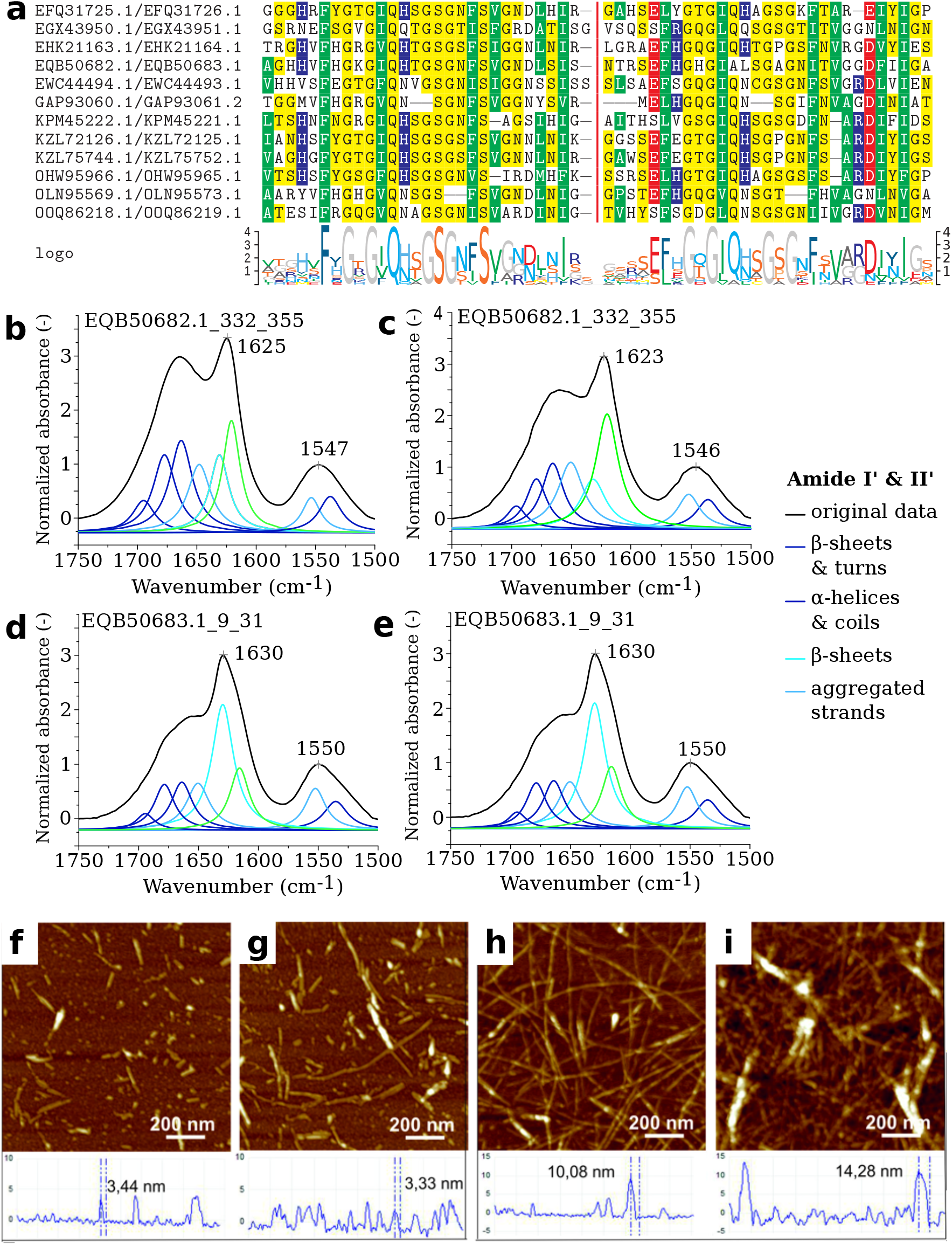
The PUASM motif. Panel **a:** Alignment of the PUASM motif pairs (left: C-terminal, right: N-terminal). Colors indicate residue hydrophobicity, curly brackets — the motif ranges. Panels **bcde:** Normalized ATR-FTIR spectra of air-dried peptide films of EQB50682.1_332_355 (**bc**) and EQB50683.1_9_31 (**de**) with sub-bands obtained from the curve fitting procedure in the amide bands region (1750–1500 cm^−1^) registered at 20°C. Panels **fghi:** AFM images with cross-section profiles of peptides EQB50682.1_332_355 (**fh**) and EQB50683.1_9_31 (**gi**). Samples were measured and imaged after dissolving (**bdfg**) and after 40 days of incubation at 37°C (98.6°F) (**cehi**).

To check if biochemical properties of the PUASM motif are consistent with its presumed role as the amyloid signaling motif, we experimentally analyzed a representative pair of motifs of this family, namely, PNP_UDP-side C-terminal EQB50682.1_332_355 (VFHGKGIQHTGSGNFSVGNDLSIS) and NLR-side N-terminal EQB50683.1_9_31 (FHGHGIALSGAGNITVGGDFIIG) from a plant pathogenic fungus *Colletotrichum gloeosporioides* Cg-14 [89] (Tab. S1 and Fig. S8). The selected fragments entirely cover the matches of PUASM profiles and the pairwisely conserved C-terminal extensions.

#### *In vitro* study of amyloidogenic properties

The aggregation propensities of the PUASM peptides were determined experimentally using the Attenuated Total Reflectance – Fourier Transform InfraRed spectroscopy (ATR-FTIR), Atomic Force Microscopy (AFM), and the Thioflavin T fluorescence assay (ThT), in accordance to the MIRRAGGE standard [90]. The ATR-FTIR spectroscopy allows determination of secondary structure and monitoring structural changes of peptides upon aggregation processes [91, 92, 93], while AFM is useful for detection and visualization of aggregates [94]. In turn, ThT is considered to be a “gold standard” for identifying amyloid fibrils [95, 96]. It is widely accepted that a combination of these techniques is necessary to ascertain if a particular peptide or protein is able to form the amyloid assemblies [97, 98, 90].

Analysis of the ATR-FTIR spectra in the range of 1750–1500 cm^−1^ (Fig. 5bcde, S9 and Tab. S2, S3) confirmed aggregation properties of studied peptides. The maximum of Amide I’ band was observed at 1625 cm^−1^ and 1630 cm^−1^ for EQB50682.1_332_355 and EQB50683.1_9_31, respectively. This signature is considered to be a spectroscopic marker of the formation of intermolecular aggregates [99] and corresponds to the cross-*β* amyloid architecture [92]. High absorbances in region of 1670–1660 cm^−1^ are characteristic of parallel *β*-helix structure and commonly observed in infrared spectra of peptides and proteins with beta solenoid conformations, e.g. HET-s [100] or PrPSc [101]. Further analysis of the deconvoluted and derivative spectra revealed more detailed information about the structure of aggregates. While for both studied peptides the aggregation process was observed immediately after dissolving, N-terminal EQB50683.1_9_31 aggregated faster and formed more rigid assemblies (Fig. 5bd). The percentage area of the subband corresponding to intramolecular *β*-structure at about 1630 cm^−1^ was 31% and 14%, for peptide EQB50683.1_9_31 and EQB50682.1_332_355 respectively. Moreover, the low-frequency component of Amide I’ band, which is generally assigned to highly ordered intermolecular structures, in the spectra of peptide EQB50683.1_9_31 is observed at 1616 cm^−1^. In turns EQB50682.1_332_35 exhibits this signature at higher wavenumbers (1616 cm^−1^), indicating a looser fibrilar structure [102].

Atomic Force Microscopy (AFM) images of both PUASM peptides were acquired for two conditions related to the spectroscopy studies: after dissolving, and after 40 days of incubation in room temperature. The aggregation process of the peptides was present already in the sample after dissolving as the fibers with height of 3.44 *±* 0.3 nm and 3.33 *±* 0.3 nm, respectively for EQB50682.1_332_355 and EQB50683.1_9_31, were observed (Fig. 5fg). The height of the object observable in AFM is comparable with the size of the HET-s peptides obtained by the solid-state NMR technique (pdb:2kj3) [30]. Peptide aggregation was further enhanced in the samples imaged after 40 days (Fig. 5hi) when the height of the aggregates reached 10.08 *±* 0.9 nm and 14.28 *±* 1.3 nm, respectively for EQB50682.1_332_355 and EQB50683.1_9_31. Along with the increasing aggregation process, the fibers underwent morphological changes. In the case of peptide EQB50682.1_332_355, greater flexibility of the fibers was observed, which may be related to the looser packing of the fiber structure. On the other hand, in the case of peptide EQB50683.1_9_31, a greater ability to form aggregates and greater stiffness of the fibers were observed, which may be related to tighter packing of the fiber structure. These observations are in line with the ATR-FTIR measurements (Fig. 5ce).

Thioflavin T (ThT) fluorescence assay was used to assess the kinetics of aggregation process. We observed a typical sigmoidal nucleation–polymerization curve for peptide C-terminal EQB50682.1_332_355, starting with a lag phase of 2 hours (Fig. S10), followed by a rapid growth phase from 2–2.20 h, and ending at a stable plateau with the maximum ThT intensity. A significant increase in the fluorescence emission was observed for peptide N-terminal EQB50683.1_9_31 (about 5 times higher than for peptide EQB50682.1_332_355). What is more, the lag phase was not observed, the curve starts from the elongation phase, which corresponds to the oligomers and protofibrils states. The steep ThT curve with quicker attainment of plateau indicate faster aggregation process of peptide EQB50683.1_9_31 in comparison to peptide EQB50682.1_332_355.

#### *In vivo* study of signaling capability

It was previously reported that fungal, bacterial and mammalian amyloid motifs could form prions *in vivo* in the *Podospora anserina* model [27, 20, 103, 36]. To determine if PUASMs could also form prions in vivo, we expressed the PNP_UDP-side C-terminal EQB50682.1_332_355 (VFHGKGIQHTGSGNFSVGNDLSIS) from a plant pathogenic fungus *Colletotrichum gloeosporioides* Cg-14 [89] in *P. anserina*. Prion propagation and transmission was monitored using fluorescence microscopy as the formation of cytoplasmic fluorescent foci. The GFP-PUASM fusion lead to an initial diffused fluorescence signal (Fig. 6). Upon subculture, the number of transformants showing cytoplasmic foci gradually increased over time as typically observed for other prion amyloid motifs [103]. The rate of this spontaneous transition from the diffuse state to the aggregated state was monitored over 75 days (Tab. S4). In contrast to the GFP-PUASM construct, fusion constructs displaying the motif N-terminally (PUASM-GFP and PUASM-RFP) remained diffused and did not form foci. A similar situation was observed previously for the HELLF and RHIM motifs for which N-terminal position of the GFP/RFP inhibited foci formation [103].

**Figure 6:**
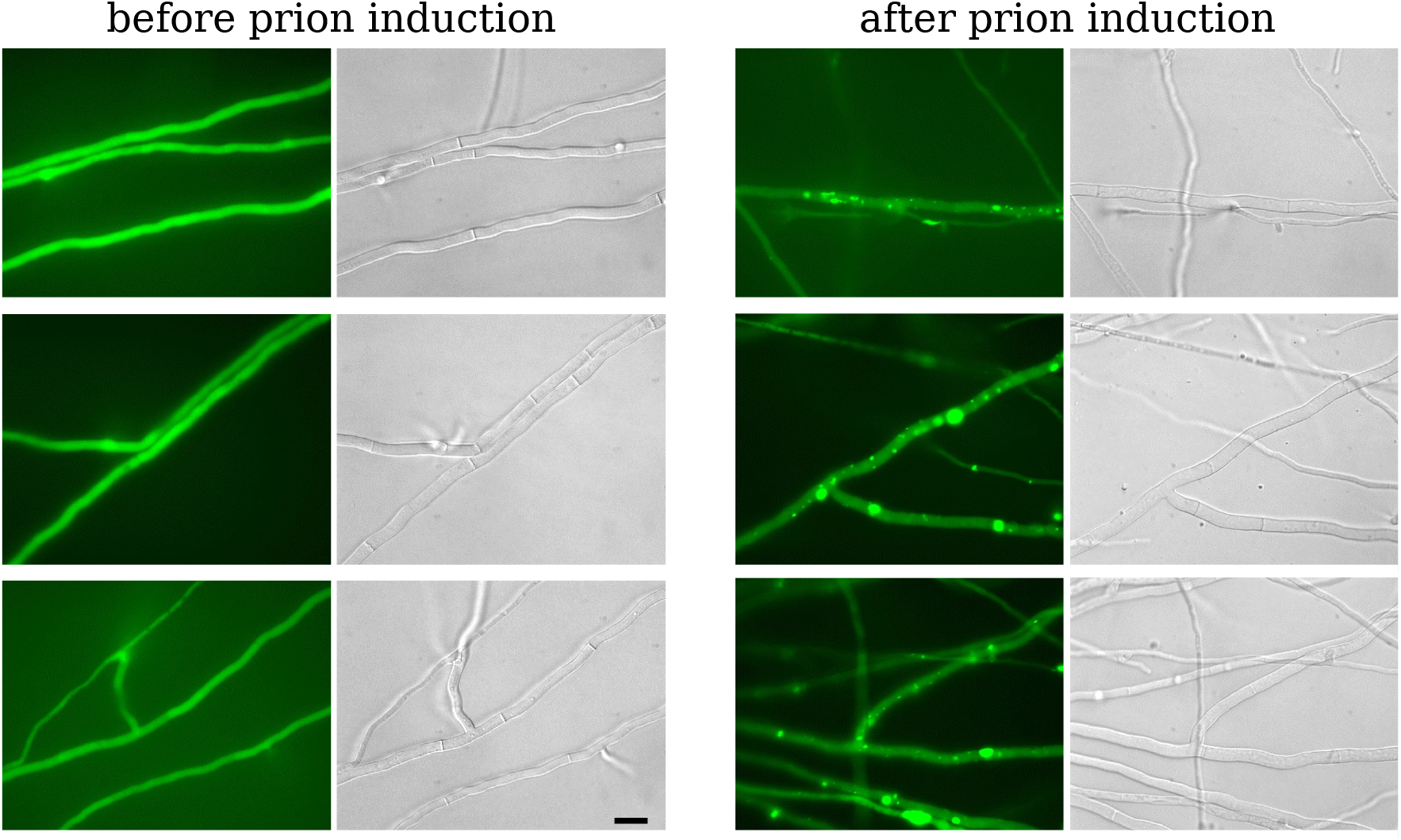
Expression of GFP-fused PUASM motif in *Podospora anserina*. Micrographs of *P. anserina* strains expressing molecular fusions of PUASM motif with GFP (in N-terminus) (Scale bar 5 *µ*m). Transformants displayed an initial diffuse fluorescence phenotype (**left** side of the panels) and acquired dot-like fluorescent aggregates, prion phenotype, after contact with strains already spontaneously expressing the infectious aggregated prion state (**right** side of the panels).

For all the amyloid signaling motifs tested so far in *P. anserina*, the aggregated state could propagate as a prion by cytoplasmic contact with a strain already expressing the fusion protein in the aggregated state. This assay, referred to as induced prion formation, was also performed for the PUASM fusion proteins. The GFP-PUASM fusion was converted to the foci state with high efficiency after cytoplasmic contact with a donor strain containing foci (Tab. S4, Fig. 6). Again, for the PUASM-GFP and PUASM-RFP proteins prion conversion was not observed. We conclude from these experiments that the GFP-PUASM fusion protein behaves as a prion *in vivo* in the *Podospora* model. The spontaneous and induced prion conversion was however less efficient than for other amyloid motifs [20, 103, 36].

## Discussion

### N-terminal annotations of fungal NLRs are not evenly distributed

In previous studies we computationally screened N-terminal domains of fungal NLRs using profile Hidden Markov Models (pHMM) from the Pfam database directly and complemented the search with several Pfam-like inhouse models [14, 5]. Here we expanded the most recent analysis with a more sensitive search using the state-of-the-art clustering offered by MMseqs2 and pHMM–pHMM searches with HHblits. The study increased the overall Pfam annotation coverage of N-terminal domains by about 16% (or 19% when MLKL_NTD from CDD is counted), but also highlighted remarkable deficiencies in availability of annotations.

While the vast majority of longer domains is at least partially annotated, this is true only for a definite minority of shorter domains (Fig. 1c). As about 3 400 short domains were assigned to clusters made with at least 20 sequences, the shortage of annotations cannot be easily explained by the lack of conserved sequential features. Instead, one of the reasons is the pHMM model itself, which by assessing each alignment position independently (except for indels) is not statistically powerful enough when dealing with short sequences. In other words, pHMM models of more diverse families of short sequence fragments (e.g. 20–40 amino-acids long) cannot be sensitive and specific at the same time [43]. Currently, the problem can be at least partially addressed by using more complex and computationally demanding protein sequence models, such as probabilistic context-free grammars (PCFG) [104, 20, 43] and co-evolutionary Potts models [105, 106, 107]. Another viable option are the recurrent and attention-based neural networks, which have enough computational power to describe relevant dependencies in protein sequences [108, 109, 110]. However, while modern neural networks have been successfully applied to annotation of protein families [111, 112], their performance in modeling short protein sequence fragments is yet too be evaluated.

### Differences between the Asco- and Basidiomycota effectors

The Pfam annotation coverage strongly depends on taxonomic scope. Specifically, the Basidiomycota are covered three times less than Ascomycota (20-25% vs. 70%, Fig. 1d), with other fungi in between (about 40%). Moreover, the disproportion in a share of annotated NLR effectors in asco- and basidiomycetes holds also when comparing N-terminal extension in the same length ranges. In addition, domains with homologs in other Eukaryota were significantly better annotated than domains specific to fungi (80% vs. 20%, Fig. 1e). While better coverage of more universally spread domains is not surprising, taken together, our results highlight the fact that the NLRs of fungi, and especially Basidiomycota, are still not sufficiently described. Certainly, this may have implications for understanding the NLR system in general, e.g. regarding its evolutionary history [15, 26].

The previous and present annotation reveals some resemblances between the effector domains found in ascomycetes and basidiomycetes, the most notable being the high proportion of domains in the Goodbye/HeLo/MLKL group (Fig. S1). Yet, some marked differences also come to attention, like the lack of occurrence of HET or PUP domain basiomycetes (although these domains occur in other (non-NLR) domain architectures). Another difference is the length distribution of the effector domains that globally appear much shorted in Basidiomycota. What are the underlying causes of these marked differences in the evolutionary history of the NLR-family in these two major fungal groups remains at present elusive.

### MLKL-related HeLo- and Goodbye-likes form the most abundant superfamily of effectors

Analyzing the annotation results, we found clusters with overlapping HeLo/HeLo-like and HeLo-like/Goodbye-like hits assigned through the HHblits search. The HeLo domain is a fungal homolog of human MLKL, plant RPW8 and bacterial Bell domains, while HeLo-like is closely related [32, 40, 20]. It is understood that upon oligomerization, these domains, whose central part is a four-helix bundle, expel a N-terminal alpha-helix to form a pore targeting the membrane and thus induce the cell death. The structure and function of Goodbye-like is yet to be established experimentally. The Helo-like/Goodbye-like cases, typical to Basidiomycota, were usually below the default threshold for direct Pfam annotation but were often assigned the MLKL_NTD annotation of the Conserved Domain Database. The Goodbye/HeLo/MLKL-like superfamily comprises of approximately 30% of all N-termini of fungal NLRs (Tab. 1) and is the most numerous both in Asco- and Basidiomycota.

Both the distribution of associated nucleotide-binding domain and C-terminal domains, and the paralog-to-ortholog ratio for Goodbye-like and HeLo-like domains are similar [14], which may suggest similarities in their mode of operation. However, Goodbye-likes in NLR N-termini are often associated with another annotated effector domains, which is untypical for HeLo-likes (Tab. 1). Moreover, the opposite is true for association with the amyloid signaling motifs, which is common to HeLos, HeLo-likes and basidiomycotal MLKL_NTDs but not to Goodbye-likes. The multiple sequence alignment and predicted structures of Goodbye/HeLo/MLKL-like domains reveal the matching region covers the four-helix bundle (Fig. 2). In a plant homolog of HeLo, the N-terminal helix of the bundle (and entire protein) is known to play a significant role in triggering the cell death process [72]. However, in BaMLKL and Goodbye-likes, the bundle is extended N-terminally by one or more helices. Thus, while the common evolutionary ancestry of HeLo-like and Goodbye-like seems rather evident, the question of their functional similarity remains open.

### Effector families distribution: the widespread and the confined

In addition to the Goodbye/HeLo/MLKL-likes, the other two superfamilies highly abundant in Ascomycota (a dozen of percent each, Fig. 1e and Fig. S1) are SesB-like AB hydrolases and the purine and uridine phosphorylases, which both provides enzymatic functions. Apart from them, the only other more frequent longer domains are Patatin (2-3% of Asco- and Basidiomycota) and HET (ca. 3% of Ascomycota). Interestingly, a couple of dozens of domains from the TIR (Toll/Interleukin-1 receptor-like) clan, remotely homologous to HET, where found in Pezizomycotina and in Chytridiomycota. All other longer domains — whether determined by the Pfam annotation or the MMseqs2 clustering — were represented by less than 100 sequences. This indicates that the current Pfam annotations (plus MLKL_NTD and a few inhouse profiles) cover all widely spread abundant domains. At the same time, there seems to exist a substantially large corpus of thousands of specialized N-termini, sometimes confined to narrow taxonomic branches. While some of them may be formed with a tuple of known domains, other could represent novel families (likely being difficult targets for structure prediction due to small alignments). Naturally, we cannot rule out that in some cases larger families were superficially partitioned into small clusters (with less than 20 members). Nevertheless, with regard to our previous analyses [14, 5], this study suggests less diversity in major effector classes (5-7 rather than 12-13), but highlights a likely abundance of specialized domains.

### A large fraction of the effector domain are tentatively involved in regulated cell death

With the limitation that the evolutionary relation of Goodbye-like domains does not necessarily imply functional similarity, it appears that a substantial fraction of the effector domains in both ascomycetes and basidiomycetes is predicted to control regulated cell death. Involvement in regulated cell death has been reported not only for the HeLo/Goodbye/MLKL group but also for the HET domain [113], the Patatin [18] domain and more indirectly for the SesB-like domain [40]. One needs to add to this list the amyloid signaling motifs that control separate downstream cell-death effector domains. Globally, it would appear of at least one third to one half of the fungal NLRs could be involved in some kind of regulated cell death process. This high proportion raises the question of whether some of the other domains (whether annotated or not) could also play a role in regulated cell death.

### NLR-associated amyloid-like motifs in Ascomycota are more diverse than previously reported

We significantly improved the annotation coverage of NLR amyloid signaling domains. By pipelining clustering with MMseqs2, filtering sequences with our recent PCFG-based generalized model of beta-arch amyloid motifs, extracting specialized motifs with MEME, and generating their profiles with HMMER, we identified amyloid-like motifs in roughly one third of the 3 400 short N-terminal domains (Fig. 1 and Tab. 3).

In Ascomycota, within several hundreds of extracted motif instances, we discovered a new motif family, NLR32, which is uniquely associated with the PNP_UDP domain (thus termed PUASM, Fig. 3). Similarly to other ascomycotal amyloid signaling motifs, HRAM, PP and *σ*, effector proteins with C-terminal PUASM are often coded by direct genomic neighbors of the PUASM–NLR genes. Such genomic co-localization facilitates co-inheritence of the two genes of the functional unit in the event of a recombination process. This may be of special importance for the NLR signaling pathway, which is polymorphic in population given the death-and-birth evolution. The amyloid properties of the NLR32 motif were confirmed experimentally using a representative pair of N- and C-terminal sequences. While both of them generated amyloid-like fibers, the N-terminal motif aggregated quicker and formed more rigid structures, which is consistent with its expected role as the template for aggregation of the C-terminal motif. (In depth study of the co-aggregation process is left for a separate study.) The effector-side PUASM sequence was shown to be capable of forming prions *in vivo* in the *Podospora anserina* model.

Interestingly, we found the PNP_UDP domain associated also with a HRAM motif variant (NLR12) and with one another amyloid-like motif discovered in this study (PF01048_40, Fig. S3). For NLR12, only in a few cases the effector–motif and motif–NLR pairs present in the genome were co-localized (that is encoded by adjacent genes). In some genomes, the NLR-side motifs were relatively more frequent (mean ratio 5.7:1, Tab. 3). We speculate that the presence of many NLRs controlling the same effector could potentially relieve the need for genomic co-localization.

### Less diversity of ASM in fungi as compared to bacteria even after extensive search

Although, we have successfully identified new motifs and validated the most abundant of them, the expand diversity of ASM remains higher in bacteria than in fungi using similar identification procedures, which is not inconsistent of the larger phylogenetic breath of the scanned bacterial genome as compared to the fungal ensemble.

### NLR-related amyloid signaling is present in Agaricomycetes

In Basidiomycota, virtually none of the hundreds of NLR-side amyloid-like motifs found in our survey was genomically co-localized with C-terminal side motifs. Moreover, motifs closely resembling NLR-side amyloid-like motifs were absent in C-termini of previously identified effector proteins whatever their location in genome. Eventually, by analyzing C-termini of single-domain MLKL-like proteins we identified likely associations in *Moniliophthora roreri* and *Fibularhizoctonia* sp. CBS 109695. Similarly by analyzing two genomes with HeLo-HRAM-HRAM architectures, we found HRAM-NLR proteins in *Sphaerobolus stellatus* and NLR05/NLR22-NLR proteins in *Gymnopus luxurians*. As NLR05/NLR22 meta-family of amyloid-like motifs shares with HRAM the hydrophobic pattern and the largely conserved C-terminal G, the study suggests that the amyloid signaling could be preserved despite substantial sequence variability. Eventually, limits of this preservation requires experimental verification [36, 114].

Taken together, presented results support the presence of amyloid signaling in Basidiomycota, or more specifically in Agaricomycetes, in the context of NLR-based regulation of HeLo-/MLKL_NTD-likes. Moreover, they suggest that NLR05/08/22/29/44 meta-family of motifs is a basidiomycotal variety of the HRAM motif or its homolog. However, there are significant differences with regard to Ascomycota. First, while NLR-side amyloid signaling motifs are present in roughly half of Ascomycota strains, they were only found in 1*/*4 (30%) of Basidiomycota (Agaricomycetes) strains. Second, while there are typically only few such motifs per ascomycotal strain, there are usually dozens per basidiomycotal strain. At the same time, basidiomycotal effector-side C-terminal motifs are seemingly less frequent than NLR-side N-terminal motifs (Fig. S4-S7). Indeed, the high number of NLR-side motifs corresponds to enrichment of basidiomycotal sequences among shorter N-terminal domains (Fig. 1b). This could at least partially explain the lack of genomic co-localization of NLRs and their effectors linked by amyloid signaling motifs.

One interesting finding is presence NLRs with intra-proteins amyloid-like motifs in *Fibularhizoctonia* sp. CBS 109695. Different central and C-terminal domain association in comparison to NLRs with N-terminal ASM-likes suggest also different functions of the motifs in both cases. Therefore, we hypothesize that these internal amyloid motifs may serve as scaffolds to stabilize the NLR oligomers, similar to cRHIM motifs in the RIP1K/RIP3K complex [45]. In these lines, it is possible that also some other amyloid-like motifs identified in the current study but with no matching effector-side counterparts participate in the assembly of the NLR signalosome or are involved in interactions with motifs located outside the C-terminus of the associated protein.

## Materials and Methods

### Computational methods

#### Annotation of NLR N-termini

A set of 36,141 NLR proteins from 487 fungal strains was identified in a previous study through the PSI-BLAST [115] search among completely sequenced fungal genomes in the NCBI nr database [14, 20]. 32 962 N-termini at least 20 amino-acids long (91%), delimited according to the NACHT or NB-ARC query matches, were further considered, of which 18,674 (57%) were annotated using direct matches to Pfam [25] or inhouse HMM profiles (Supplementary File 2) [14, 20]. The set of N-termini at least 20 amino-acid long was clustered with MM-seqs2 [46] in mode 1 (21 758 N-termini in 127 clusters, 15 105 already annotated). Then, sequences in each cluster with at least 20 members were aligned using Clustal-Omega [116] (Supplementary File 3) and searched for homologs in UniRef30 [47, 48] using HHblits [49] (parameters: -e 0.001 -n 2 -E 0.01 -Z 1000000 -M 50). Subsequently, the resulting alignment was used to search Pfam (HHblits parameters: -e 0.001 -n 1 -E 1 -Z 1000000). The clustering required mutual coverage of at least 80% of sequence length, and the annotations were only assigned to sequences which covered at least 50% of the match to the Pfam profile. The resulting cluster-level annotations were retained only if the alignment match to the Pfam profile covered at least 50% of the profile length, and assigned only to individual sequences which covered at least 50% of the match. After completing the main processing, the set of N-termini was re-scanned for the Crinkler domain (PF20147) added recently to the Pfam database.

The tabularized results of the annotation are provided in Supplementary File 4. The overlapping Pfam annotations were resolved as in [14, 20]. The double HeLo/HeLo-like annotations were kept in the Supplementary File 4 and in Tab. 1 but were represented as HeLo in Fig. 1f and Fig. S1. In addition, basidiomycotal sequences from clusters doubly annotated as Goodbye-like/Helo-like, as well as from clusters with CDD [65] MLKL_NTD annotations, were denoted as BaMLKL (see Results).

#### Comparative analysis of Goodbye-, HeLo- and MLKL-likes\

The largest clusters annotated as HeLo, HeLo-like, Goodbye-like and BaMLKL were submitted to AlphaFold2 structure prediction [69] through the ColabFold advanced notebook [70]. Standard parameters of the notebook were applied except of (1) using the cluster alignments instead of searching genetic databases, (2) trimming off fragments just upstream the NACHT domain were applicable. Successful models – with the mean predicted lDDT score [69] above 70 overall, and around 80 or more for the core helix bundle – and respective ColabFold outputs are provided in Supplementary File 5. For each cluster, the highest rank model was selected and structurally aligned to the experimentally solved MLKL domain (pdb:6zvo) using TM-align [71].

#### Characterization of unannotated longer N-termini

For five unannotated clusters with at least ten members at the identity threshold of 70% and the median length above 100 amino acids homologs were searched in UniProt [117] through the web-based hmmsearch with standard parameters [118], and prediction of the three dimensional structure was attempted using AlphaFold2 [69] through the ColabFold advanced notebook [70]. Standard parameters were used except of adding the MMseqs2 alignments to input (sequences just upstream the NACHT domain was trimmed off). Good quality structures (the predicted lDDT score above 70) were obtained for three clusters, KEY84097, KFH66451 and PQE30996 (Supplementary File 6). The proposed annotations for the five clusters (Tab. 2) are assigned to member sequences in the table in Supplementary File 4 and included in the TIR-like and “other” groups in Fig. 1f and Fig. S1.

#### Extraction of amyloid-like motifs in short N-termini

A subset of 54 NLR N-termini clusters with mean/medium sequence length at most 160/161 amino acids was scanned using the PCFG-CM software [119, 43] probabilistic grammatical model inferred from ten BASS families [43] (Supplementary File 7) with scanning window of 20 to 40 amino acids and the smoothing factor of 10 PAM [43]. Very high scoring fragments (maximum log10 score at least 3.5, mean log10 score above 1.67) were found in 18 clusters with 1456 sequences. The N-terminal sequences were made non-redundant at the identity level of 90% using CD-HIT 4.7 [87, 88] and submitted to motif extraction with MEME 5.0.5 [120, 82]. For each of 51 motifs found at the E-value threshold of 1, HMM profiles were built with HHMER 3.2.1 [86] and used for searching against the full set of grammar-fitting N-terminals (at E-value of 1*e −* 2). Then, obtained hits were extended by 5 amino acids in each direction and realigned using Clustal-Omega with the auto parameter. For each motif, the extended sequences were re-examined for consistency with the grammatical model (maximum log10 score at least 3, mean log10 score above 1). For 16 motifs which passed the grammatical filter, the alignments were used to build final HMM profiles (Supplementary File 8).

#### Analysis of N-terminal amyloid-like motifs

The HMM profiles of the 16 motifs were used for scanning all N-termini longer than 10 amino acids (the independent domain E-value threshold of 1*e −* 2). Hits are included in the Supplementary Tab. 4 with coordinates (outermost in rare cases of double ASM hits). For further analysis only hits in N-termini shorter than 200 amino acids not located beyond position 150 were considered. Motif sequences in envelopes of 5 amino acids were tested for the beta-arch structure with ArchCandy 2.0 [42] using the recommended threshold of 0.56.

For each motif-containing NLR sequence, proteins coded by genes within the *±*20kbp neighborhood of the genes encoding these NLRs were fetched from NCBI GenBank [121] or EMBL ENA [122] using an in-house Python (version 3.7.3) script (aided by requests [123] and xmltodict [124] packages). The set was confined to proteins in the length range of 200-400 amino acids, which is typical for proteins with single domain architectures known to be associated to NLRs via amyloid signaling [33, 20]. Next, C-termini (100 amino acids) of the found neighboring proteins were scanned for the presence of the motifs using HMMER (the independent domain E-value threshold of 1*e −* 2). Pairwise hits of the same motifs in N-termini of NLRs and C-termini of genomically neighboring proteins are collected in Supplementary File 9.

#### Homology search of effector domains

Homologs of NLR effector domains, as listed in [5], were iteratively searched using Pfam profiles: Pkinase (PF00069), Peptidase_S8 (PF00082), C2 (PF00168), PNP_UDP_1 (PF01048), TIR (PF01582), Patatin (PF01734), RelA_SpoT (PF04607), DUF676 (PF05057), HET (PF06985), PK_Tyr_Ser-Thr (PF07714), PGAP1 (PF07819), Abhydrolase_6 (PF12697), CHAT (PF12770), TIR_2 (PF13676), HeLo (PF14479), NACHT_N (PF17100), SesA (PF17107), Goodbye (PF17109) and Helo_like_N (PF17111). First, HMM profiles for each of the domains were used for searching against a local copy of the non-redundant protein sequences database (NCBI’s “nr”, downloaded in November 2019) [121] using HHMER 3.2.1 with E-value of 1*e−* 2. Found proteins were then used to build the new HMM profiles and the search was repeated until the number of hits did not change by more than 7%. In addition, all fungal NACHT (PF05729) and NB-ARC (PF00931) proteins (as listed in the Pfam database in January 2020) were retrieved. C-termini of effector domains and N-termini of NACHT/NB-ARC NLRs were extracted and – in both cases – only fragments between 10 and 150 aa were selected for further analysis. (This effectively excluded proteins with effector + NOD architectures.) The final set included around 200k of effector C-termini and 6.8k NLR N-termini (redundant).

#### Identification of paired amyloid motifs

Sets of N- and C-terminals were clustered using CD-HIT to reduce redundancy at the 70% similarity threshold (separately for each effector domain, together for NACHT and NB-ARC). Motif search was performed using MEME with the following parameters: -nmotifs 100, -minsites 1% of sequences but no less than 5 and no more than 10, -maxsites 500, -minw 10, -maxw 30, -mod anr. For each of the identified motifs, HMM profiles were built in a two-stage procedure as described above with an exception that initial profiles were used for searching against all N- and C-termini. The same-motif hits in effector proteins and NLRs were matched based on genomic proximity (up to 20 kbp) of genes encoding the proteins. Motifs with at least 3 non-redundant pairs of motif instances (Supplementary File 10) were then clustered on the basis of their co-occurrence in pairs of sequences.

Finally, ASM hits in N-termini of all NLRs investigated in this study and in C-termini of all effector domains found were matched on the strain level (through the BioSample and BioProject identifiers) in order to identify potentially correlated pairs, which are not co-localized in genomes (Supplementary File 11).

#### Specialized searches for amyloid motifs in Basidiomycota

MLKL_NTD homologs were searched in UniProt [117] through the web-based hmmsearch [118] with standard parameters starting from the alignment of the largest BaMLKL cluster in Basidiomycota (representative protein: KIM77258), trimmed to the NACHT_N match. Hits were further restricted to GenBank sequences with length up to 400 amino acids and no Pfam P-loop_NTpase clan (CL0023) annotation at E-value of 1. C-termini (100aa) of resulting 242 BaMLKL homologs were scanned with the PCFG BASS model (Supplementary File 7) with the same parameters as above (except the minimum scanning window length of 15). For proteomes with the most promising hits in BaMLKL homologs (log10 score above 3, eight sequences from six species), N-termini (150 aa) of all NLR proteins were again scanned with the grammars. Promising N-terminal hits were obtained for *Moniliophthora roreri* (strains 2995 and 2997), *Laccaria amethystina* (strain LaAM-08-1), and *Fibularhizoctonia* sp. CBS 109695. The matched fragments were aligned with their C-terminal counterparts on the per genome basis with Mafft [125] in an accurate mode (–maxiterate 1000 –localpair). The NLR N-terminal and BaMLKL C-terminal motifs aligned satisfactorily for *M. roreri* (we only analyzed strain 2997 due to high similarity between the strains) and *Fibularhizoctonia* sp. CBS 109695. The alignments were then extended and trimmed manually (Fig. S4 and S5). In addition, the sequences were scanned with the 16 HMM profiles of amyloid-like motifs (E-value of 1*e−* 2).

Eventually, fungal proteomes in UniProt where scanned using web-based jackhmmer [118] with standard parameters starting from the double HET-s motif from Q03689 (AAB94631) (residues 218–289) of *Podospora anserina*, which resulted in finding five complete HeLo-HRAM-HRAM proteins in two Agaricomycetes: *Sphaerobolus stellatus* SS14 and *Gymnopus luxurians* FD-317 M1. NLRs in these genomes were then scanned with the PCFG model and the hits exceeding the log10 score threshold of 2.33 were aligned with their C-terminal counterparts on the per genome basis with Mafft [125] in the accurate mode. Finally, the alignments were curated manually (poorly aligned sequences were excluded, sequences were extended or trimmed if necessary, Fig. S6 and S7).

#### Visualization

Basic data processing and visualization was conducted in Python using pandas [126, 127], matplotlib [128] and seaborn [129] packages, as well as in LibreOffice, GIMP and Inkscape. Multiple sequence alignments and logos were generated using TeXshade [130]. The graph of logos in Fig. S3 was generated with graphviz 2.40.1 [131]. Visualizations of structural models were generated with RasMol [132] (Fig. 2) or taken directly from the ColabFold notebook [70] (Fig. S2).

## Experimental methods

### In vitro analysis

#### Peptide synthesis

All commercially available reagents and solvents were purchased from Merck, Sigma-Aldrich and Lipopharm.pl, and used without further purification. Peptides EQB50682.1_332_355 (VFHGKGIQHTGSGNFSVGNDLSIS) and EQB50683.1_9_31 (FHGHGIALSGAGNITVGGDFIIG) were synthesized with an automated solid-phase peptide synthesizer (Liberty Blue, CEM) using rink amide AM resin (loading: 0.59 mmol/g). Fmoc deprotection was achieved using 20% piperidine in DMF for 1 min at 90°C. A double-coupling procedure was performed with 0.5 M solution of DIC and 0.25 M solution of OXYMA (1:1) in DMF for 4 min at 90°C. Cleavage of the peptides from the resin was accomplished with the mixture of TFA/TIS/H_2_O (95:2.5:2.5) after 3 h of shaking.The crude peptide was precipitated with ice-cold Et_2_O and centrifuged (8000 rpm, 15 min, 2°C). Peptides were purified using preparative HPLC (Knauer Prep) with a C18 column (Thermo Scientific, Hypersil Gold 12 *µ*l, 250 *×* 20 mm) with water/acetonitrile (0.05% TFA) eluent system.

#### Peptide analytics

Analytical high-performance liquid chromatography (HPLC) was performed using Kinetex 5*µ* EVO C18 100A 150 *×* 4.6 mm column. Program (eluent A: 0.05% TFA in H_2_O, eluent B: 0.05% TFA in acetonitrile, flow 0.5 mL/min): A: t=0 min, 90% A; t=45 min (25 min in case of EQB50682.1_332_355). Peptides were studied by WATERS LCT Premier XE System consisting of high resolution mass spectrometer (MS) with a time of flight (TOF).

#### Attenuated Total Reflectance – Fourier Transform Infrared Spectroscopy (ATR-FTIR)

Lyophilized peptides were dissolved in D_2_O (deuterium oxide, 99.8% D, Carl Roth, GmbH, Germany) to a final concentration of 2 mg/ml. The spectroscopic measurements were performed directly after dissolving peptides in a solvent, after 7 and 40 days of incubation process at 37°C (98.6°F), and after 40 days of incubation at 4°C (39.2°F). Each time, 10 *µ*l of peptide solution was dropped directly on the diamond surface and was allowed to dry out. ATR-FTIR spectra were recorded using a Nicolet 6700 FTIR Spectrometer (Thermo Scientific, USA) with Golden Gate Mk II ATR Accessory with Heated Diamond Top-plate (PIKE Technologies). The spectrometer was continuously purged with dry air. Directly before sampling, the background spectrum of diamond/air was collected as a reference. For each spectrum 512 scans with a resolution of 4 cm^−1^ were co-added. All spectra were obtained in the range of 4000–450 cm^−1^ at 20°C (68.0°F).

#### Spectroscopy data treatment

ATR-FTIR spectra were initially preprocessed using OMNICTM software (version 8, Thermo Fisher Scientific, USA): atmospheric and ATR correction. All spectra were analyzed using the OriginPro (version 2019, OriginLab Corporation, USA). The analysis included: baseline correction, smoothing using the Savitzky-Golay polynomial filter (polynomial order 2, a window size of 9 points) [133] and normalization to 1 for the Amide II’ band. Spectra in the amide bands region (1750–1500 cm^−1^) were deconvoluted into subcomponents using the Lorentz function based on second and fourth derivative spectra.

#### Atomic Force Microscopy

AFM images were acquired in tapping mode using a Nanoscope IIId scanning probe microscope with Extender Module (Bruker) in the dynamic modus. An active vibration isolation platform was applied. Olympus etched silicon cantilevers were used with a typical resonance frequency in the range of 100–200 kHz and a spring constant of 40 N/m. The set-point amplitude of the cantilever was maintained by the feedback circuitry at 80% of the free oscillation amplitude of the cantilever. The samples with peptides were placed on freshly cleaved ultra-clean mica (Nano and More) and incubated at room temperature for 30 s. The mica discs were then rinsed with ultra-clean purified 18.2 MΩ deionized water and dried using gentle nitrogen gas flow. All samples were measured at room temperature in air. Structural analysis and height measurements of acquired images were performed with Nanoscope v.6.13 software.

#### Thioflavin T (ThT) fluorescence assay

ThT powder was dissolved in MilliQ to final concentration 2 mM and filtered through 0.22 *µ*m syringe. ThT solution was dissolved in 50 mM Tris-HCl (pH= 7.4) to final concentration 10 *µ*M and filtered. The 90 *µ*L of ThT buffer was mixed with 10 *µ*L of peptide solution (concentration 400 *µ*M) in the 96-wells plate. Samples were measured on the SpectraMax*Q*R Gemini^TM^ XPS Microplate (Molecular Devices LLC). The measurements were conducted in RT. The excitation wavelength was set at 450 nm and the emission was recorded in the range from 470 to 500 nm. Each group of experiment contained three parallel samples and the data were averaged after measurements.

### In vivo analysis

#### Strains and plasmids

The *Podospora anserina* Δhellp (ΔPa_5_8070) Δhet-s (ΔPa_3_620) Δhellf (ΔPa_3_9900) strain [103] was used as recipient strain for the expression of molecular fusions of PUASM (PNP_UDP-side C-terminal EQB50682.1_332_355 VFHGKGIQHTGSGNFSVGNDLSIS) from the plant pathogenic fungus *Colletotrichum gloeosporioides* Cg-14 [89] and the GFP (green fluorescent protein) or RFP (red fluorescent protein). These fusions were expressed from plasmids based on the pGEM-T backbone (Promega) named pOP [40] and containing either the GFP or RFP encoding gene, or in a derivative of the pAN52.1 GFP vector [134], named pGB6-GFP and containing the GFP encoding gene. In both cases, the molecular fusions were under the control of the strong constitutive *P. anserina* gpd (glyceraldehyde-3-phosphate dehydrogenase) promoter. The Δhellp Δhet-s Δhellf strain was transformed as described [135] with a fusion construct along with a second vector carrying a ble phleomycin-resistance gene, pPaBle (using a 10:1 molar ratio). Phleomycin-resistant transformants were selected, grown for 30 h at 26°C and screened for the expression of the transgenes using fluorescence microscopy. PUASM was amplified with specific primers either 5’ ggcttaattaaATGGTCTTTCATGGCAAGGGCATCC 3’ and 5’ ggcagatcttgctccGGAGATGCTGAGATCG 3’ for cloning in pOP plasmids, or 5’ ggcgcggccgcGTCTTTCATGGCAAGGGCATC 3’ and 5’ ggcGGATC-CTTAGGAGATGCTGAGATCGTTGCC 3’ for cloning in the pGB6 plasmid (capital letters correspond to the PUASM sequence). The PCR products were cloned upstream of the GFP or RFP coding sequence in the pOP plasmids using PacI/BglII restriction enzymes to generate the pOPPUASM-GFP and pOPPUASM-RFP vectors in which in addition to the BglII site, a two amino acid linker (GA) was introduced between the sequences encoding PUASM and GFP or RFP and cloned downstream of the GFP using NotI/BamHI restriction enzymes to generate the pGB6-GFP-PUASM plasmid.

#### Microscopy

*P. anserina* hyphæwere inoculated on solid medium and cultivated for 24 to 48 h at 26°C. The medium was then cut out, placed on a glass slide and examined with a Leica DMRXA microscope equipped with a Micromax CCD (Princeton Instruments) controlled by the Metamorph 5.06 software (Roper Scientific). The microscope was fitted with a Leica PL APO 63X immersion lens.

#### Prion propagation

Methods for determination of prion formation and propagation were previously described [136, 20]. Prion formation and propagation can be observed using microscopy by monitoring the formation of fluorescent dots. Spontaneous prion formation is first monitored as the rate of spontaneously acquired prion phenotype (dot formation) in the initially prion-free subculture after 5, 11, 18, 32, 49 and 75 days of growth at 26°C on corn-meal agar using microscopy as described. Prion formation can also be measured as the ability to propagate prions from a donor strain (containing prion) to a prion-free strain (induced strain). In practice, prion-free strains are confronted on solid corn-meal agar medium for 2 to 5 days (contact between strains were observed after 24 to 36 hours of culture) before being subcultured and observed by fluorescence microscopy for the presence of dots, this test is referred as induced prion formation. At least 18 different transformants were used and the tests were realized in triplicates. It is to note that transformants were randomly tested for prion formation allowing various expression levels of the transgene (high levels of expression are usually associated with rapid spontaneous prion formation) except for the induced conversion test where transformants expressing moderate level of transgene were preferred to limit the rate of spontaneous transition within the timing of the experiment that could mask the prion induction.

## Author contributions

**JWW**: data curation, formal analysis, investigation, methodology, software, writing; **ET**: data curation, formal analysis, investigation, methodology, software, validation, writing; **MG-G**: investigation, methodology, resources, validation, visualization, writing; **VC**: investigation, methodology, resources, validation, visualization; **NS**: investigation, validation, visualization, writing; **MS**: investigation, validation, resources, writing; **MK**: investigation, resources, supervision, validation, visualization, writing; **SJS**: conceptualization, funding acquisition, investigation, methodology, resources, validation, writing; **WD**: conceptualization, data curation, formal analysis, investigation, methodology, project administration, software, supervision, validation, visualization, writing.

## Acknowledgments

This research has been partially funded by National Science Centre, Poland, grant no. 2019/35/B/NZ2/03997 (PI: Prof. Małgorzata Kotulska) (WD, MG-G), by National Centre for Research and Development, Poland (ncbr.gov.pl), project no. POWR.03.02.00-00-I003/16 (NS), and by Politechnika Wrocławska statutory funds, and supported by Wroclaw Centre for Networking and Supercomputing grant no. 98 and the E-SCIENCE.PL infrastructure. Work in the SJS lab was supported by an ANR grant (SFAS, R-17-CE11-0035).

