## Supplementary File 1 for "Exploring a diverse world of effector domains and amyloid signaling motifs in fungal NLR proteins"

Table S1: **Analytical data for synthesized PUASM peptides.** Notations: $M_{cal}$ — calculated mass of the peptide, $M_{MS}$ — found mass of the peptide using HRMS, $t_{ret}$ — retention time in analytical HPLC spectra (see Fig. S8)

| Id/range | Formula | $M_{cal}$ | $M_{MS}$ | $t_{ret}$ |
|---|---|---|---|---|
| EQB50682.1 | $C_{107}H_{165}N_{33}O_{34}$ | [(M+2H)/2] 1229.6191 | [(M+2H)/2] 1229.6208 | 16.224 |
| 332 – 355 | | [(M+3H)/3] 820.0820 | [(M+3H)/3] 820.0734 | |
| EQB50683.1 | $C_{100}H_{153}N_{29}O_{28}$ | [(M+2H)/2] 1105.5813 | [(M+2H)/2] 1105.5806 | 18.848 |
| 9 – 31 | | [(M+3H)/3] 737.3901 | [(M+3H)/3] 737.393 | |

*Contributed equally
†Present address: Koc University, School of Medicine, Istanbul, Turkey
‡Corresponding author

Table S2: **Main Amide I' and Amide II" band components and integrated intensities** obtained from the curve fitting procedure of ATR-FTIR spectra of air-dried PUASM peptide films with tentative secondary structures assignments. The results from experiments on the day of dissolving and incubation for 7 and 40 days at 37°C (98.6°F)

| EQB50682.1_332_355 | | | | | | | |
|---|---|---|---|---|---|---|---|
| after dissolving | | after 7 days | | after 40 days | | band assignment | |
| band pos. [cm$^{-1}$] | area [%] | pos. [cm$^{-1}$] | area [%] | pos. [cm$^{-1}$] | area [%] | Amide | corresp. structure |
| 1695 | 6 | 1691 | 6 | 1690 | 7 | I' | $\beta$-sheets & $\beta$-turns |
| 1677 | 16 | 1676 | 13 | 1673 | 17 | | $\beta$-sheets & $\beta$-turns |
| 1663 | 19 | 1662 | 18 | 1658 | 16 | | $\beta$-sheets & $\beta$-turns |
| 1648 | 14 | 1646 | 21 | 1642 | 16 | | $\alpha$-helices & coils |
| **1631** | **14** | **1630** | **16** | **1625** | **18** | | **$\beta$-sheets** |
| **1621** | **18** | **1619** | **21** | **1617** | **14** | | **aggregated strands** |
| 1554 | 6 | 1549 | 2 | 1550 | 6 | II' | $\alpha$-helices |
| 1538 | 7 | 1535 | 3 | 1533 | 6 | | $\beta$-sheets & turns |

| EQB50683.1_9_31 | | | | | | | |
|---|---|---|---|---|---|---|---|
| after dissolving | | after 7 days | | after 40 days | | band assignment | |
| band pos. [cm$^{-1}$] | area [%] | pos. [cm$^{-1}$] | area [%] | pos. [cm$^{-1}$] | area [%] | Amide | corresp. structure |
| 1694 | 3 | 1696 | 3 | 1693 | 5 | I' | $\beta$-sheets & $\beta$-turns |
| 1679 | 11 | 1679 | 7 | 1674 | 15 | | $\beta$-sheets & $\beta$-turns |
| 1664 | 12 | 1667 | 12 | 1659 | 13 | | $\beta$-sheets & $\beta$-turns |
| 1650 | 12 | 1651 | 23 | 1645 | 13 | | $\alpha$-helices & coils |
| **1630** | **31** | **1628** | **26** | **1626** | **37** | | **$\beta$-sheets** |
| **1616** | **14** | **1616** | **22** | **1611** | **11** | | **aggregated strands** |
| 1552 | 10 | 1552 | 4 | 1553 | 3 | II' | $\alpha$-helices |
| 1535 | 7 | 1537 | 3 | 1538 | 3 | | $\beta$-sheets& turns |

Table S3: **Main Amide I' and Amide II' band components and integrated intensities** obtained from the curve fitting procedure of ATR-FTIR spectra of air-dried PUASM peptide films with tentative secondary structures assignments. The results from experiments on the day of dissolving and incubation for 40 days at 4°C (39.2°F)

| EQB50682.1_332_355 | | | | | |
|---|---|---|---|---|---|
| after dissolving | | after 40 days | | band assignment | |
| band pos. [cm$^{-1}$] | area [%] | pos. [cm$^{-1}$] | area [%] | Amide | corresp. structure |
| 1695 | 6 | 1696 | 4 | I' | $\beta$-sheets & $\beta$-turns |
| 1677 | 16 | 1679 | 10 | | $\beta$-sheets & $\beta$-turns |
| 1663 | 19 | 1665 | 14 | | $\beta$-sheets & $\beta$-turns |
| 1648 | 14 | 1651 | 17 | | $\alpha$-helices & coils |
| **1631** | **14** | **1632** | **13** | | **$\beta$-sheets** |
| **1621** | **18** | **1620** | **27** | | **aggregated strands** |
| 1554 | 6 | 1552 | 8 | II' | $\alpha$-helices |
| 1538 | 7 | 1536 | 7 | | $\beta$-sheets & turns |

| EQB50683.1_9_31 | | | | | |
|---|---|---|---|---|---|
| after dissolving | | after 40 days | | band assignment | |
| band pos. [cm$^{-1}$] | area [%] | pos. [cm$^{-1}$] | area [%] | Amide | corresp. structure |
| 1694 | 3 | 1695 | 3 | I' | $\beta$-sheets & $\beta$-turns |
| 1679 | 11 | 1676 | 10 | | $\beta$-sheets & $\beta$-turns |
| 1664 | 12 | 1663 | 8 | Amide I' | $\beta$-sheets & $\beta$-turns |
| 1650 | 12 | 1650 | 12 | | $\alpha$-helices & coils |
| **1630** | **31** | **1627** | **37** | | **$\beta$-sheets** |
| **1616** | **14** | **1612** | **15** | | **aggregated strands** |
| 1552 | 10 | 1551 | 9 | II' | $\alpha$-helices |
| 1535 | 7 | 1535 | 6 | | $\beta$-sheets & turns |

Table S4: **Spontaneous and induced prion formation rates of the PUASM motif.** Sol indicates a soluble state of the fusion protein; prion indicates an aggregated state appearing as foci in fluorescence microscopy. Induction of prion formation was achieved by contact with strains expressing the prion state.

| Condition | Duration | # | PUASM-GFP | PUASM-RFP | GFP-PUASM |
|---|---|---|---|---|---|
| Spontaneous prion formation after transfection | 5 days | sol | 30 | 30 | 29 |
| | | prion | 0 | 0 | 1 |
| | 11 days | sol | 30 | 30 | 26 |
| | | prion | 0 | 0 | 4 |
| | 18 days | sol | 30 | 30 | 24 |
| | | prion | 0 | 0 | 6 |
| | 32 days | sol | 30 | 30 | 21 |
| | | prion | 0 | 0 | 9 |
| | 49 days | sol | 30 | 30 | 20 |
| | | prion | 0 | 0 | 10 |
| | 75 days | sol | 30 | 30 | 18 |
| | | prion | 0 | 0 | 12 |
| Induced prion formation after contact | 12 hours | sol | 18 | 18 | 10 |
| | | prion | 0 | 0 | 8 |
| | 48 hours | sol | 18 | 18 | 7 |
| | | prion | 0 | 0 | 11 |
| | 96 hours | sol | 18 | 18 | 0 |
| | | prion | 0 | 0 | 18 |

### Figures

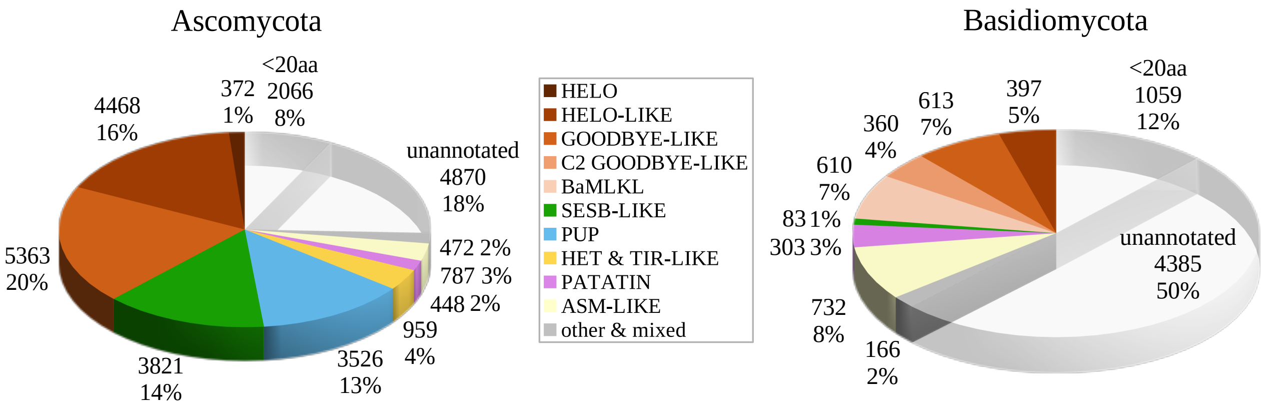


Figure S1: **Distribution of domain families in fungal NLR N-termini** with regard to phylum. See Results and Methods for details.

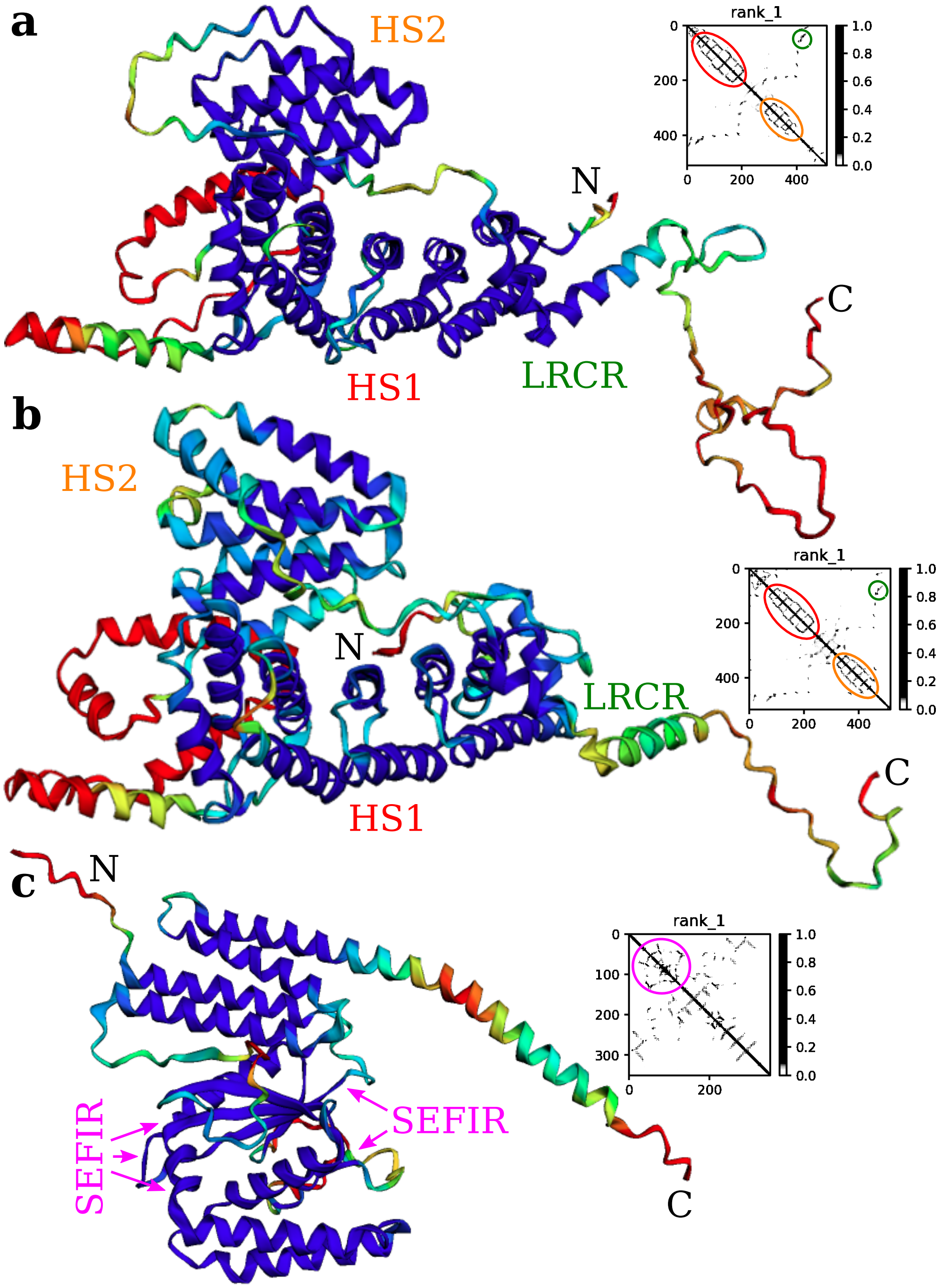


Figure S2: **Structural models of three unannotated domains** predicted with AlphaFold2. (**a**) KEY84097, (**b**) KFH66451, (**c**) PQE30966. Rainbow colors indicate model quality in terms of lDDT (below or 50: red, 60: yellow, 70: green, 80: cyan, above 90: blue). Insets show contact probability maps. Regions of special interests are marked with colored ellipses on insets and annotated on structural models. Notations: HS1,2—helix stretch 1,2; LRCR—long-range contact region; SEFIR—region matching Pfam SEFIR domain (PF08357).

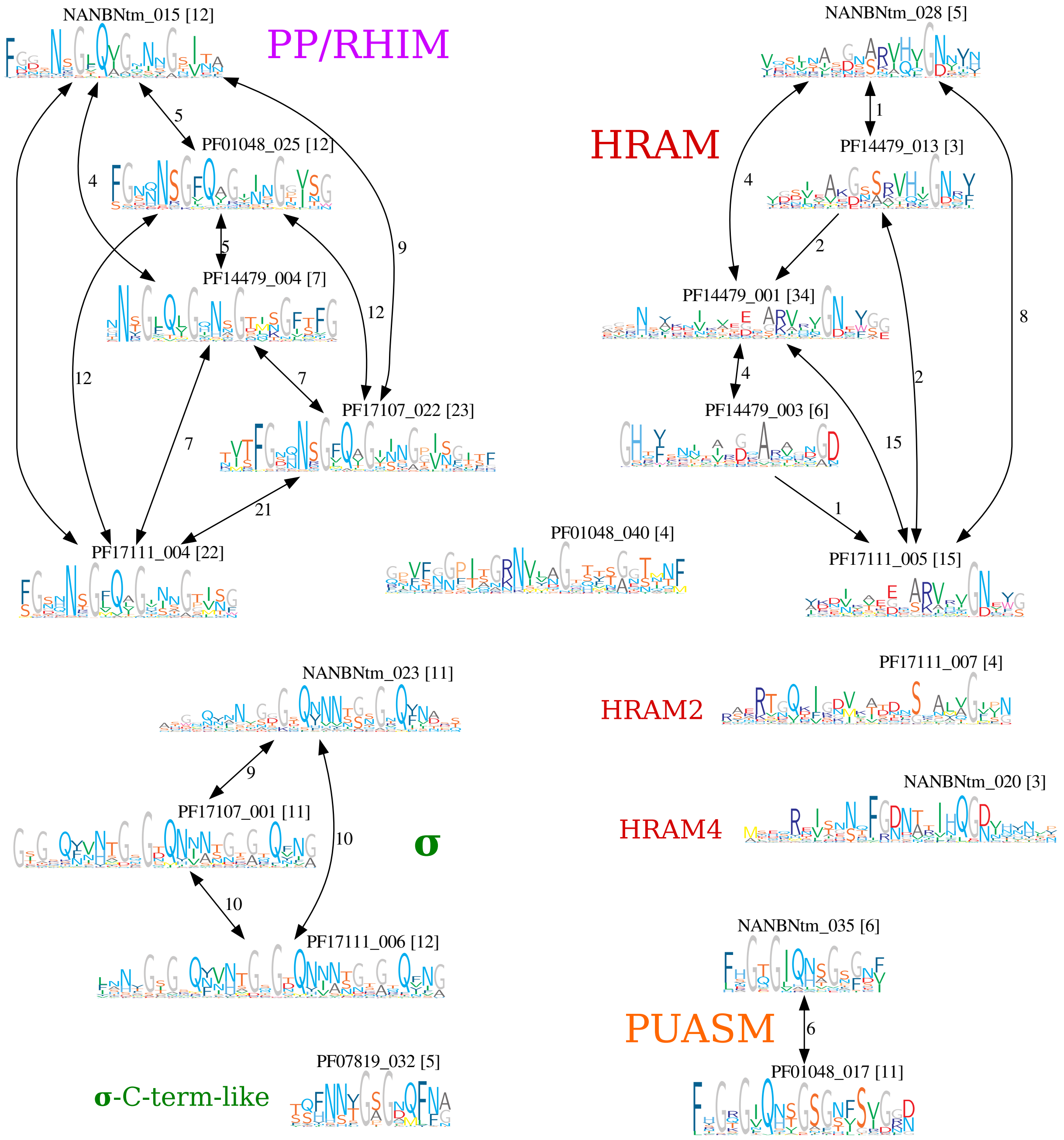


Figure S3: **Clustering amyloid signaling motifs** identified through the effector domain-primed search. Motif id indicates the effector domain (PF...) or nucleotide-binding domain (NANB) next to which the seed motif was originally extracted (see Methods) and rank in the MEME extraction. The number in brackets represents the number of pairs for each profile, the number on the arrow joining the motifs indicates the number of common matching underlying sequences.

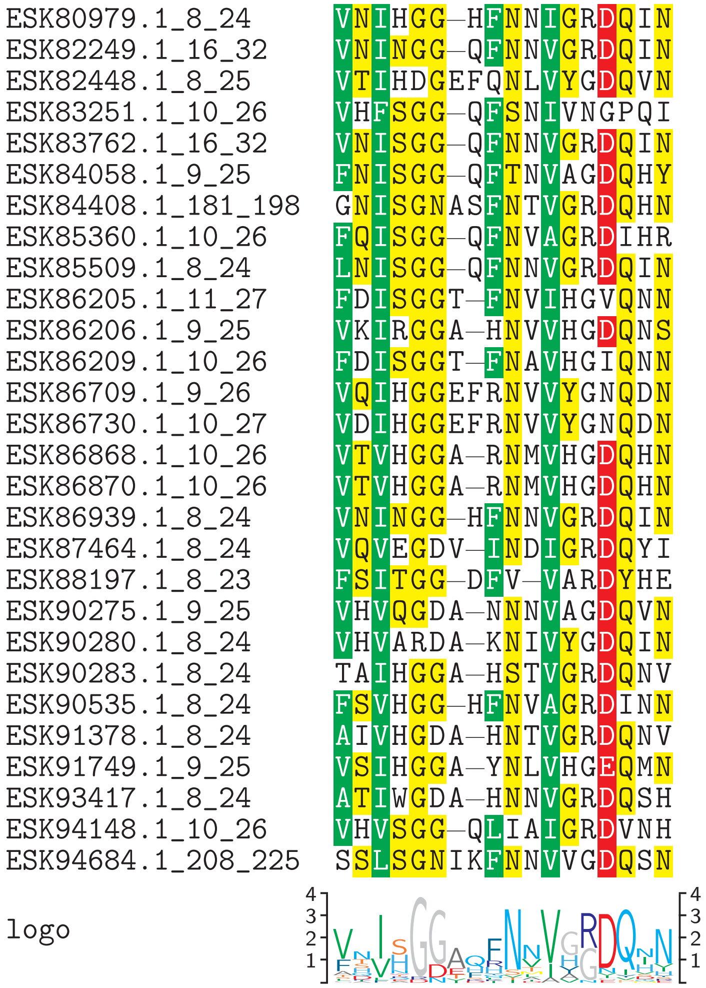


Figure S4: **Alignment of candidate amyloid signaling motifs in *Moniliophthora roreri*** (strain MCA 2997). NLR-side N-terminal motifs and MLKL-like-side C-terminal motifs were aligned with Clustal Omega in *auto* mode.

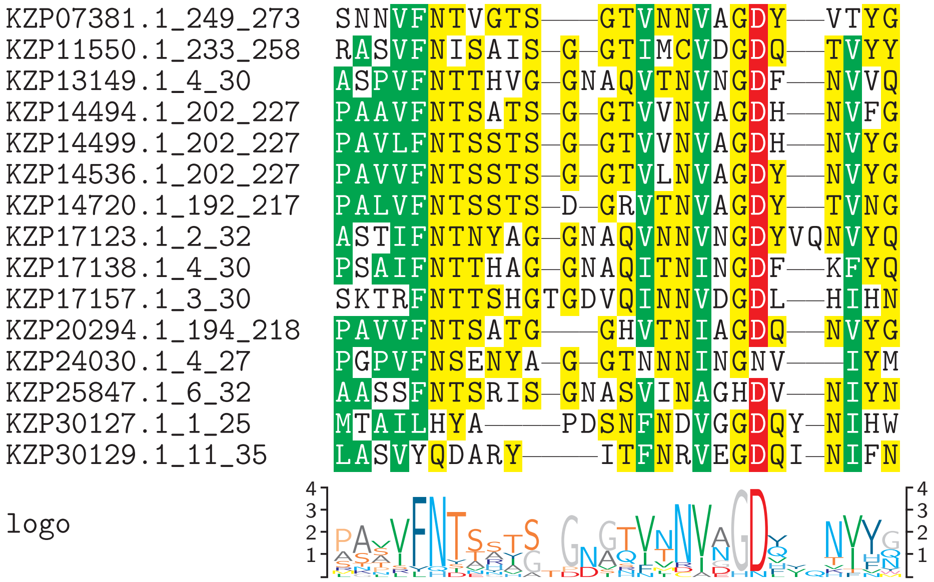


Figure S5: **Alignment of candidate amyloid signaling motifs in *Fibularhizoctonia* sp. CBS 109695.** Eight NLR-side N-terminal motifs, five intra-protein motifs from MLKL-like–NLR proteins, and two MLKL-like-side C-terminal motifs (KZP11550, KZP14720) were aligned with Mafft in *linsi* mode.

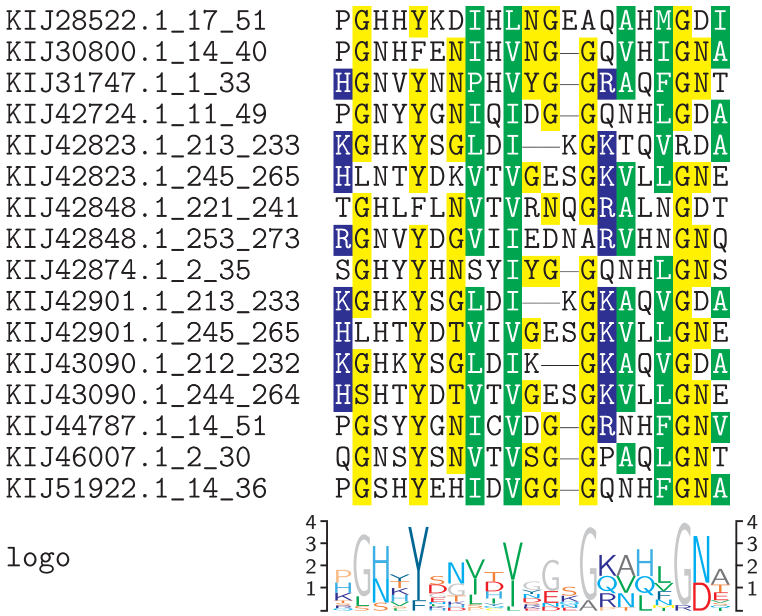


Figure S6: **Alignment of candidate amyloid signaling motifs in *t*** (strain SS14). Eight NLR-side N-terminal motifs and four HeLo-side double C-terminal HRAMs were aligned with Mafft in *linsi* mode and refined manually.

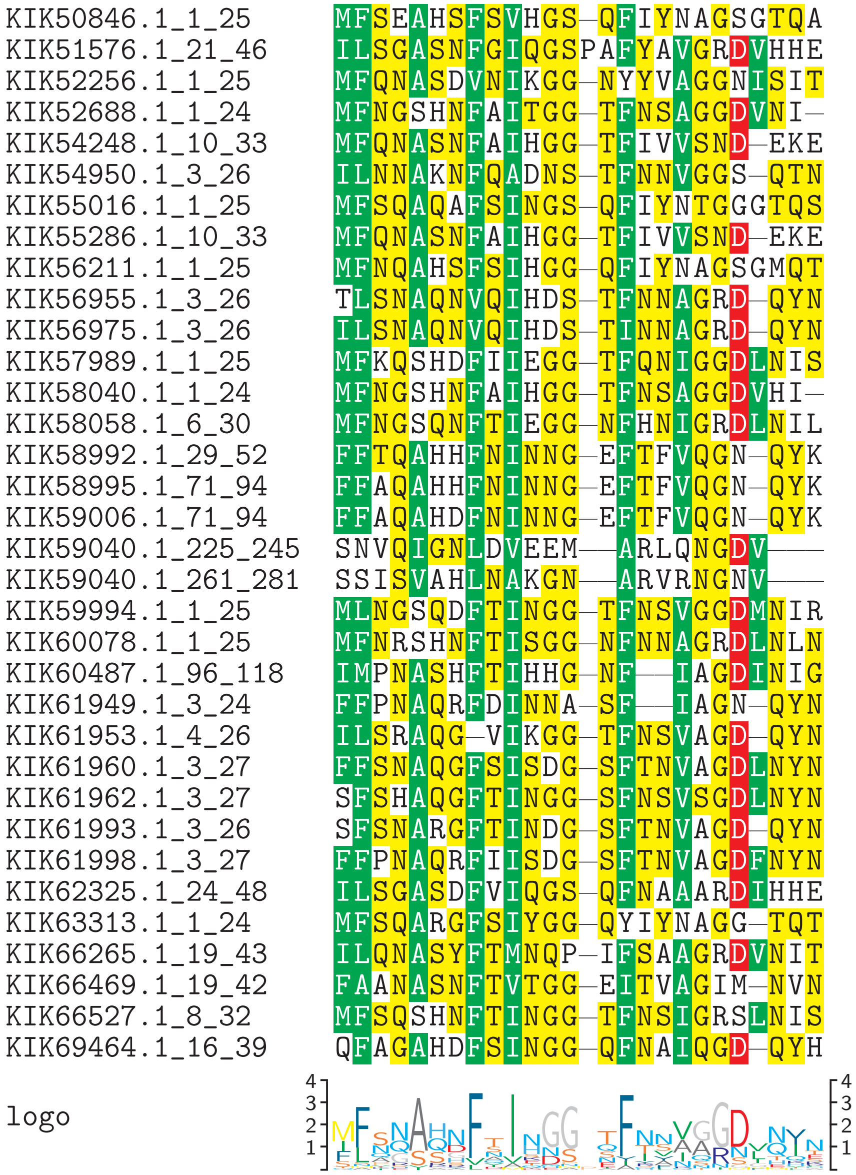


Figure S7: **Alignment of candidate amyloid signaling motifs in *Gymnopus luxurians*** (strain FD-317 M1). NLR-side N-terminal motifs of NLR05/08/22/44 family were aligned with Mafft in *linsi* mode and trimmed manually. A HeLo-side double C-terminal HRAM was aligned to them manually.

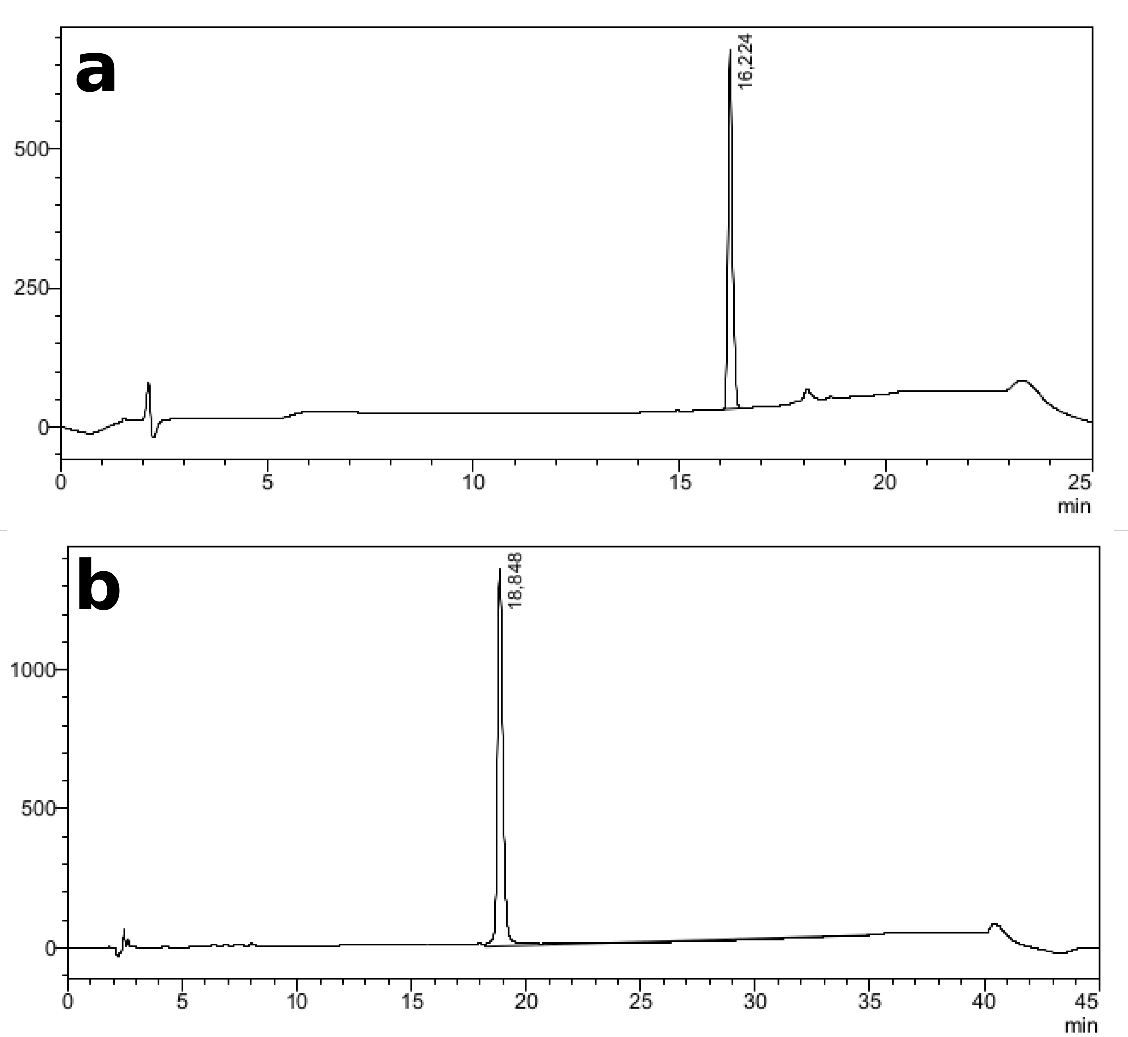


Figure S8: **Analytical HPLC chromatograms of synthesized PUASM peptides.** **a**) EQB50682.1_332_355, **b**) EQB50683.1_9_31. See also Table S1

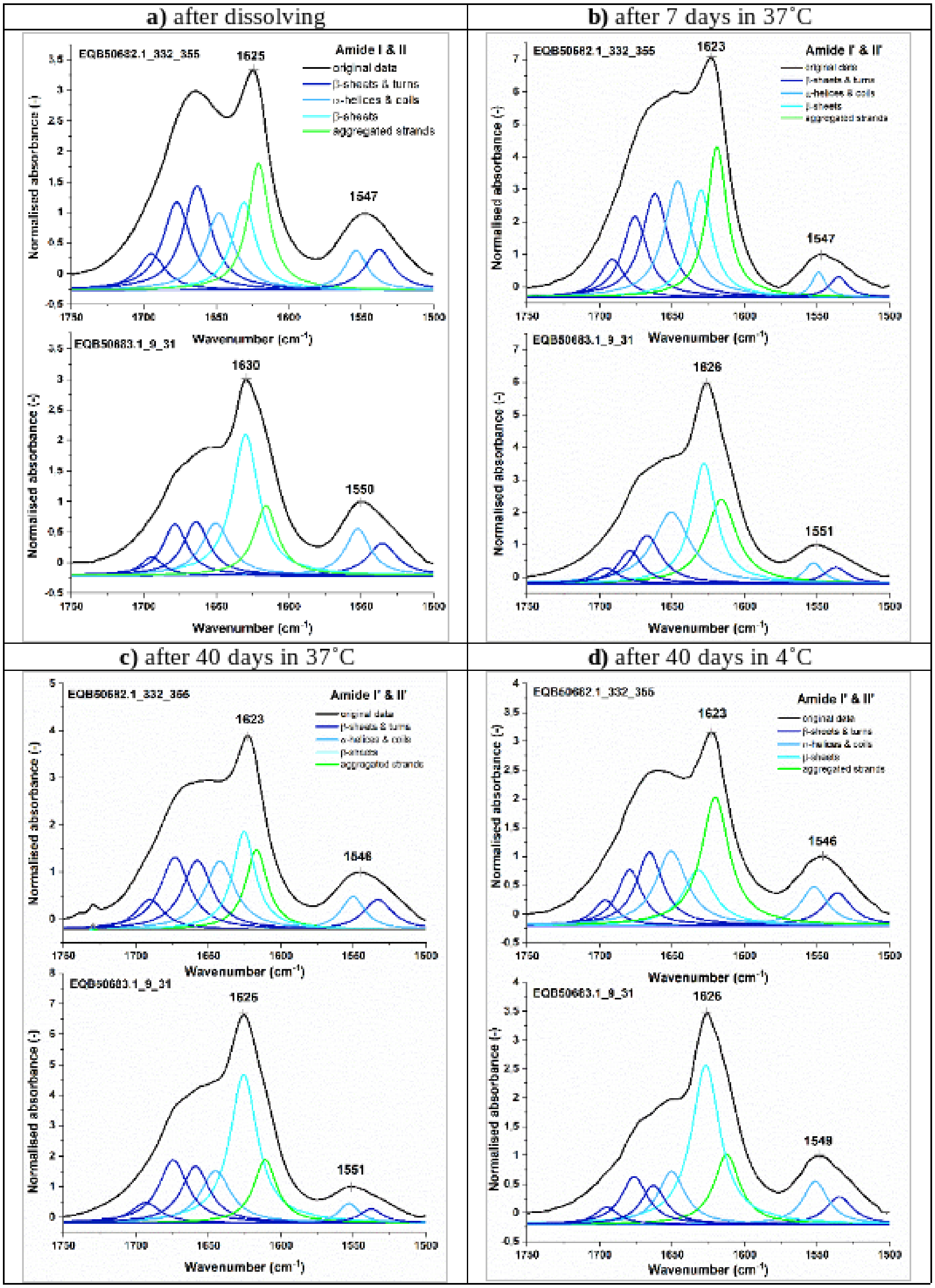


Figure S9: **Normalized ATR-FTIR spectra of air-dried peptide films** of EQB50682.1_332_355 and EQB50683.1_9_31 with sub-bands obtained from the curve fitting procedure in the amide bands region (1750–1500 cm$^{-1}$) registered at a temperature of 20°: directly after dissolving (**a**), after 7 days (**b**) and 40 days of incubation process at 37°C (98.6°F) (**c**), and after 40 days of incubation at 4°C (39.2°F) (**d**).

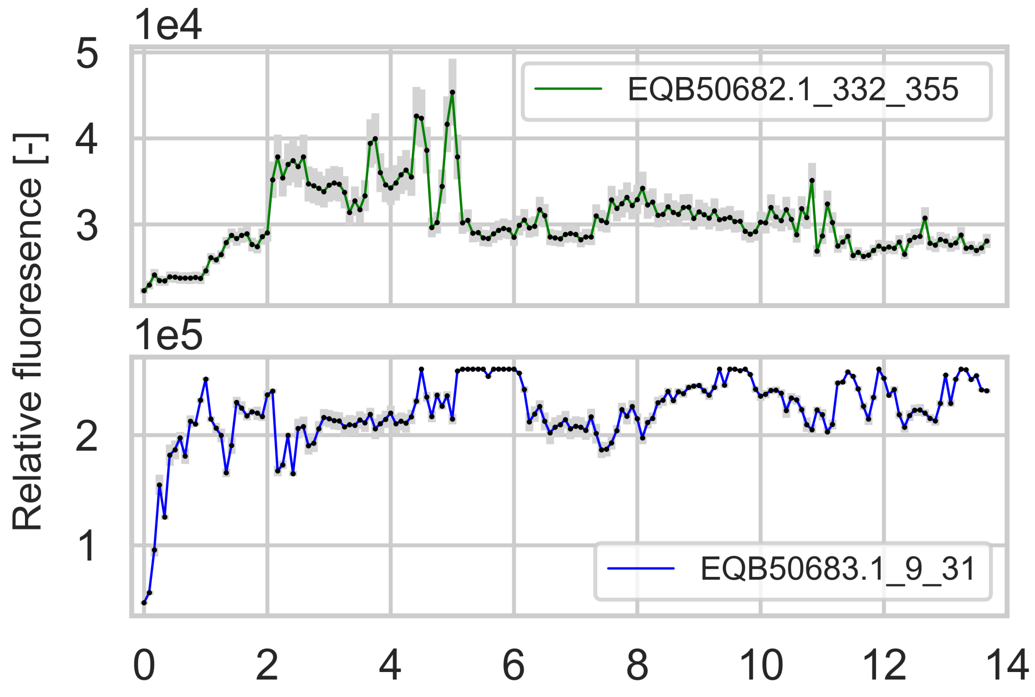


Figure S10: **Aggregation kinetics** of peptides EQB50682.1_332_355 and EQB50683.1_9_31. See Results and Methods for details.
