## Supplementary figures and images for "Exploring a diverse world of effector domains and amyloid signaling motifs in fungal NLR proteins"

### msa_coverage.filtered.png

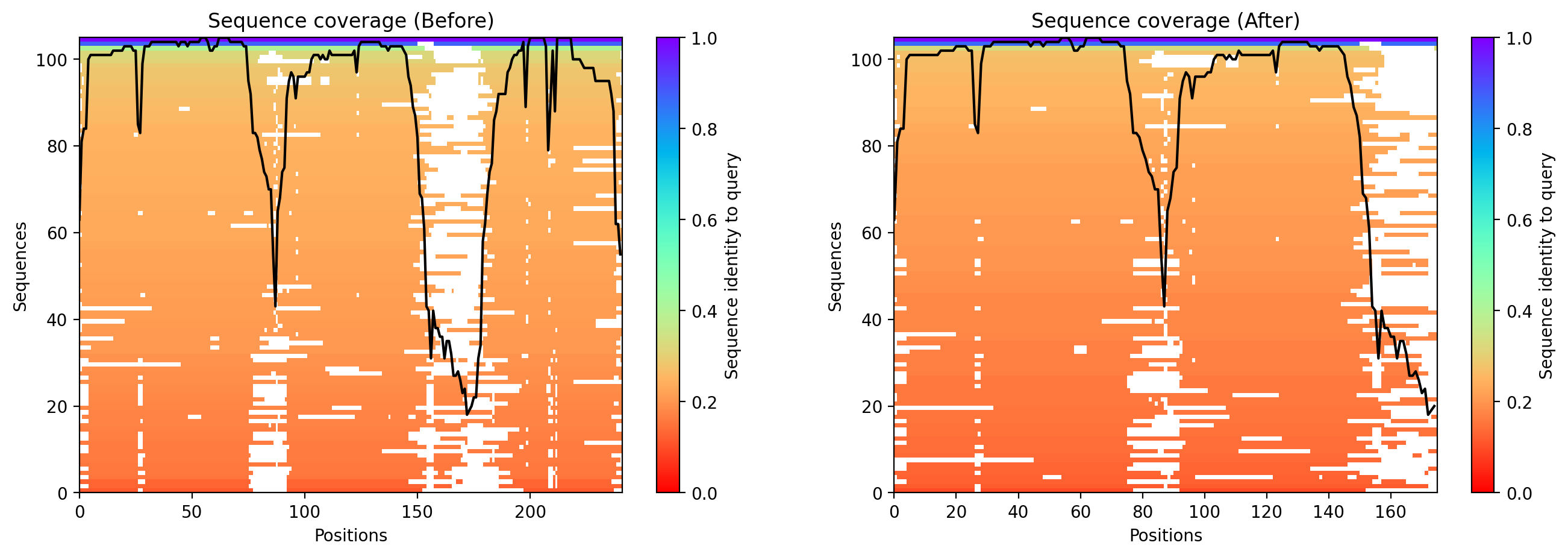

### msa_coverage.filtered.png

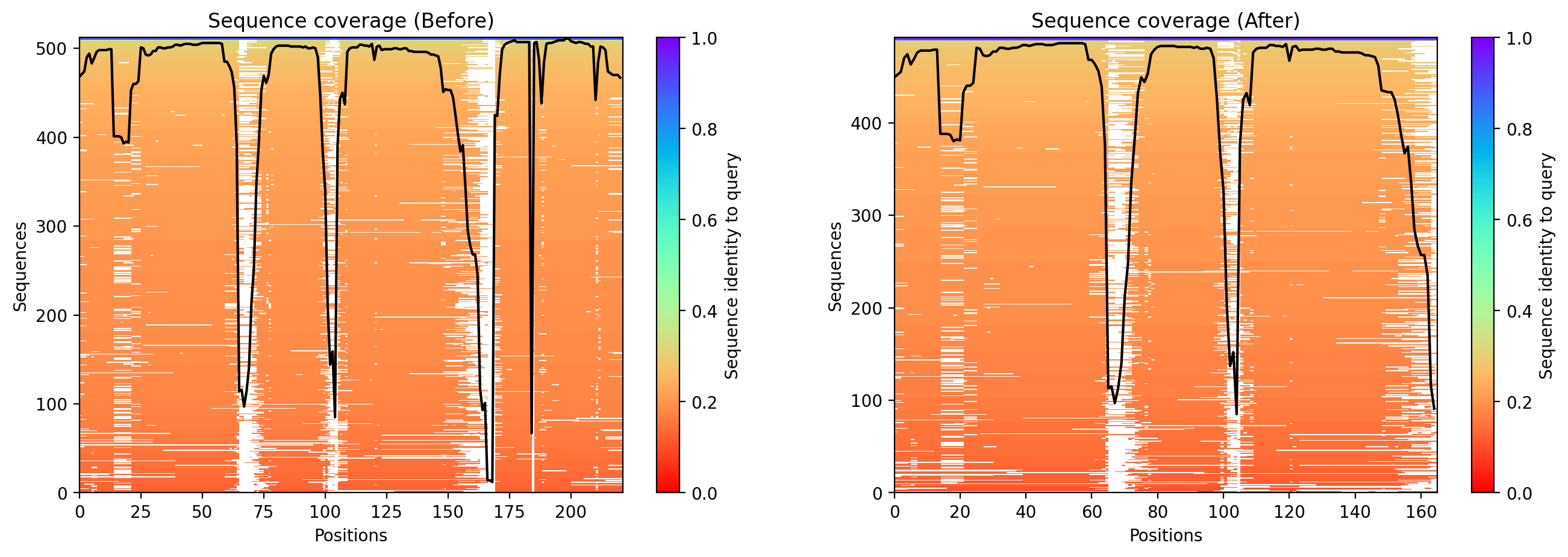

### msa_coverage.filtered.png

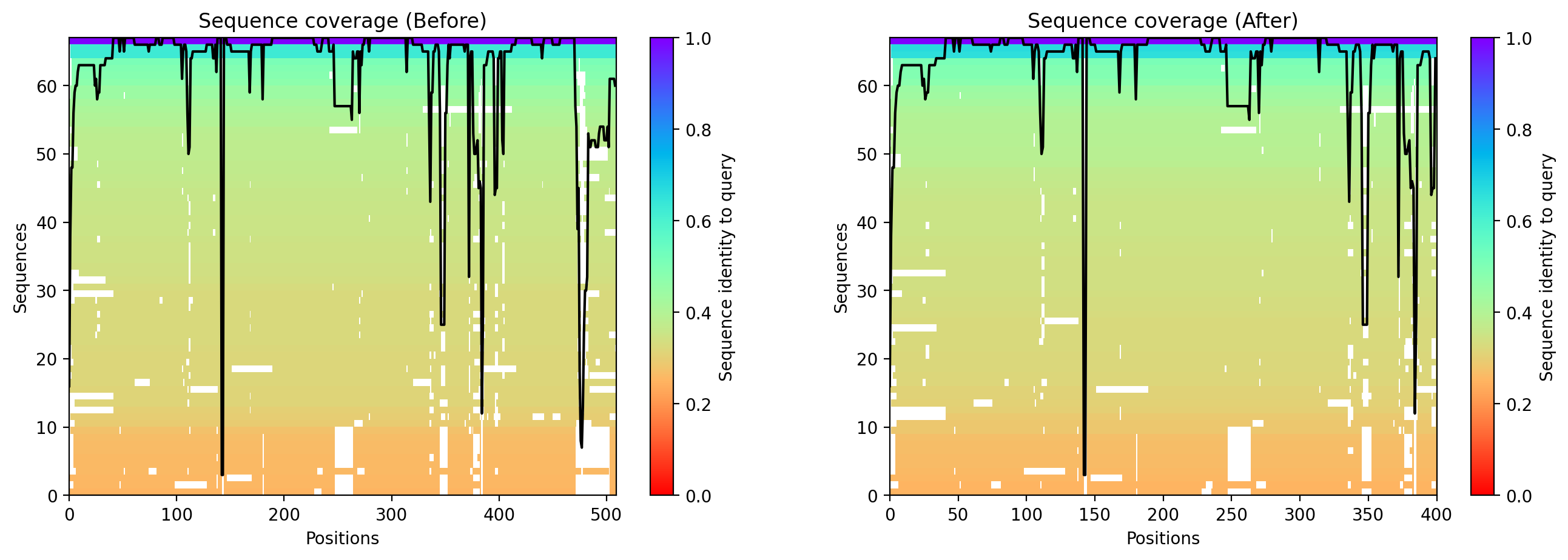

### msa_coverage.filtered.png

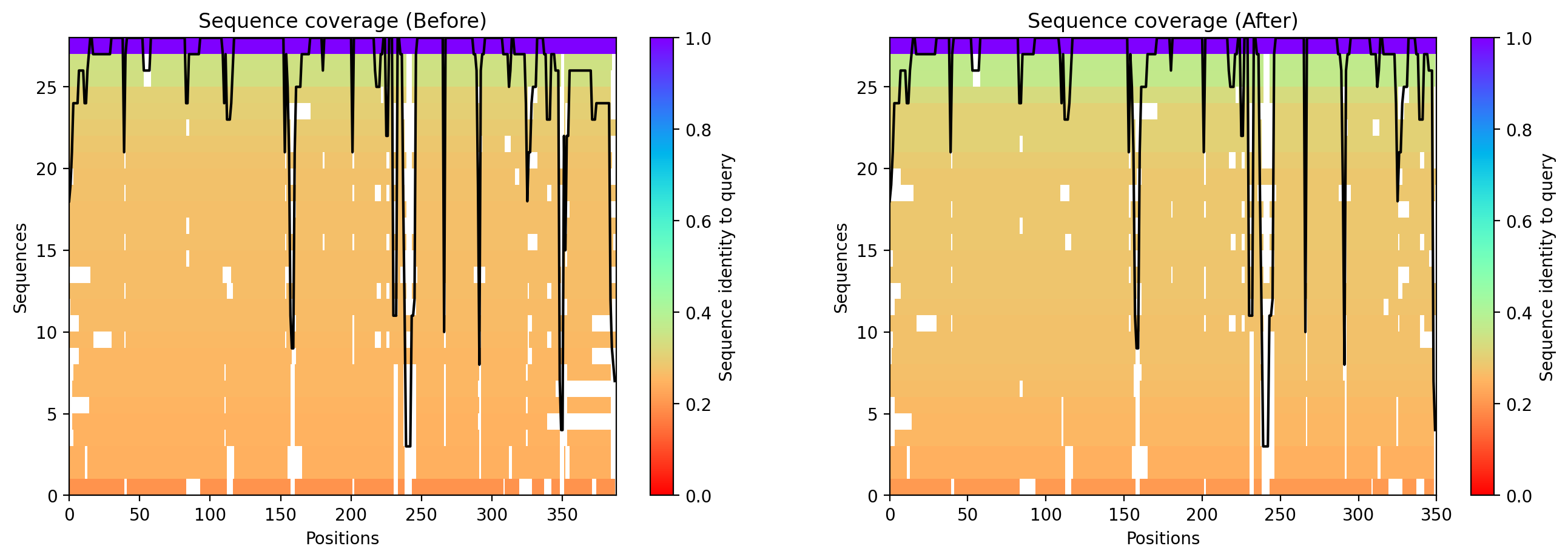

### msa_coverage.png

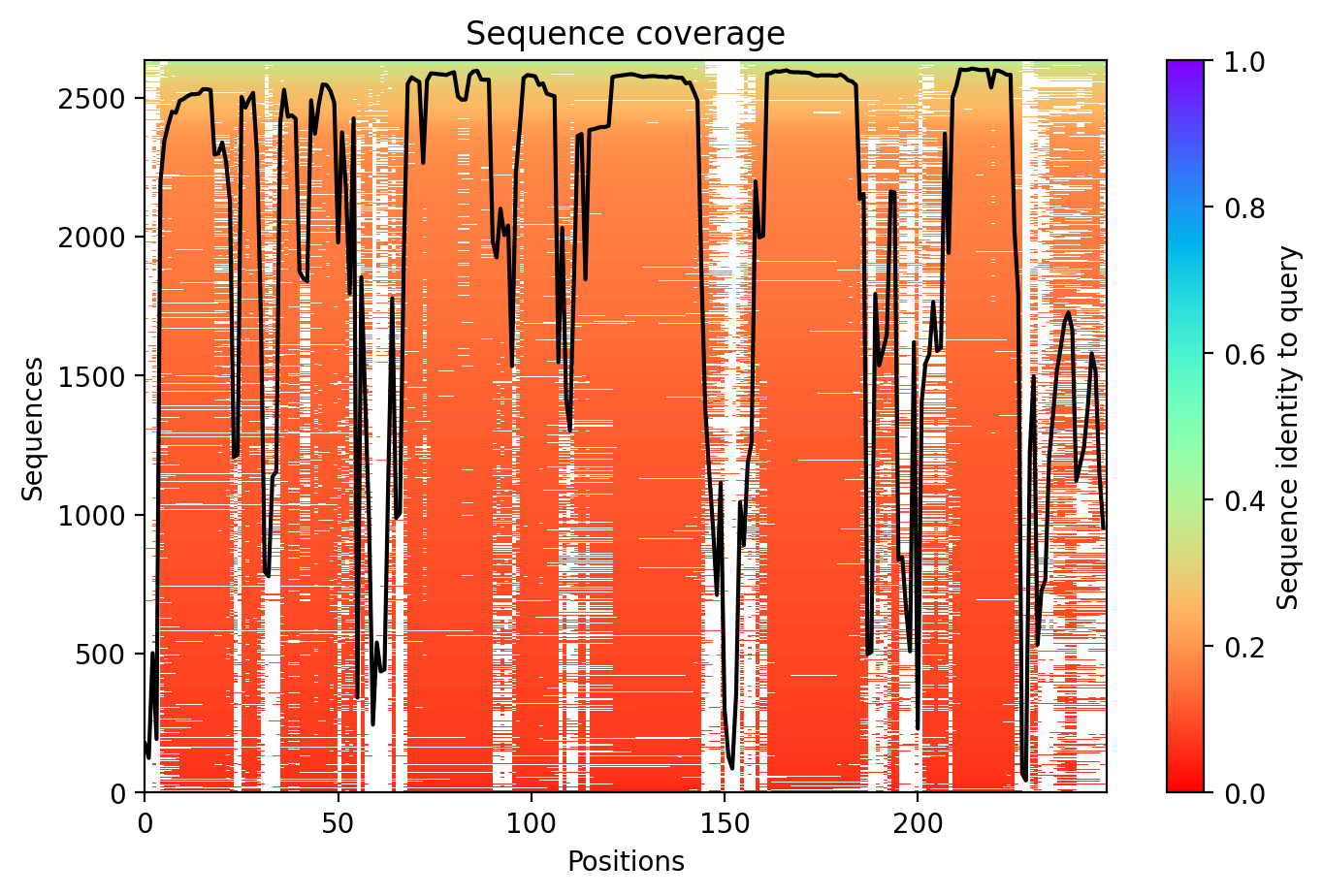

### msa_coverage.png

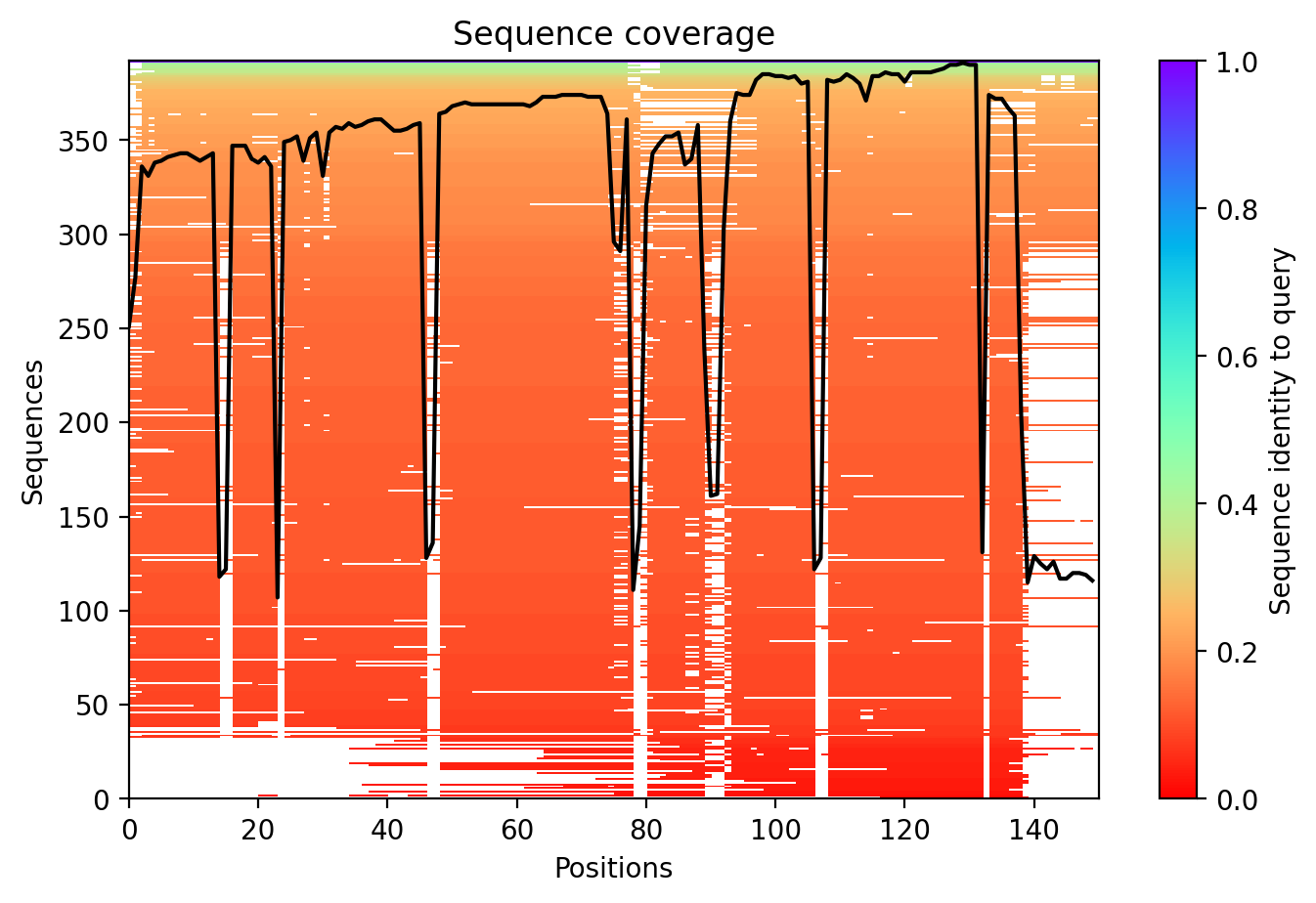

### msa_coverage.png

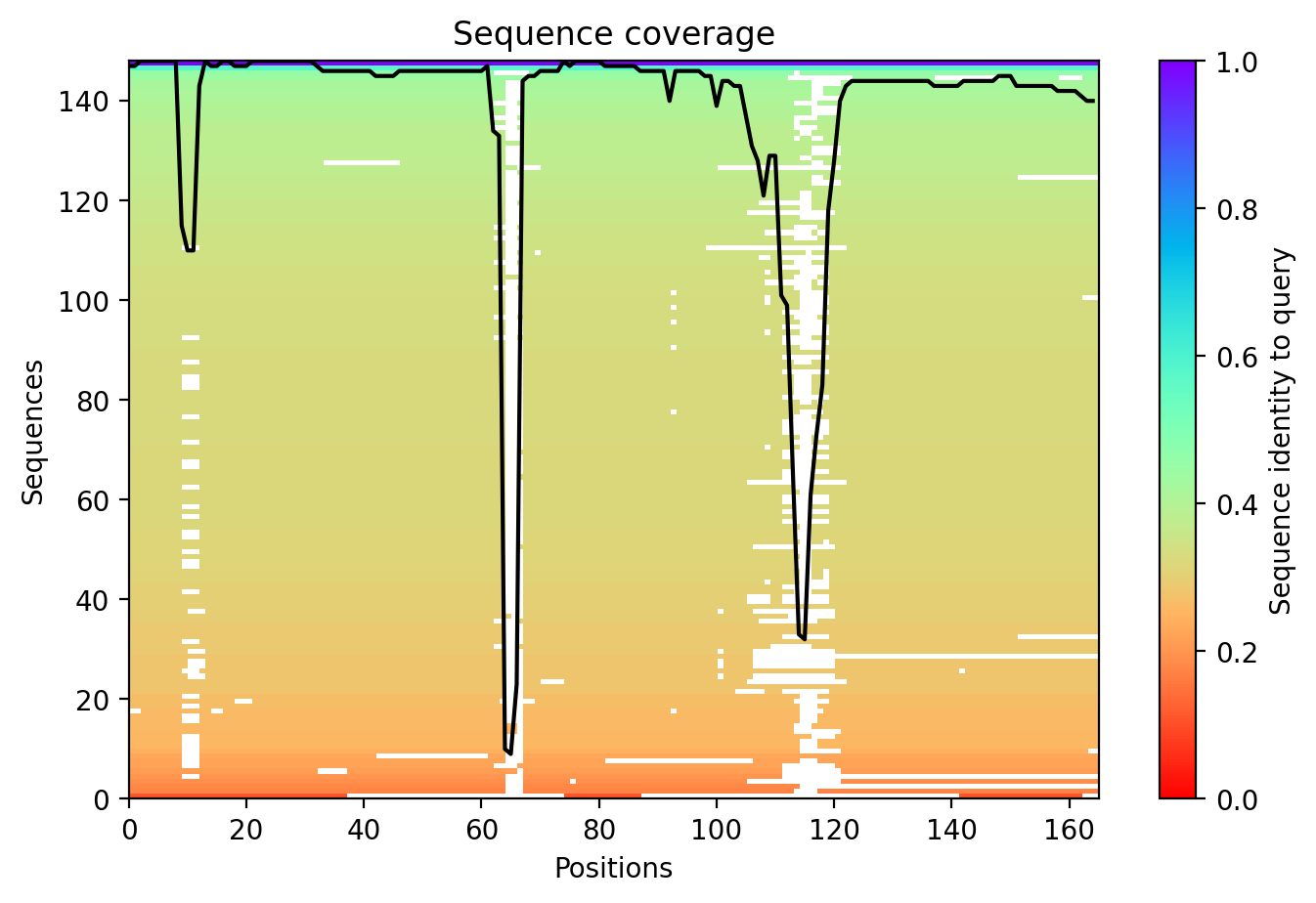

### msa_coverage.png

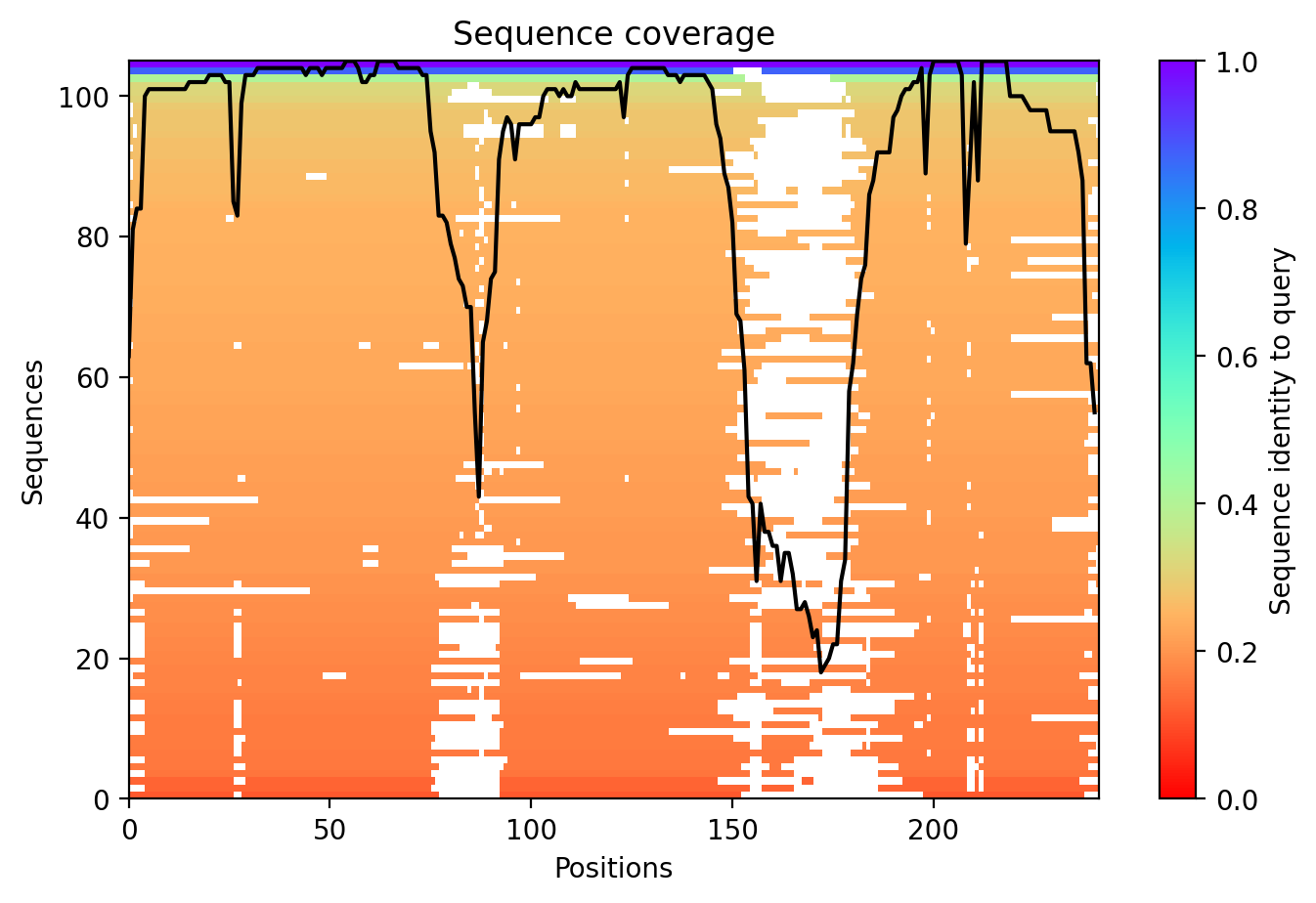

### msa_coverage.png

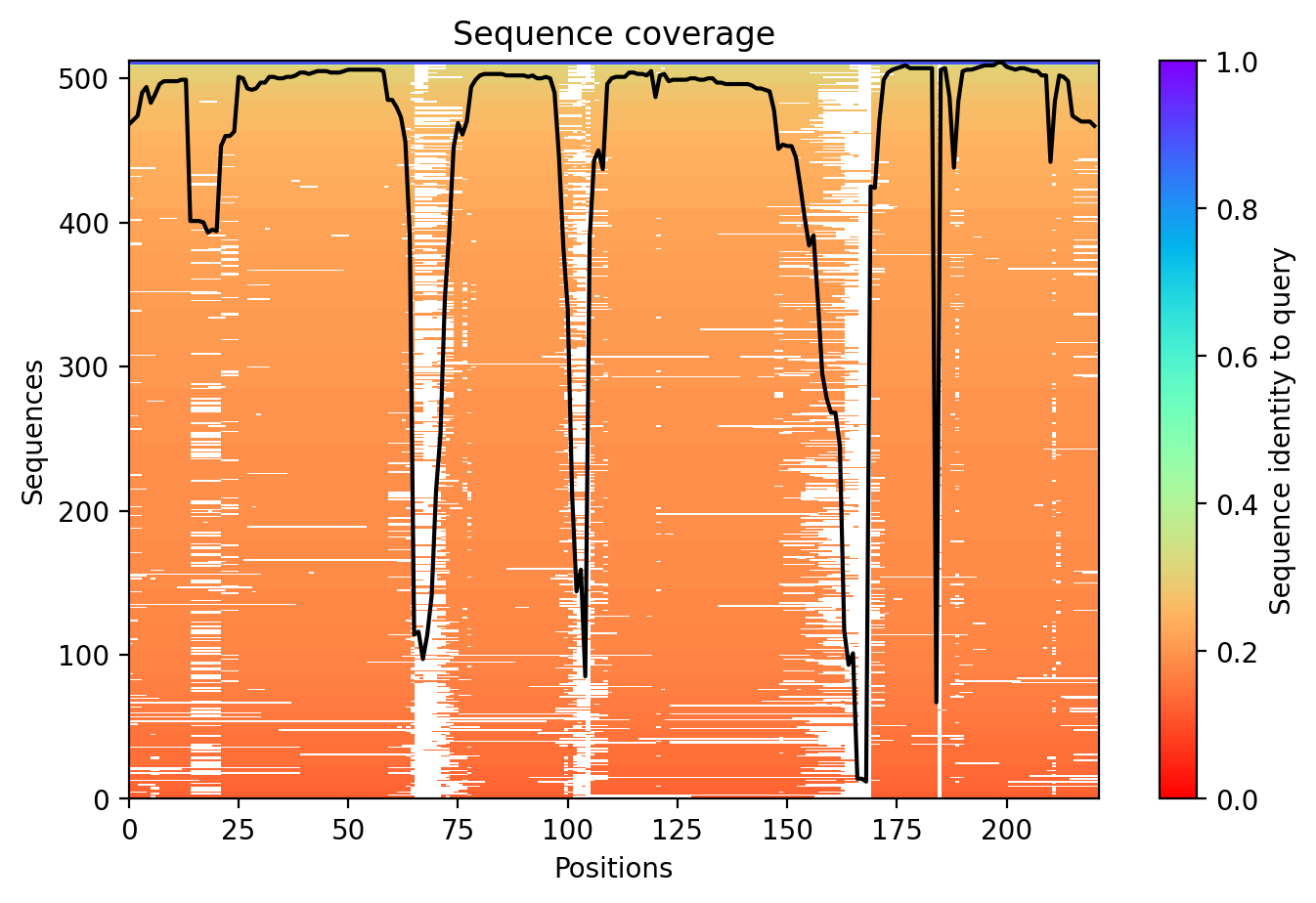

### msa_coverage.png

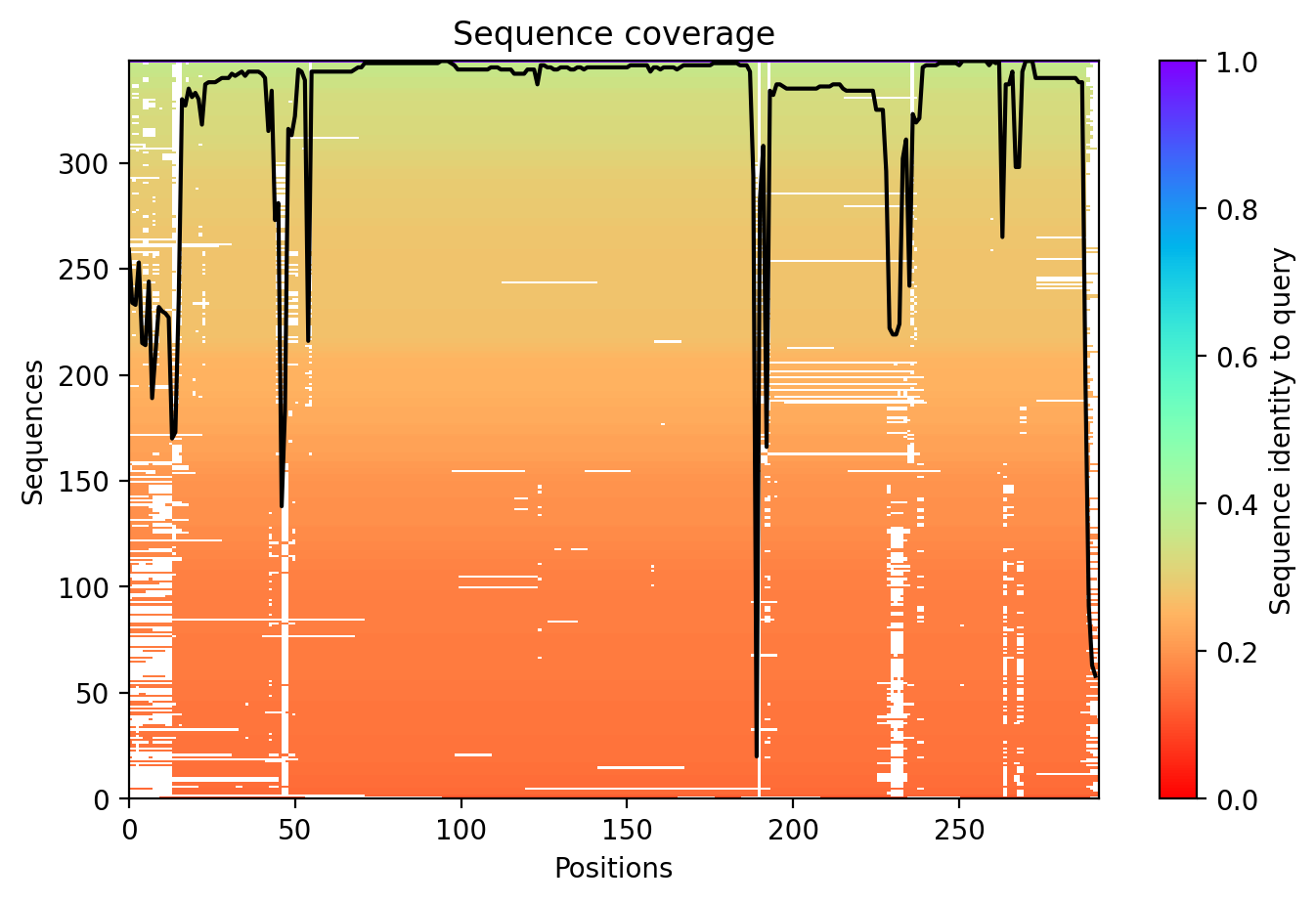

### msa_coverage.png

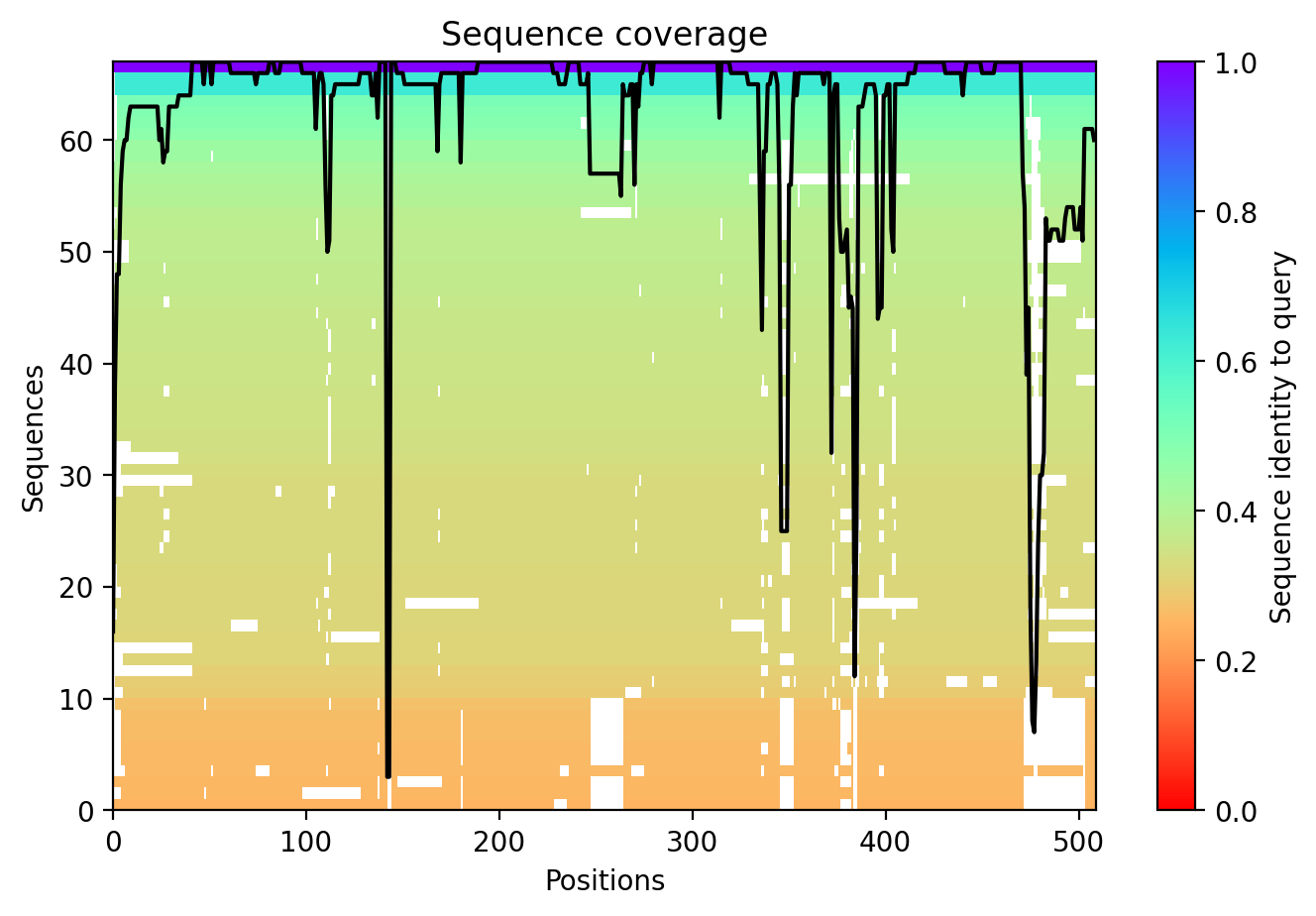

### msa_coverage.png

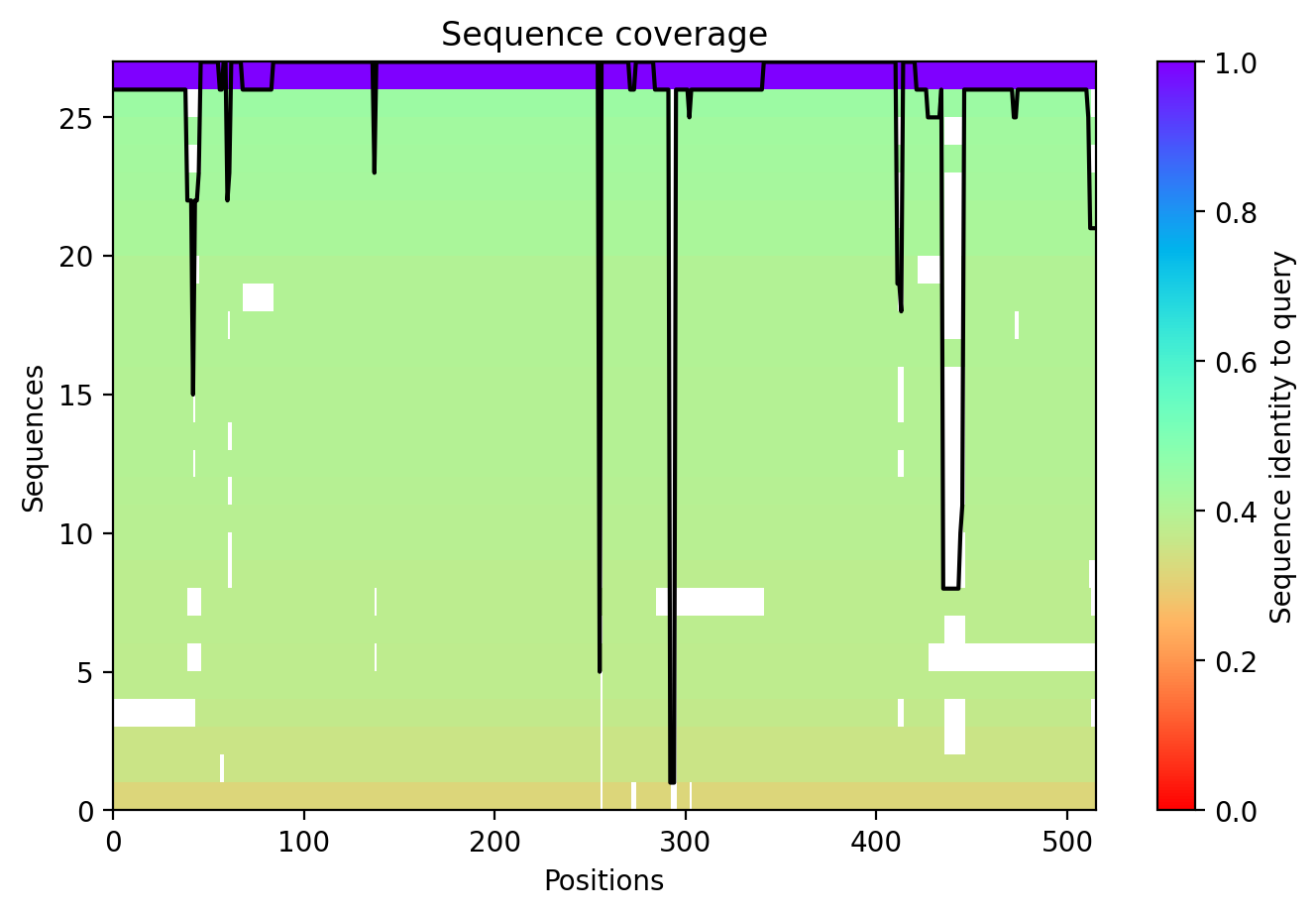

### msa_coverage.png

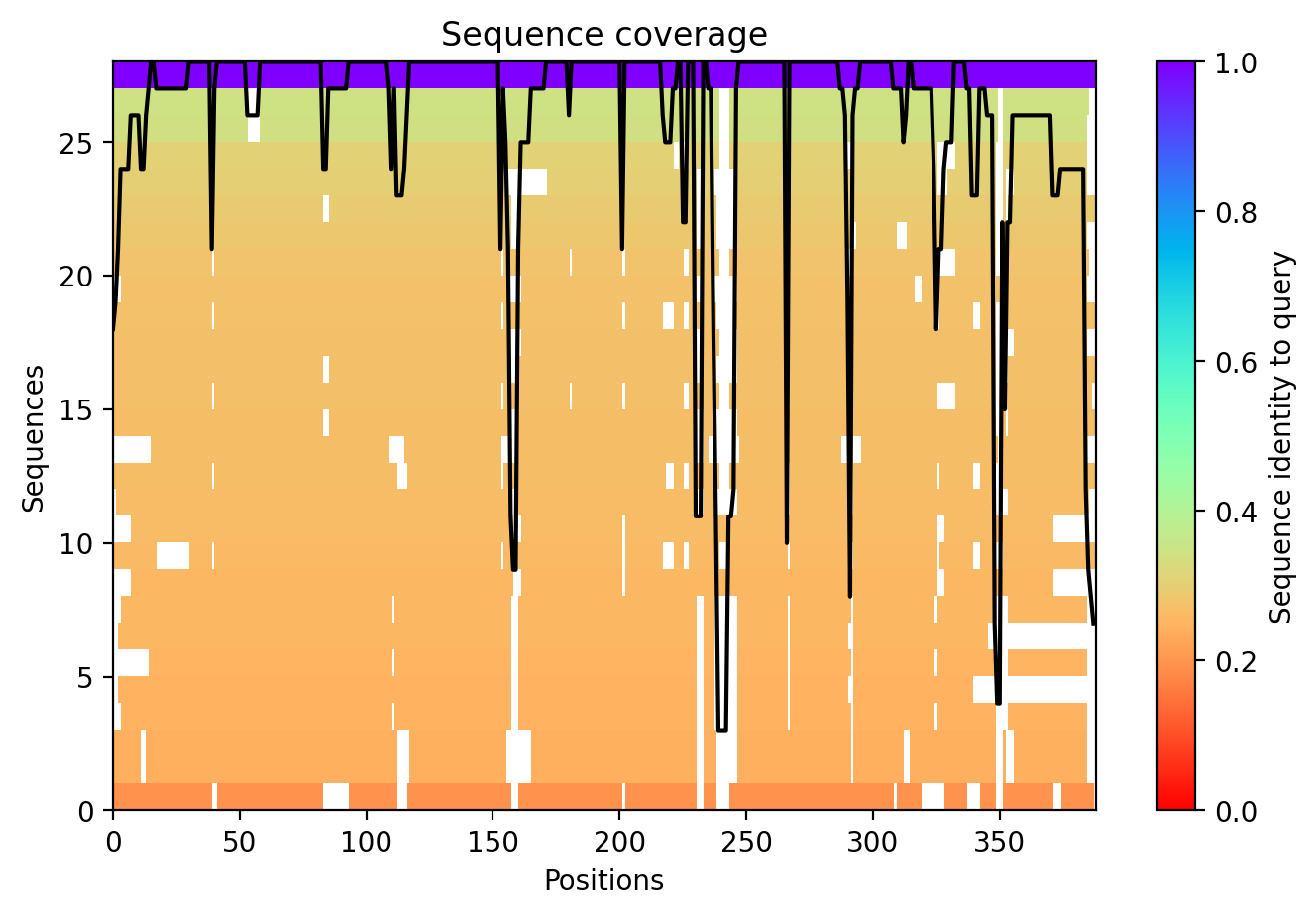

### predicted_alignment_error.png

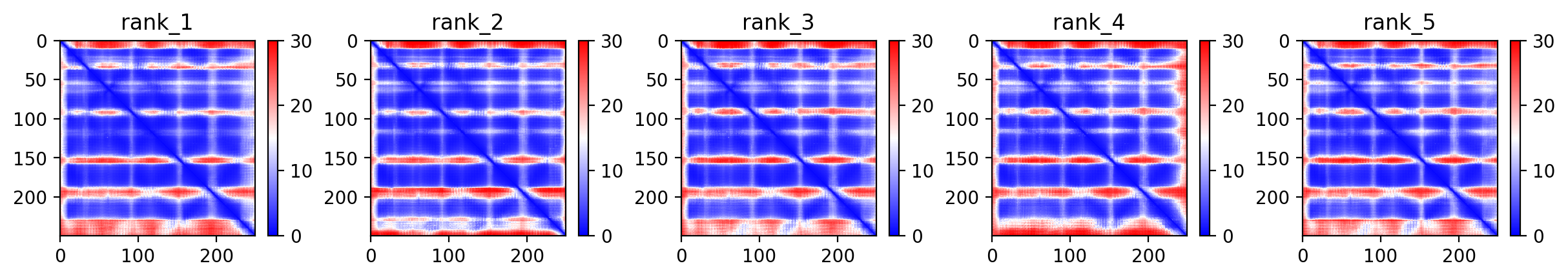

### predicted_alignment_error.png

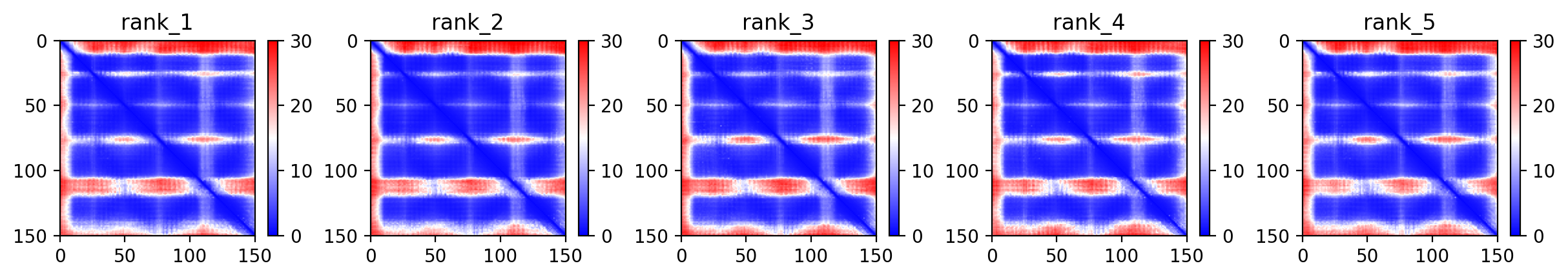

### predicted_alignment_error.png

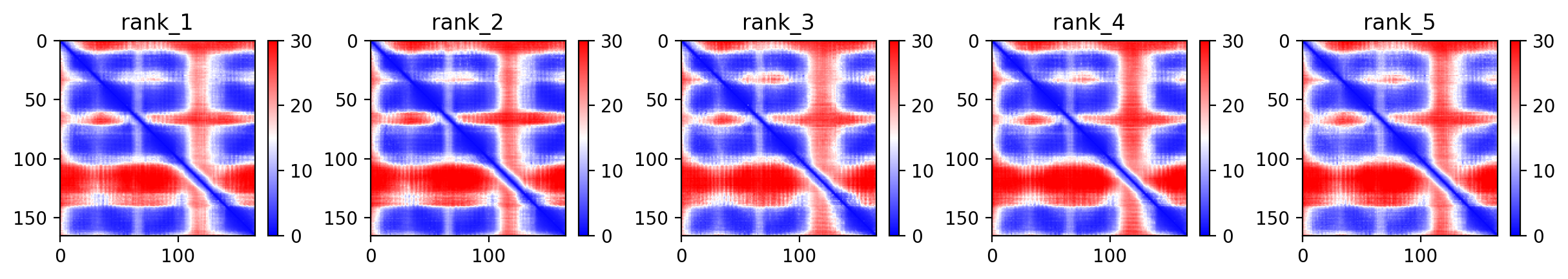

### predicted_alignment_error.png

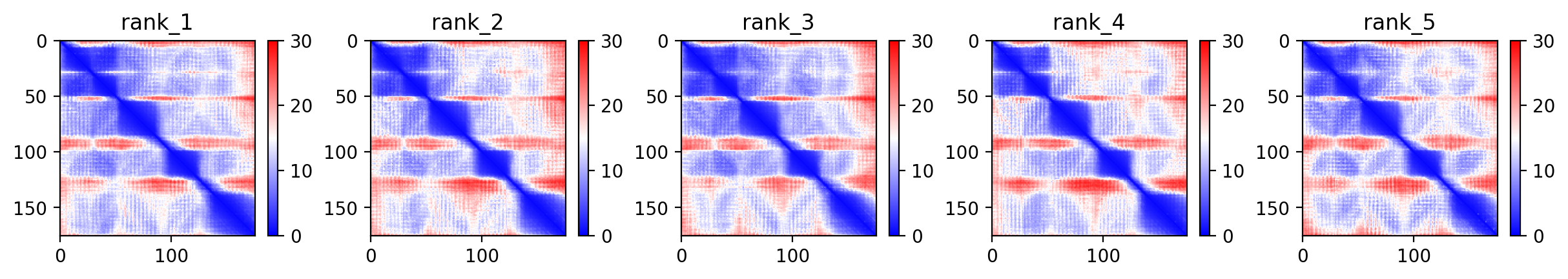

### predicted_alignment_error.png

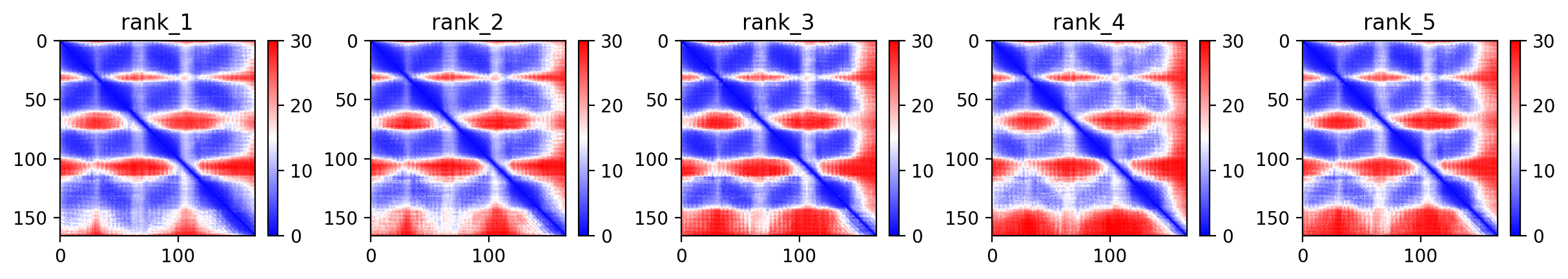

### predicted_alignment_error.png

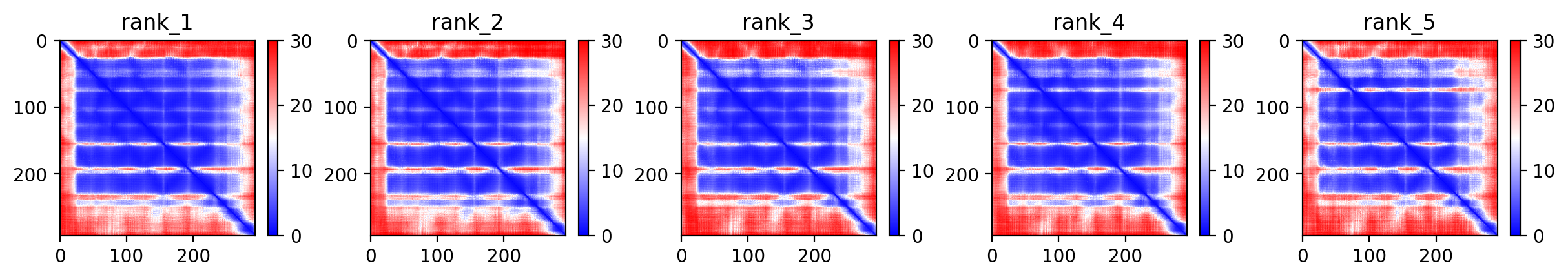

### predicted_alignment_error.png

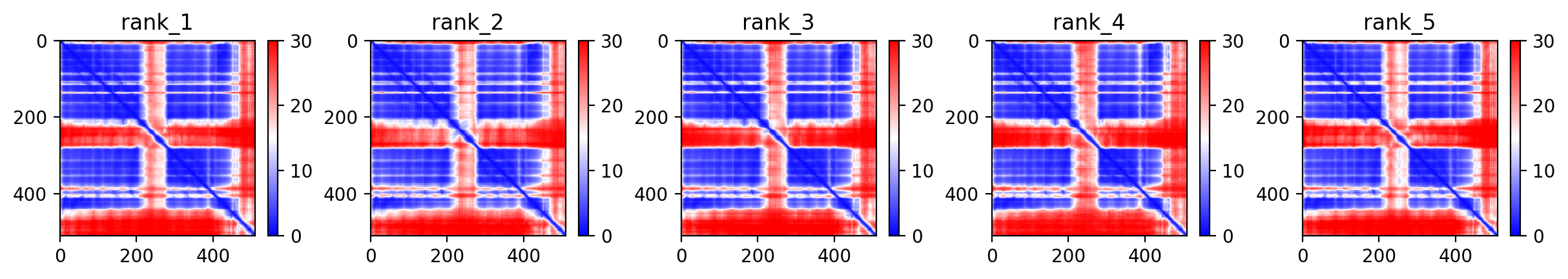

### predicted_alignment_error.png

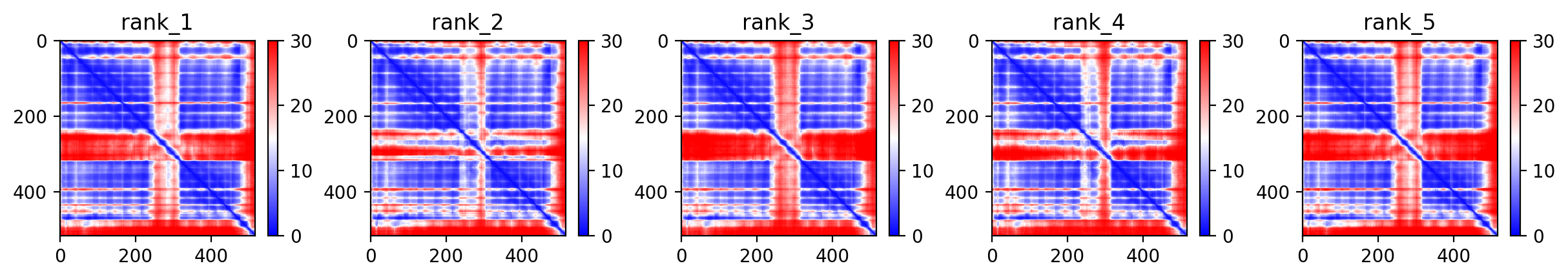

### predicted_alignment_error.png

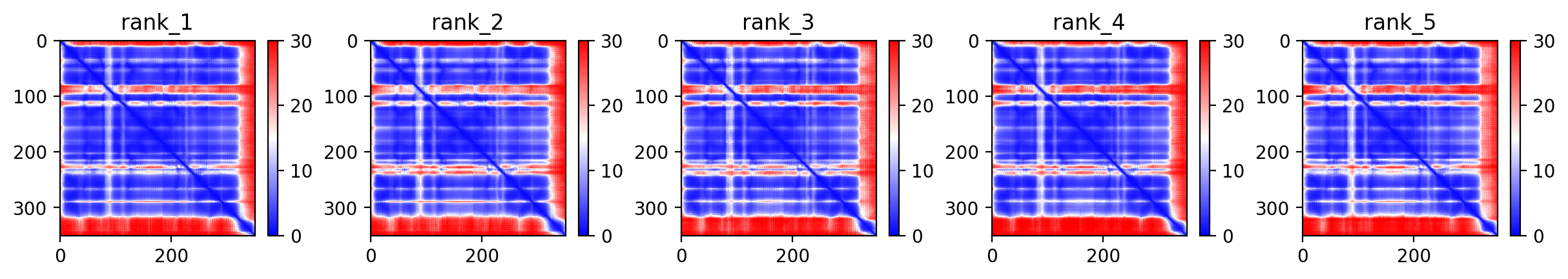

### predicted_contacts.png

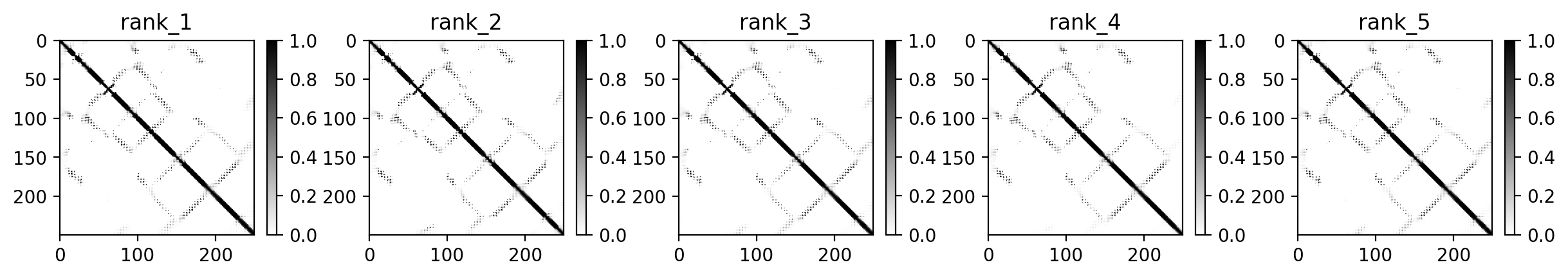

### predicted_contacts.png

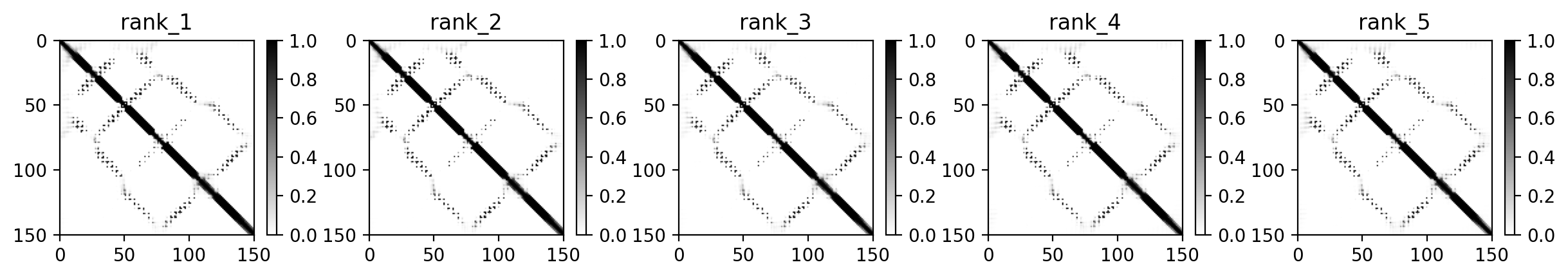

### predicted_contacts.png

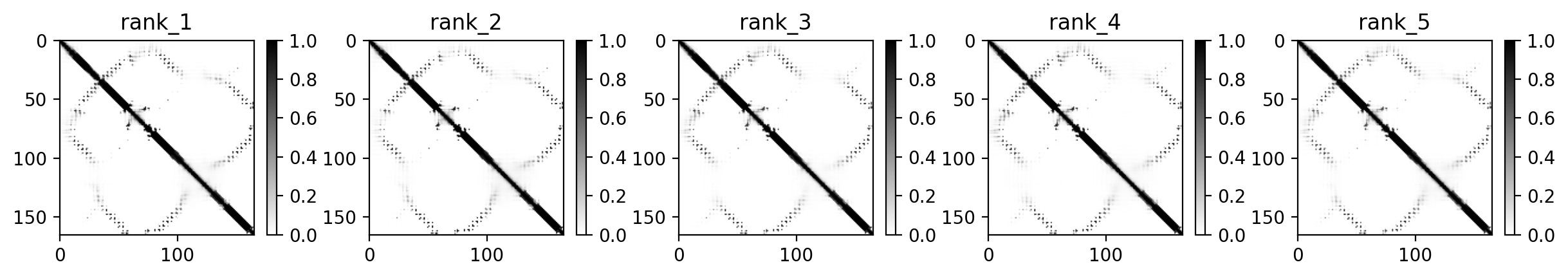

### predicted_contacts.png

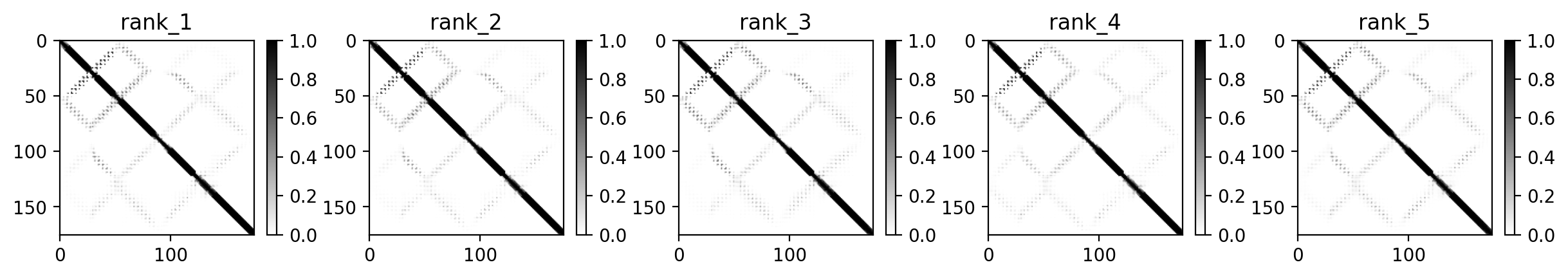

### predicted_contacts.png

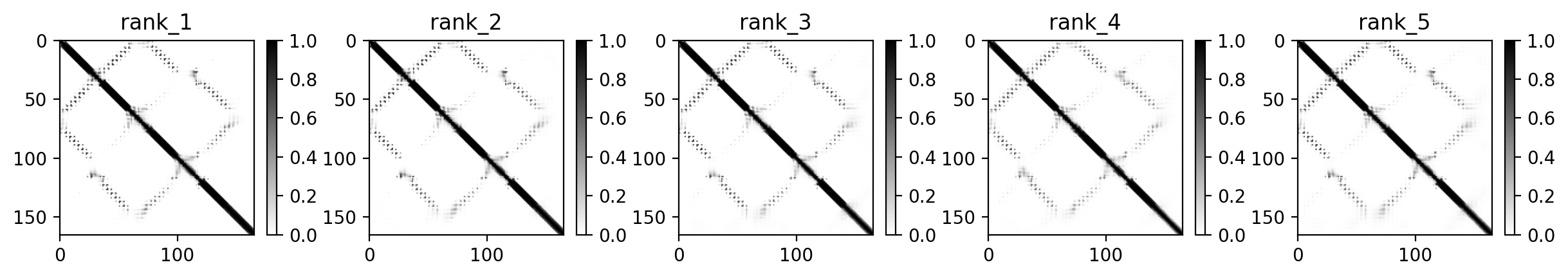

### predicted_contacts.png

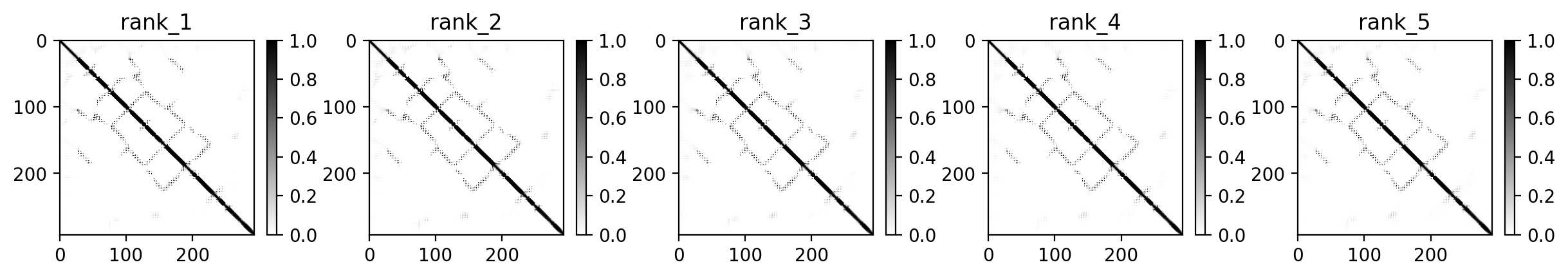

### predicted_contacts.png

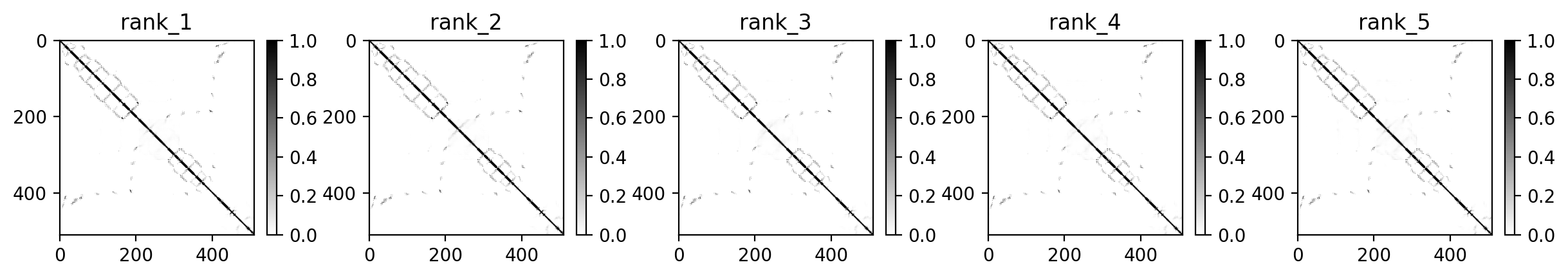

### predicted_contacts.png

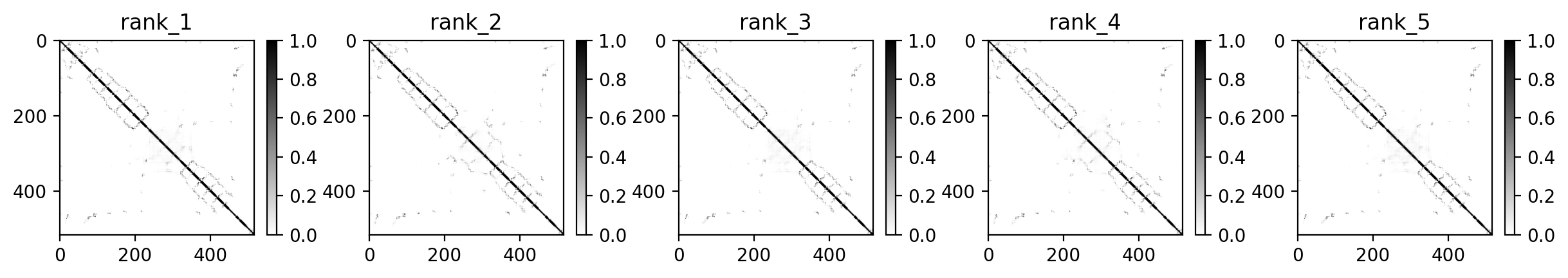

### predicted_contacts.png

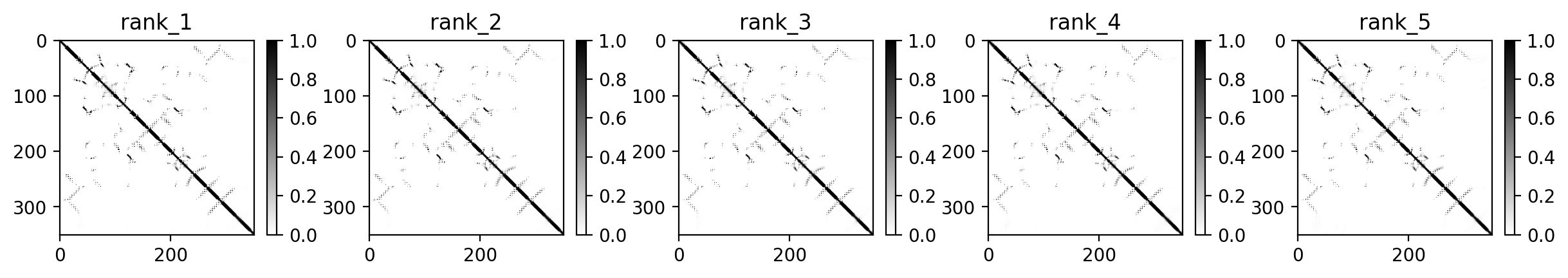

### predicted_distogram.png

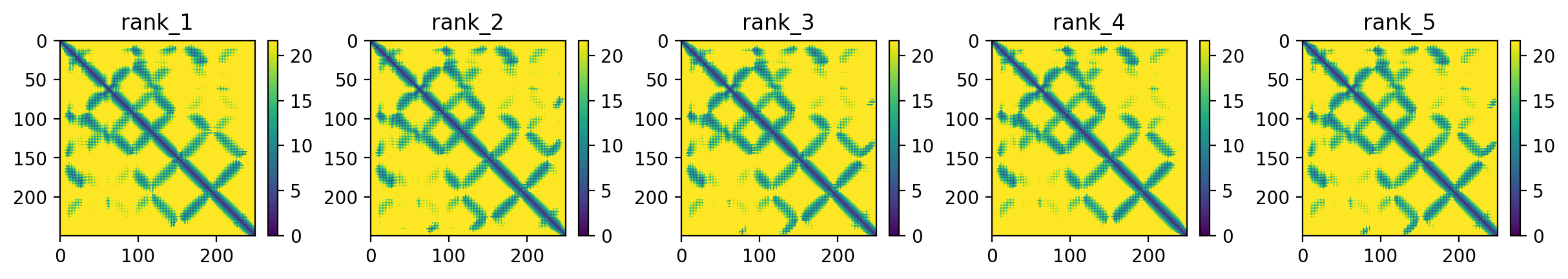

### predicted_distogram.png

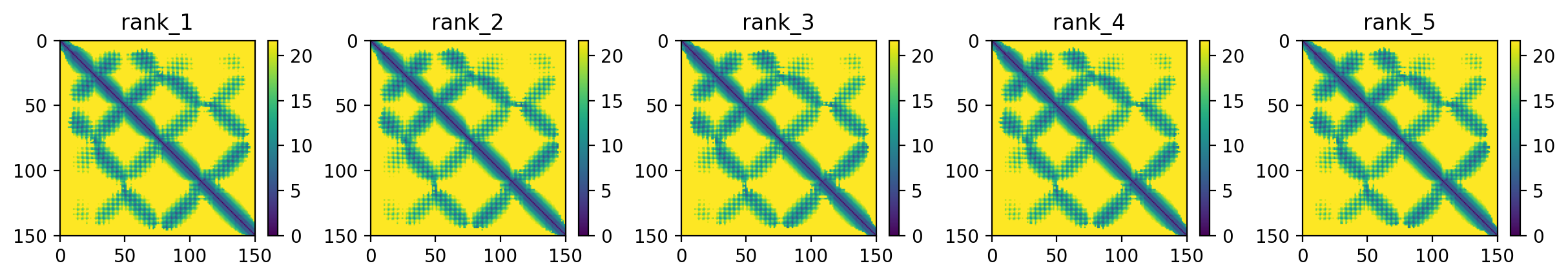

### predicted_distogram.png

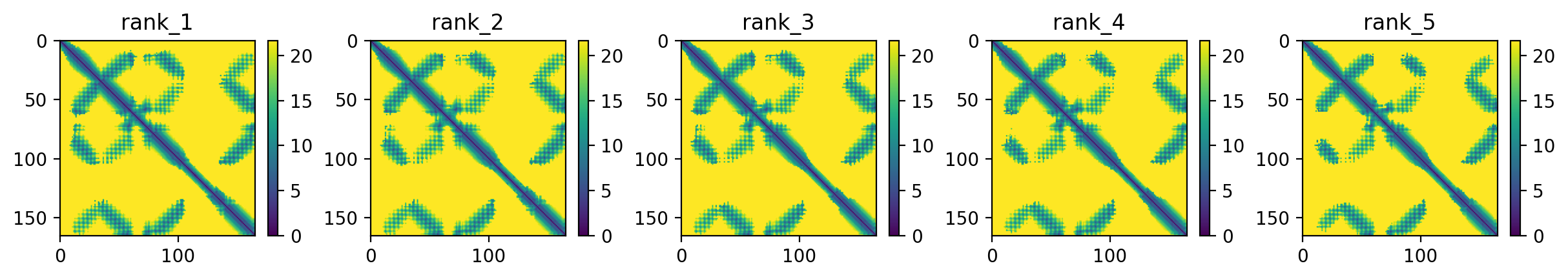

### predicted_distogram.png

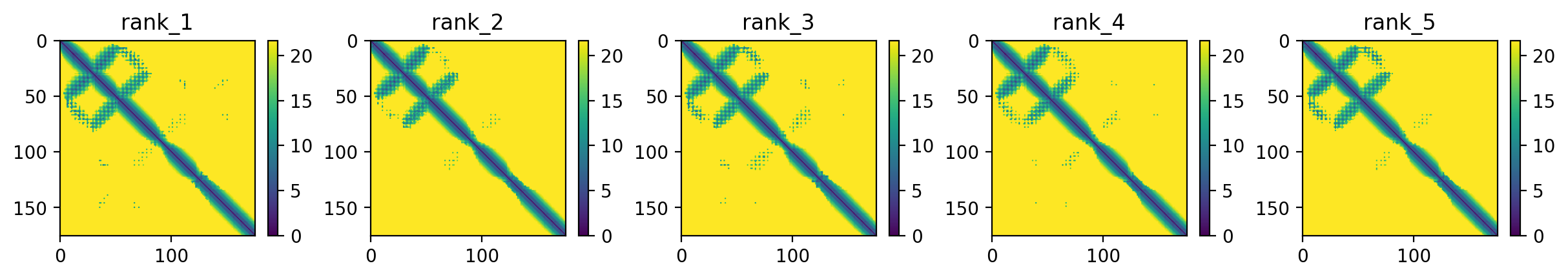

### predicted_distogram.png

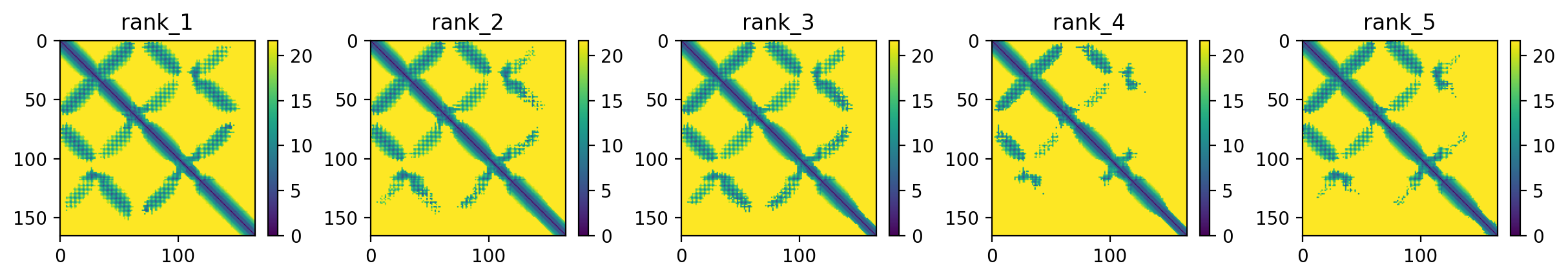

### predicted_distogram.png

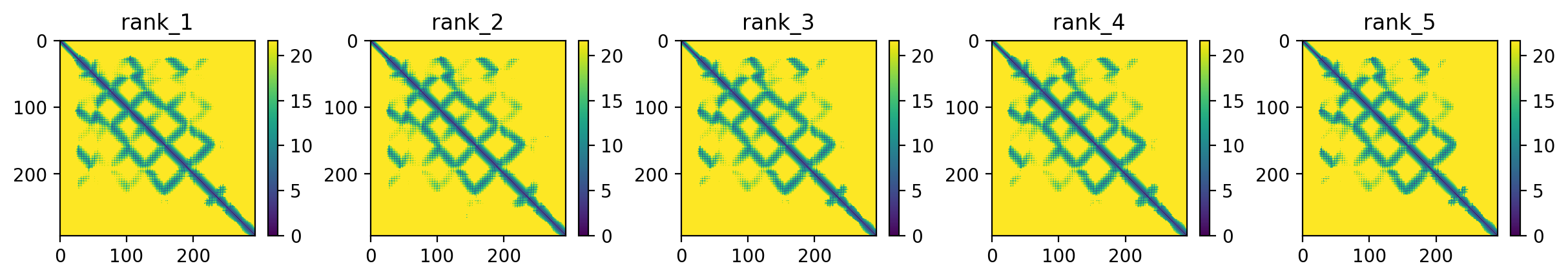

### predicted_distogram.png

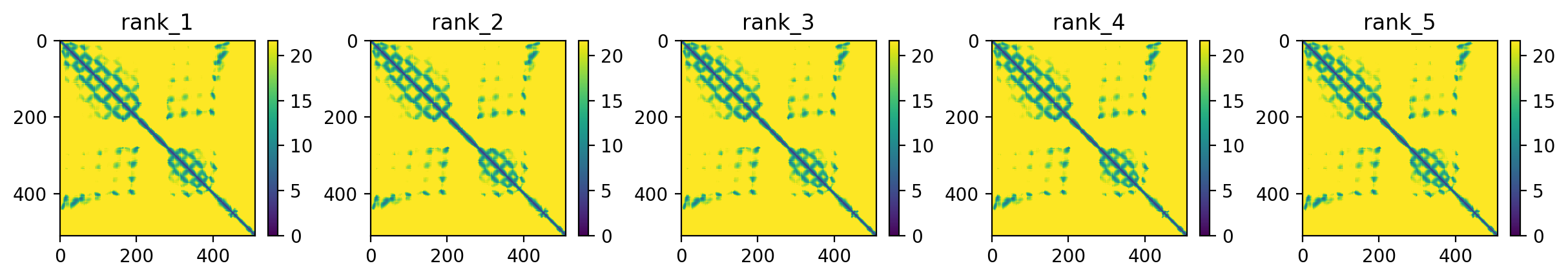

### predicted_distogram.png

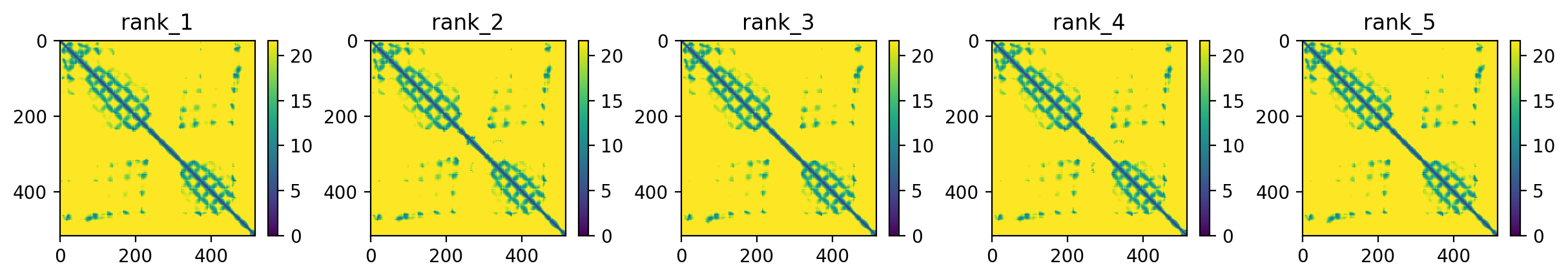

### predicted_distogram.png

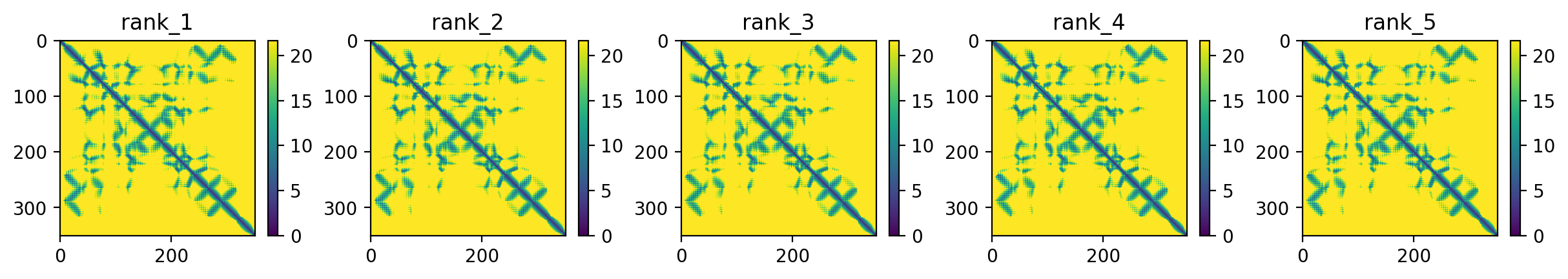

### predicted_LDDT.png

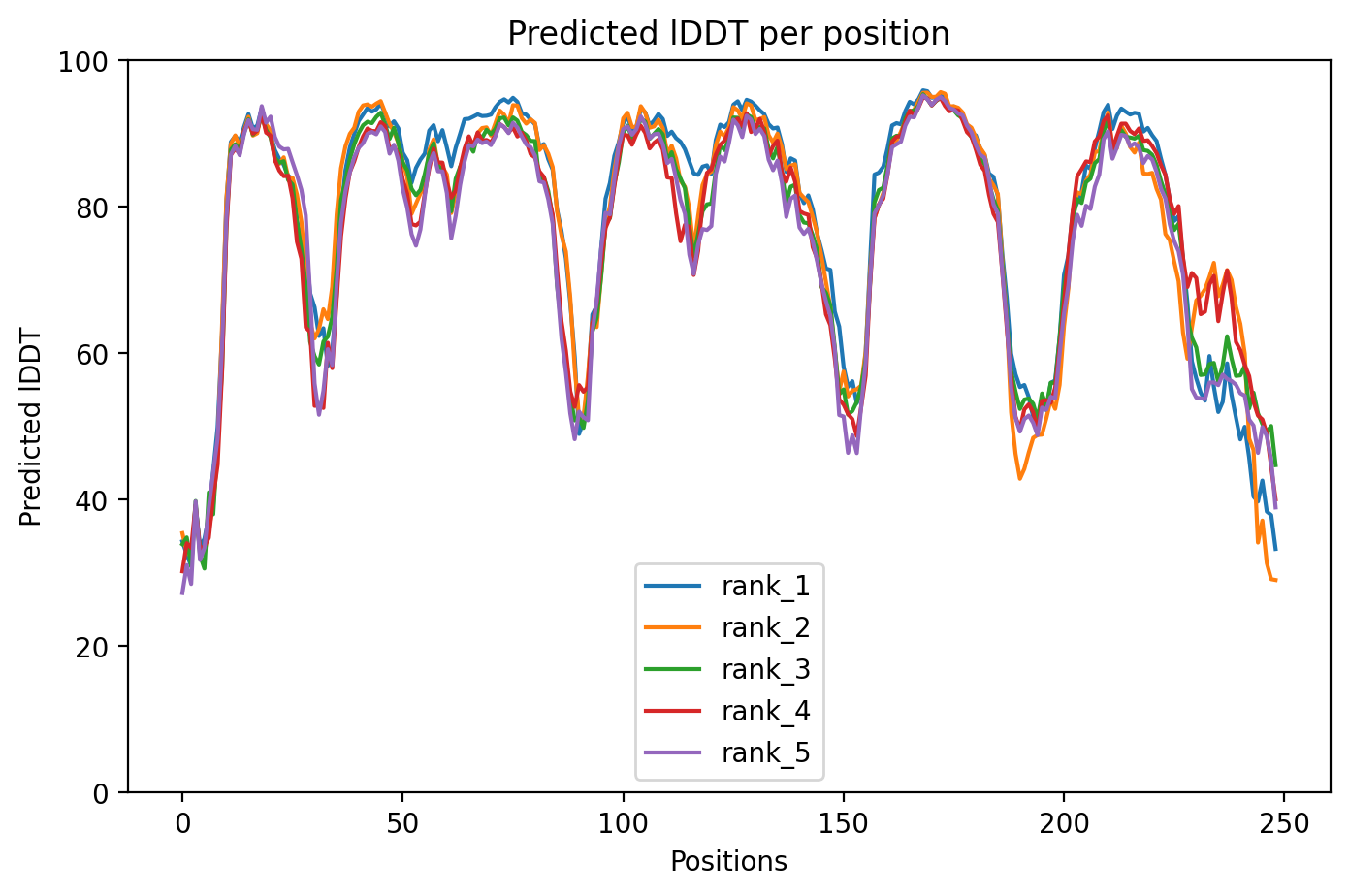

### predicted_LDDT.png

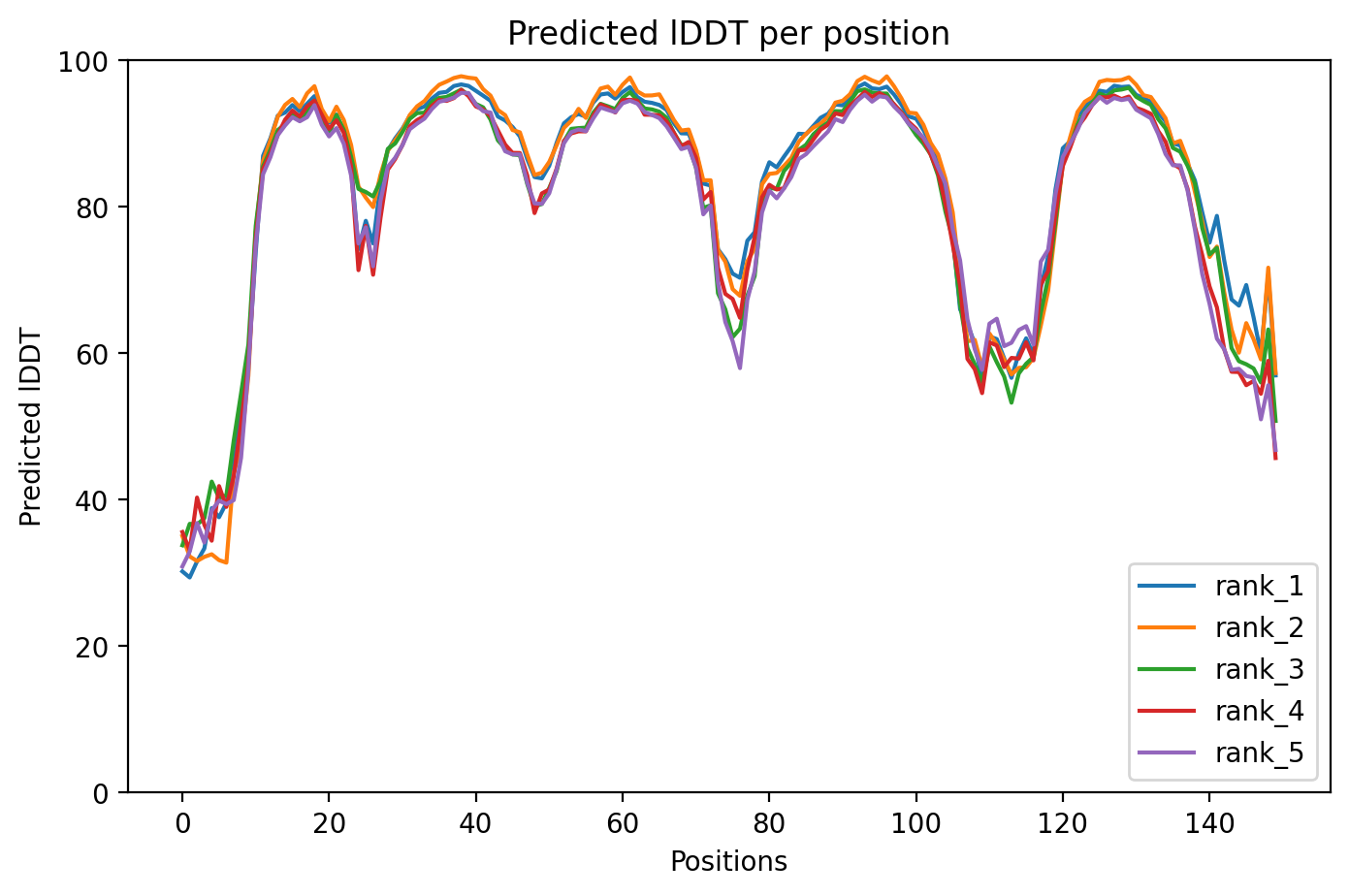

### predicted_LDDT.png

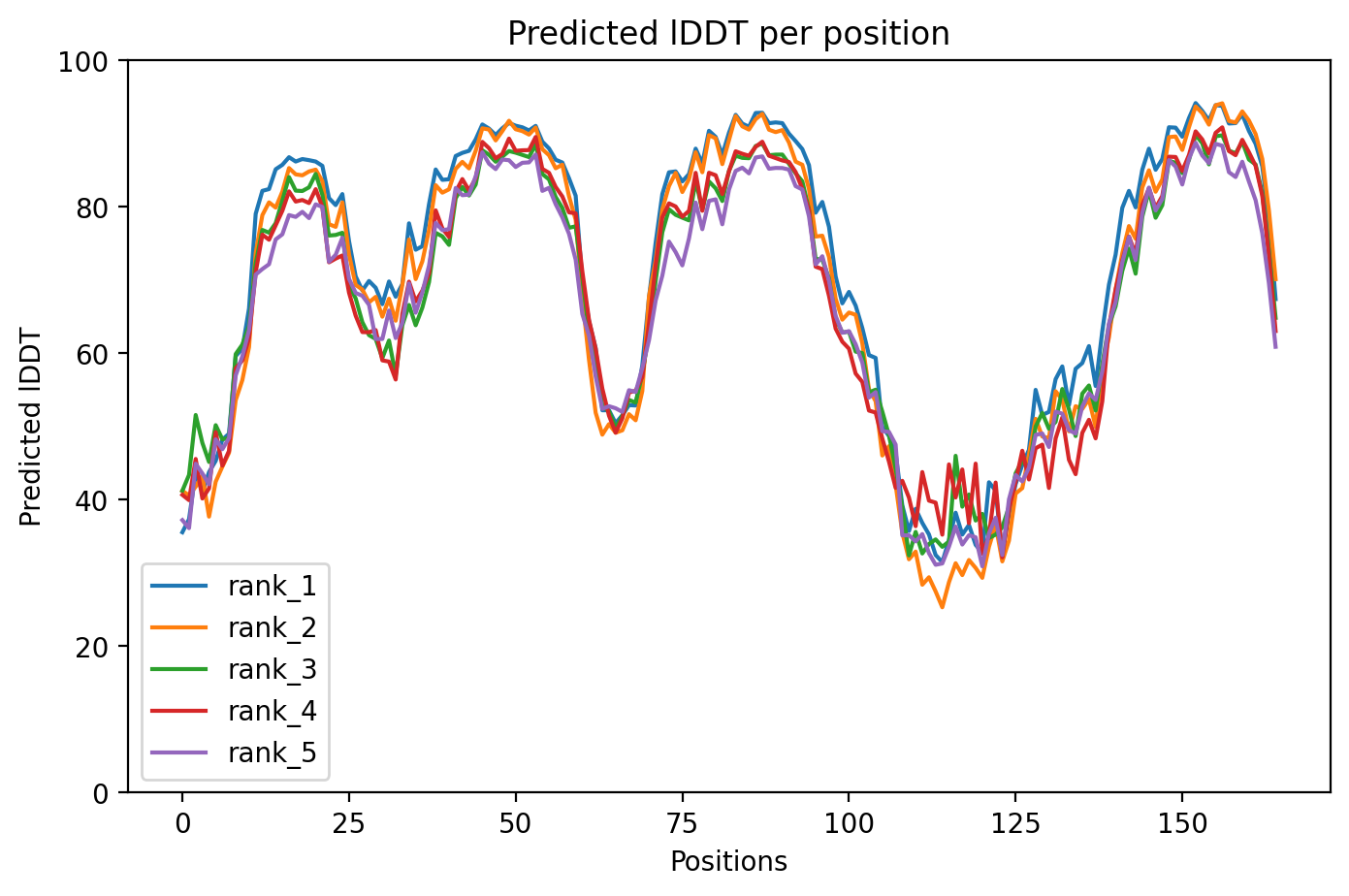

### predicted_LDDT.png

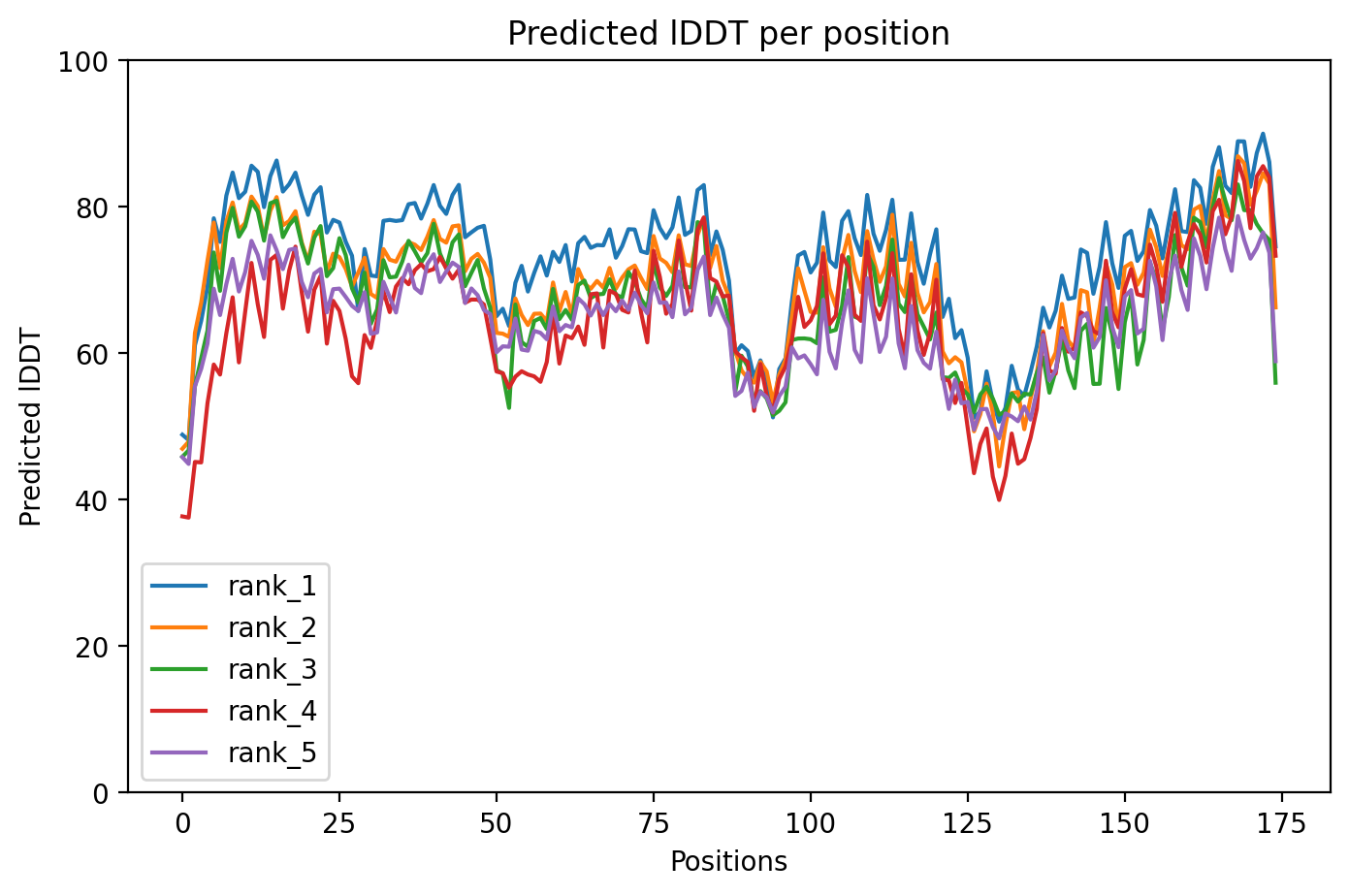

### predicted_LDDT.png

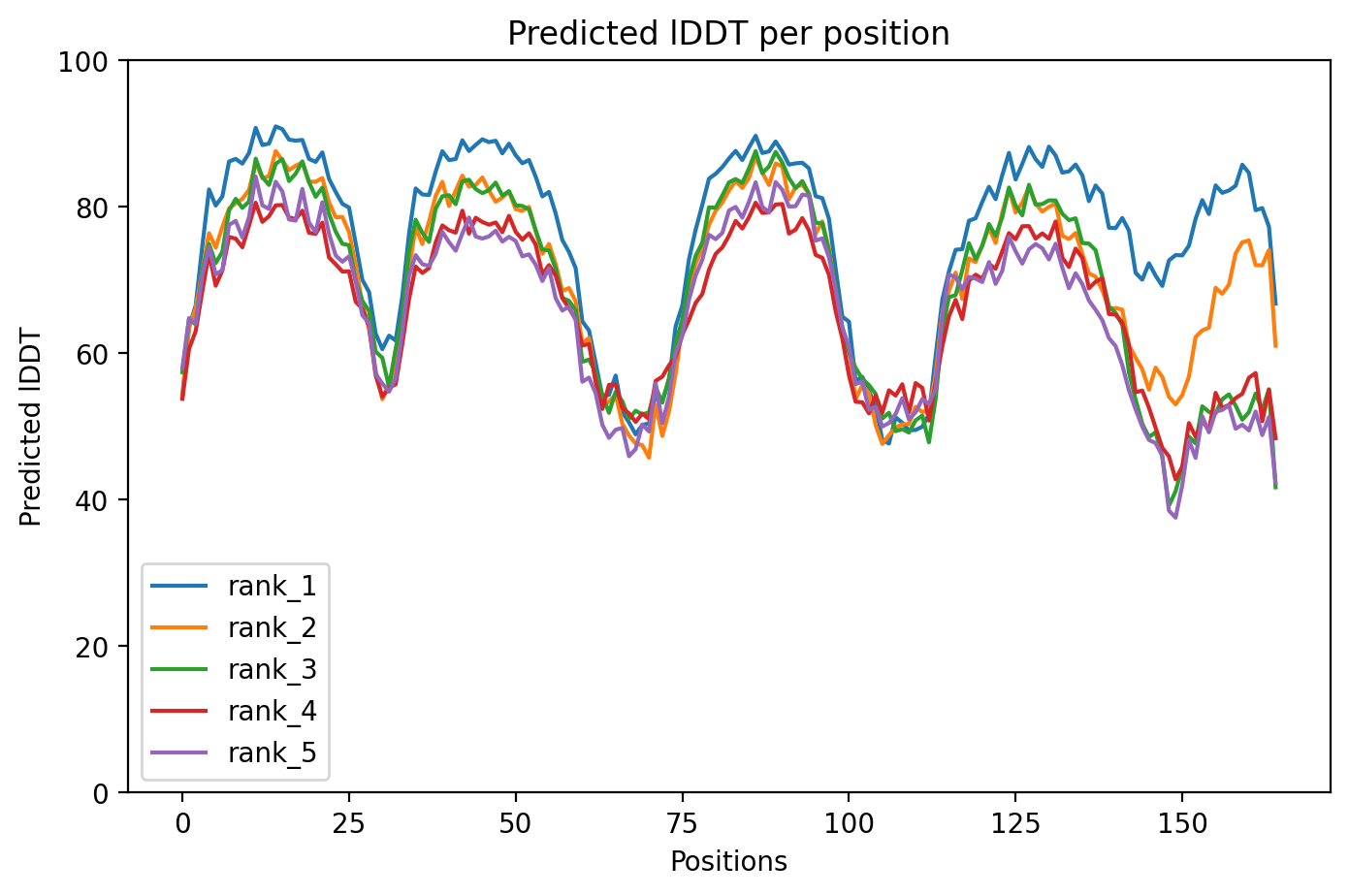

### predicted_LDDT.png

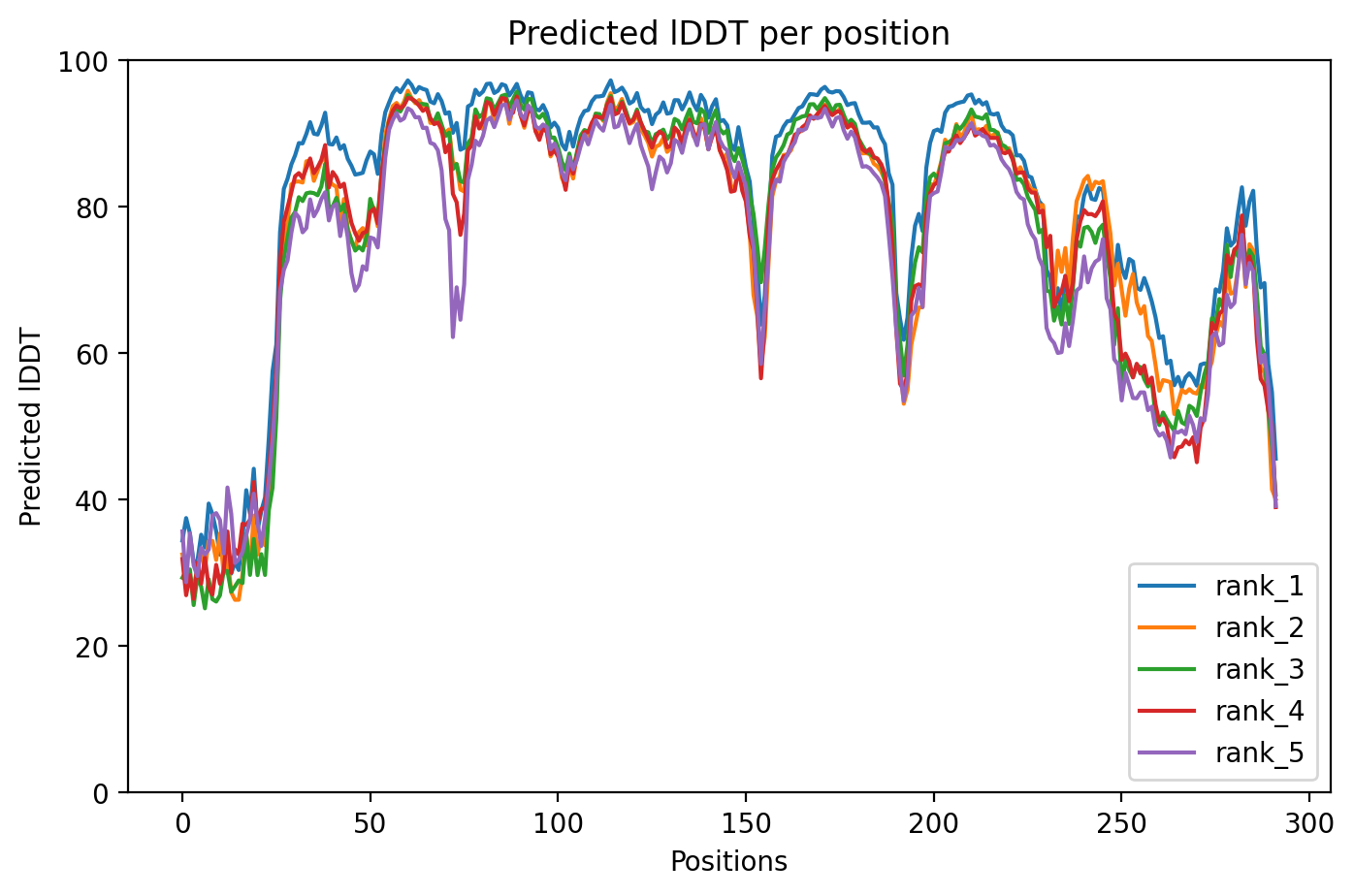

### predicted_LDDT.png

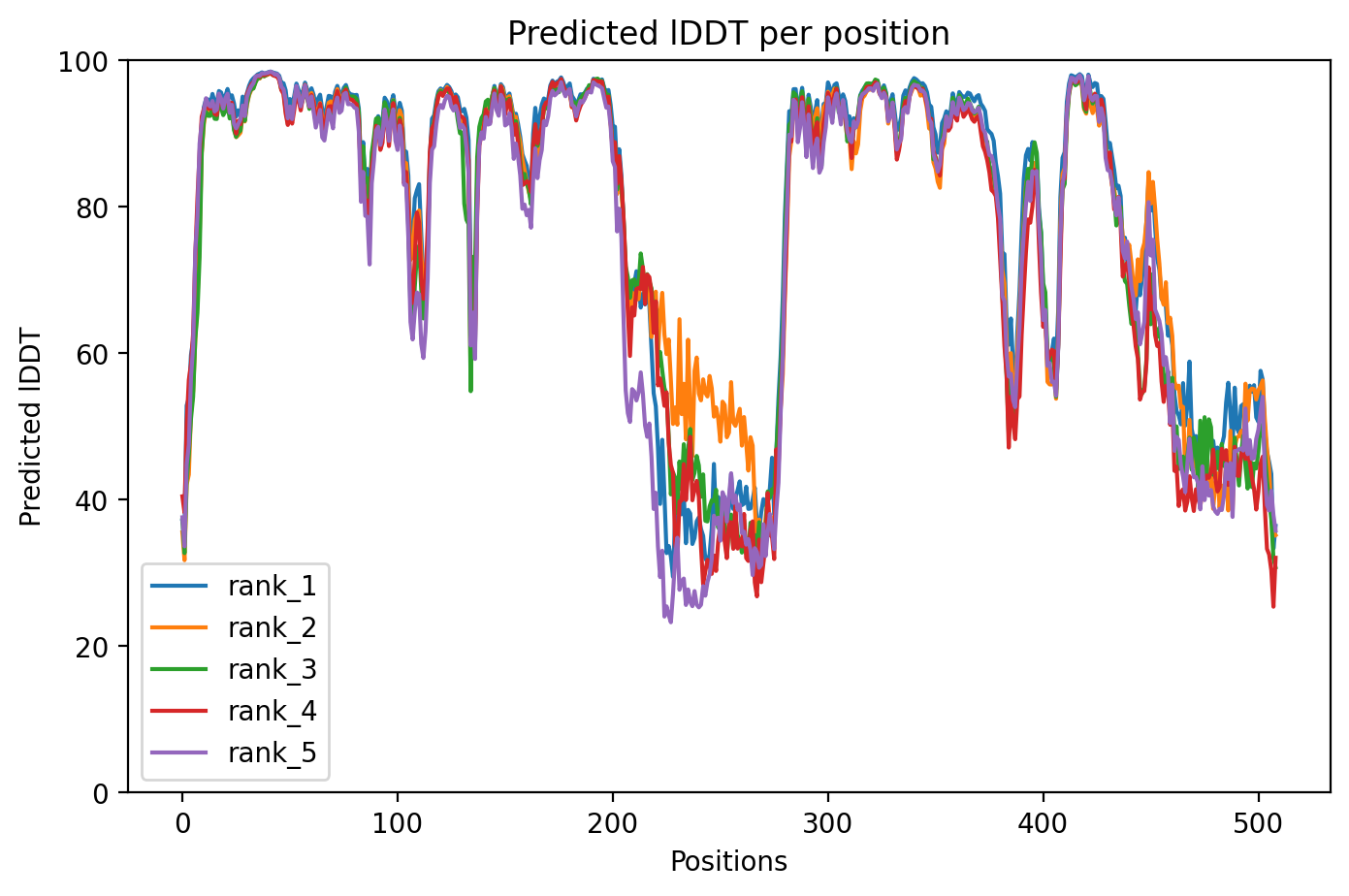

### predicted_LDDT.png

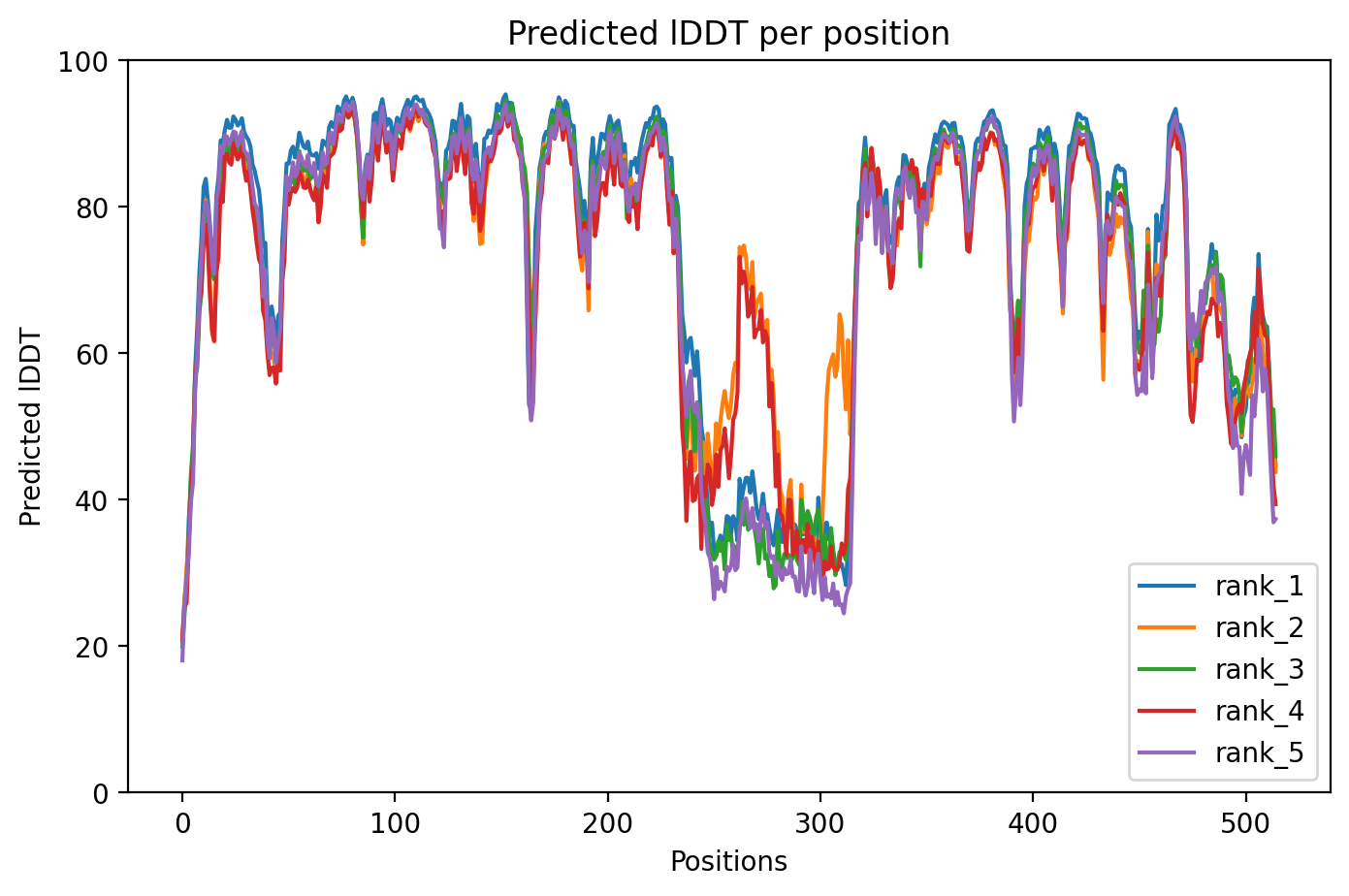

### predicted_LDDT.png

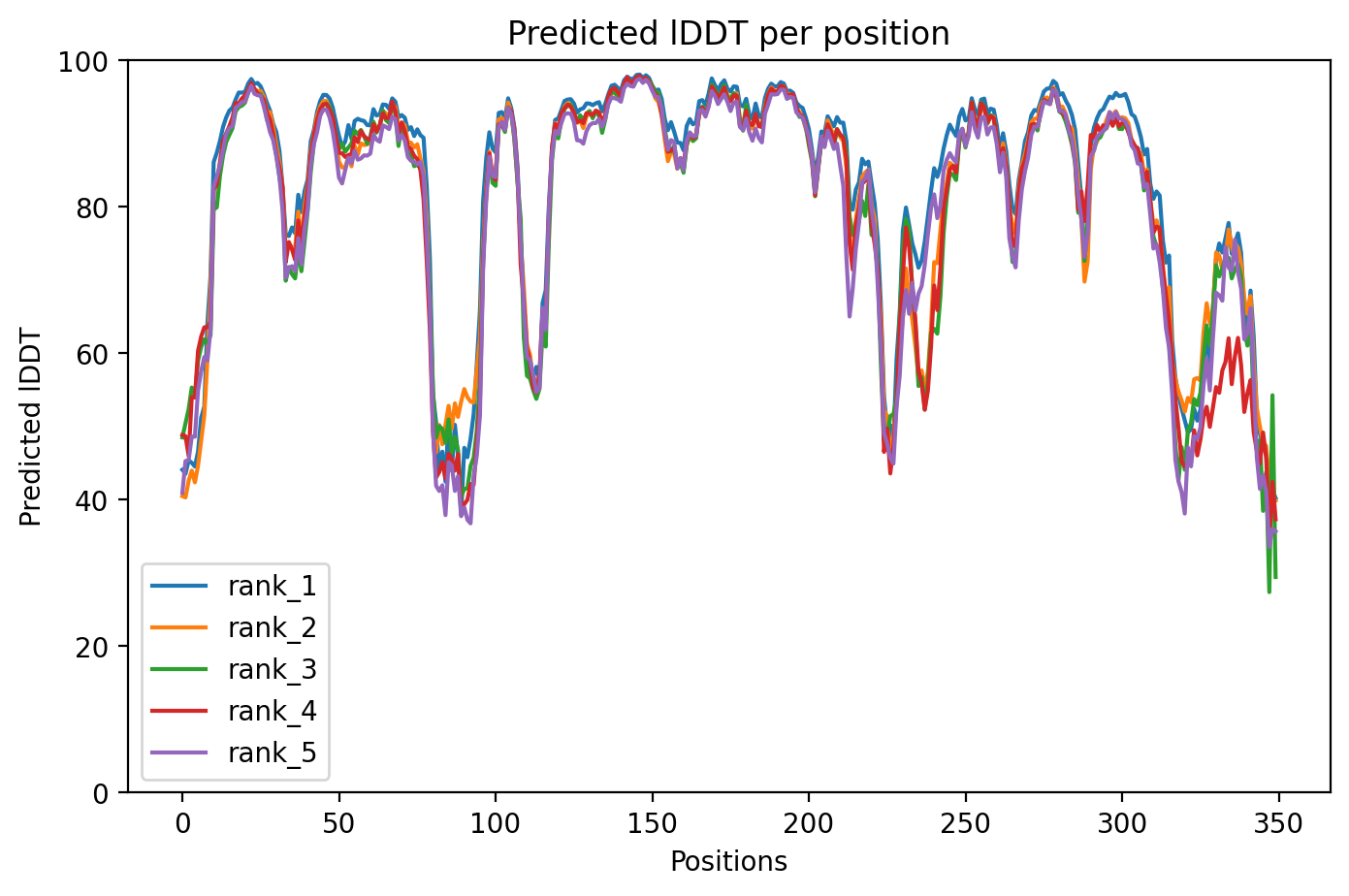

### rank_1_model_3_ptm_seed_0.png

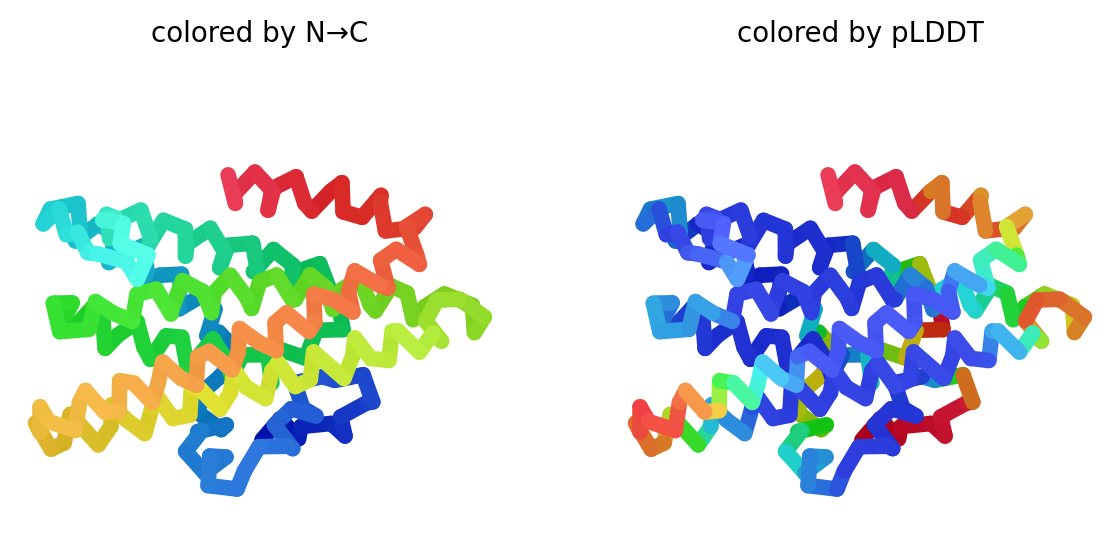

### rank_1_model_3_ptm_seed_0.png

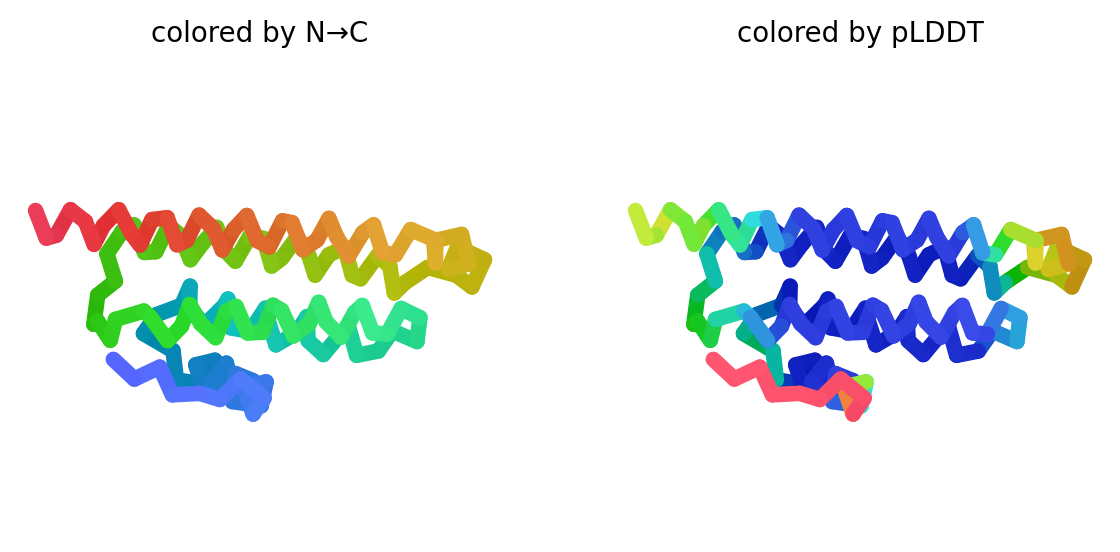

### rank_1_model_3_ptm_seed_0.png

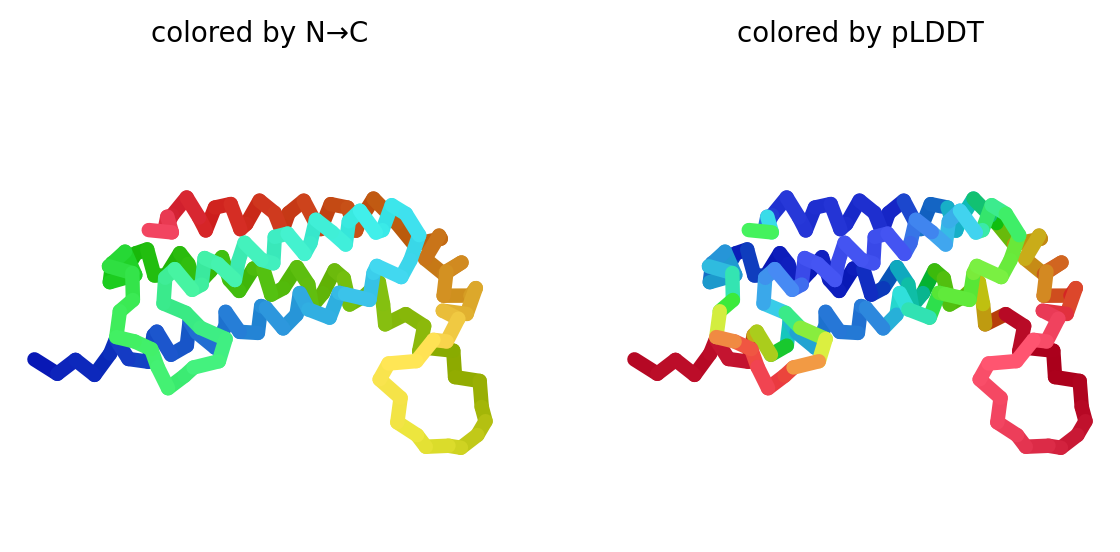

### rank_1_model_3_ptm_seed_0.png

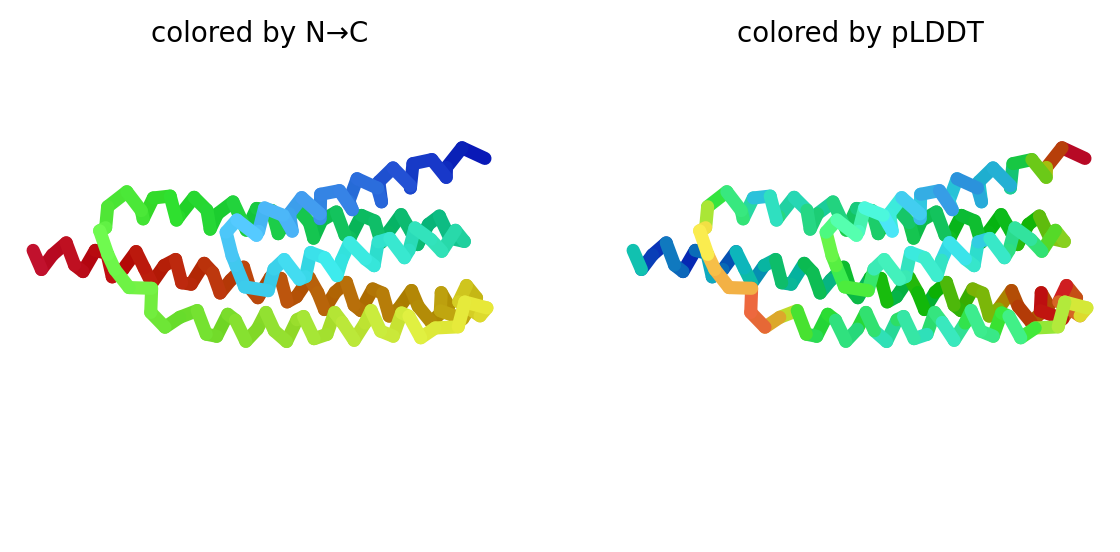

### rank_1_model_3_ptm_seed_0.png

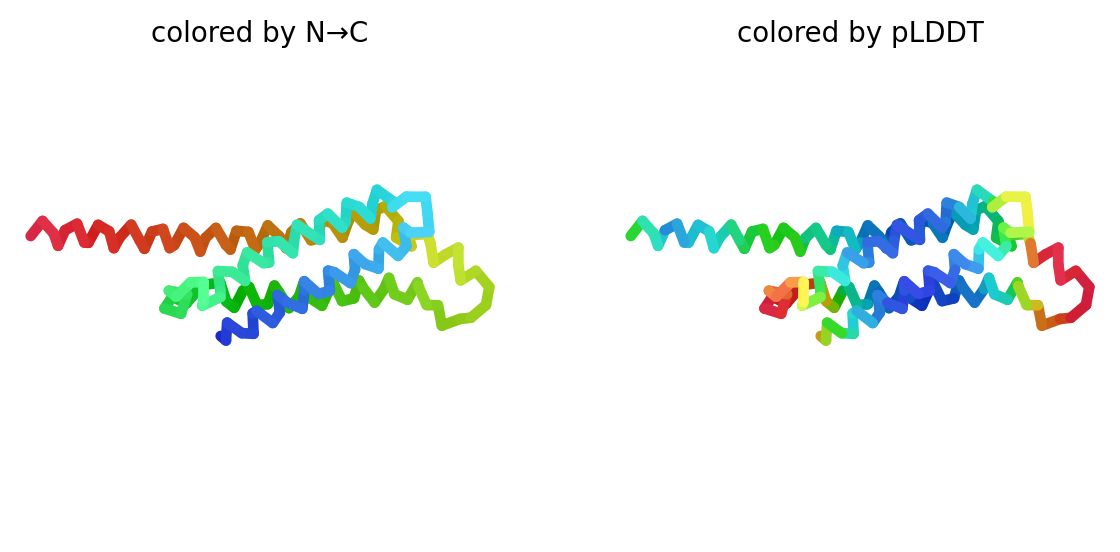

### rank_1_model_3_ptm_seed_0.png

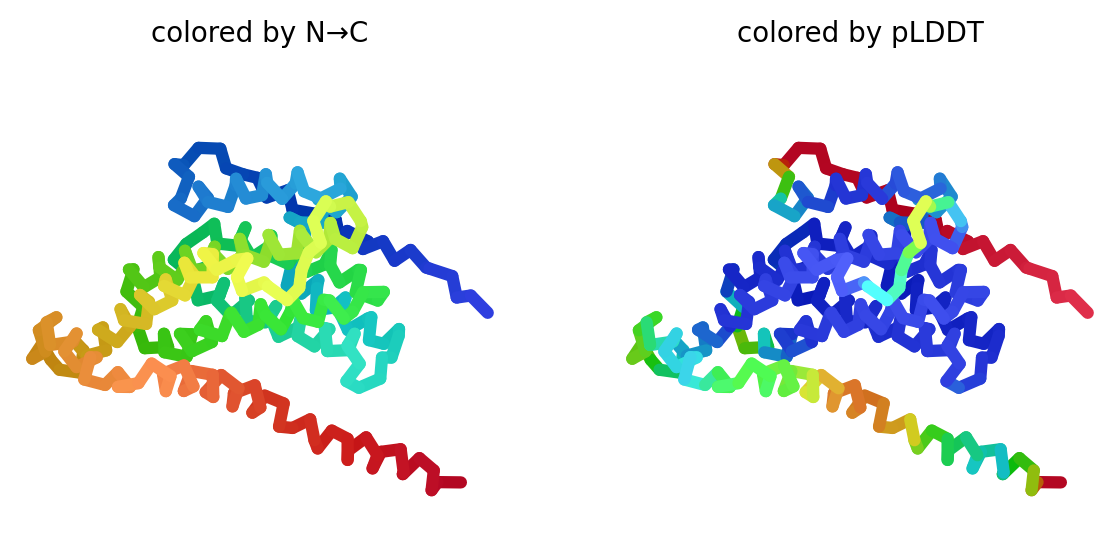

### rank_1_model_3_ptm_seed_0.png

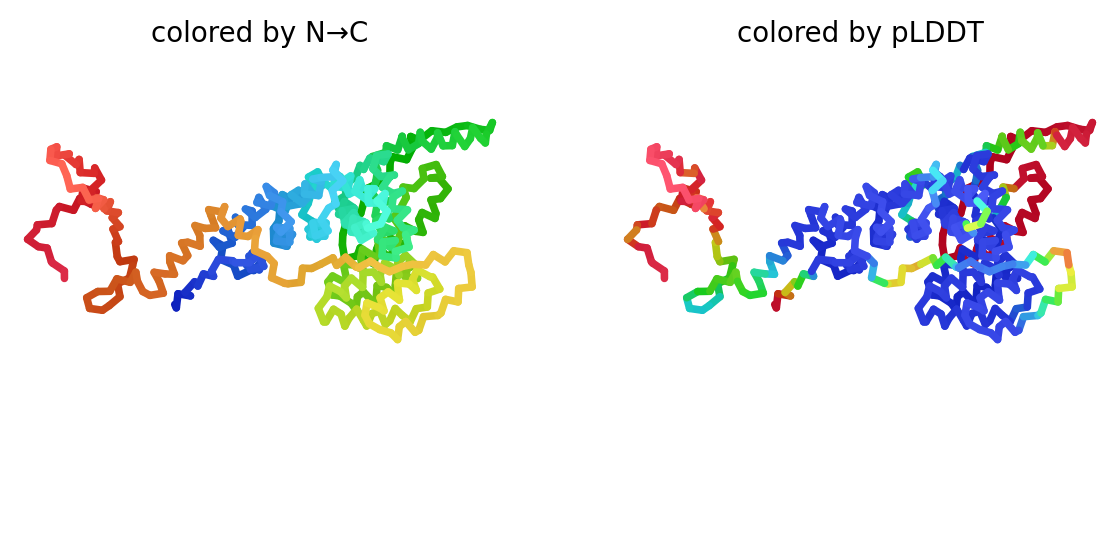

### rank_1_model_3_ptm_seed_0.png

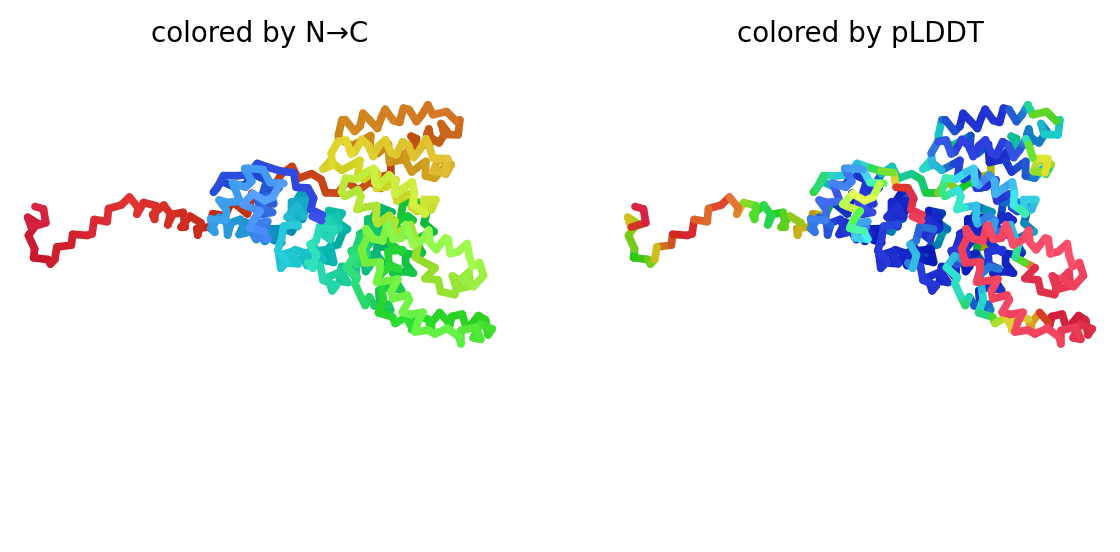

### rank_1_model_3_ptm_seed_0.png

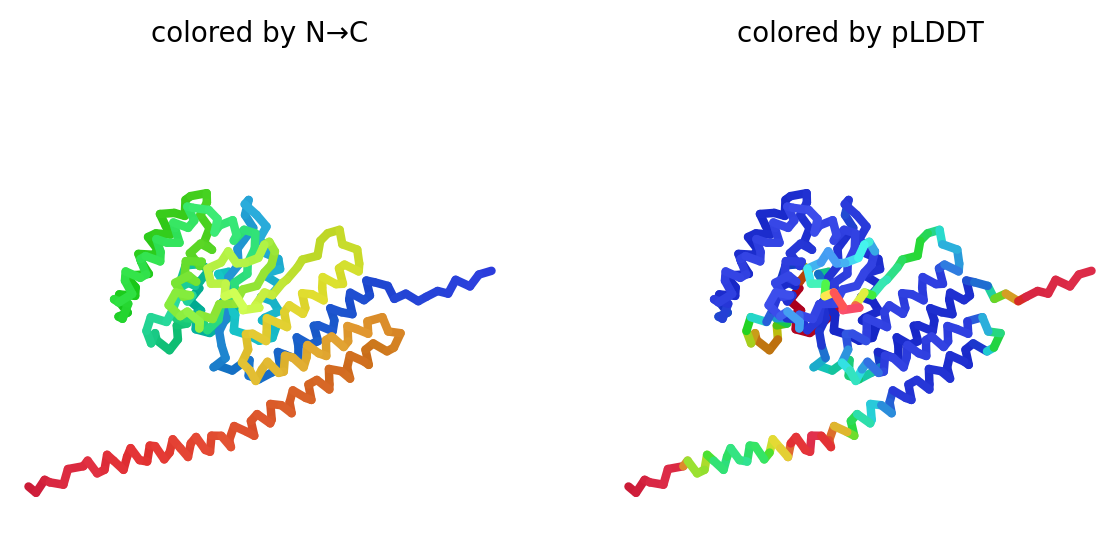

### rank_2_model_2_ptm_seed_0.png

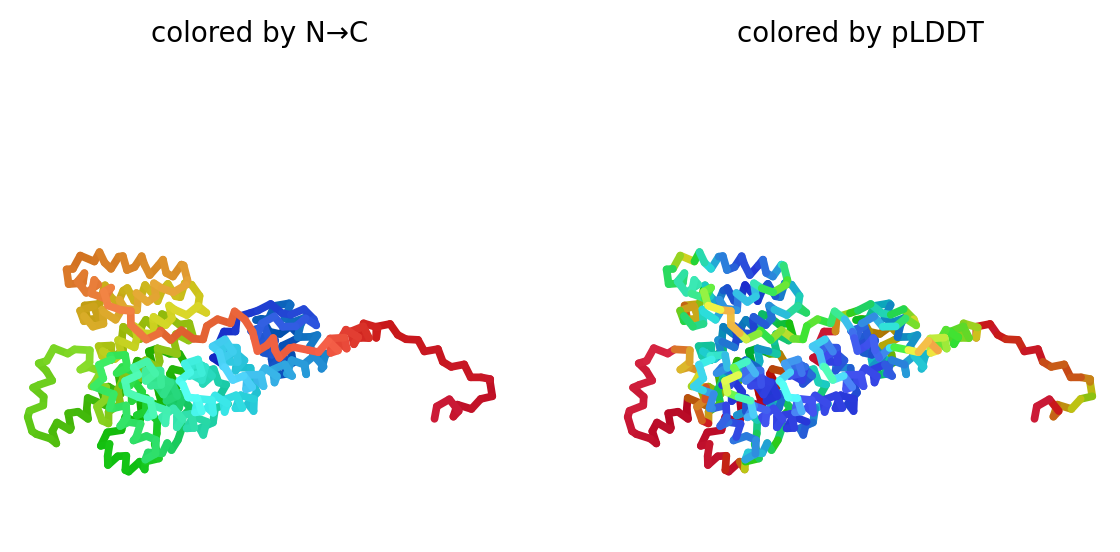

### rank_2_model_4_ptm_seed_0.png

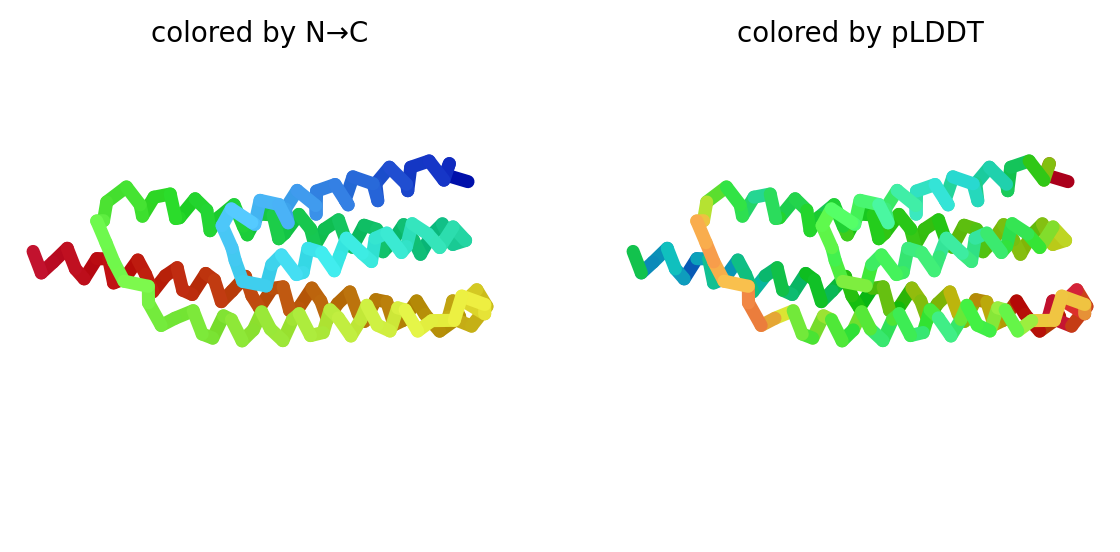

### rank_2_model_4_ptm_seed_0.png

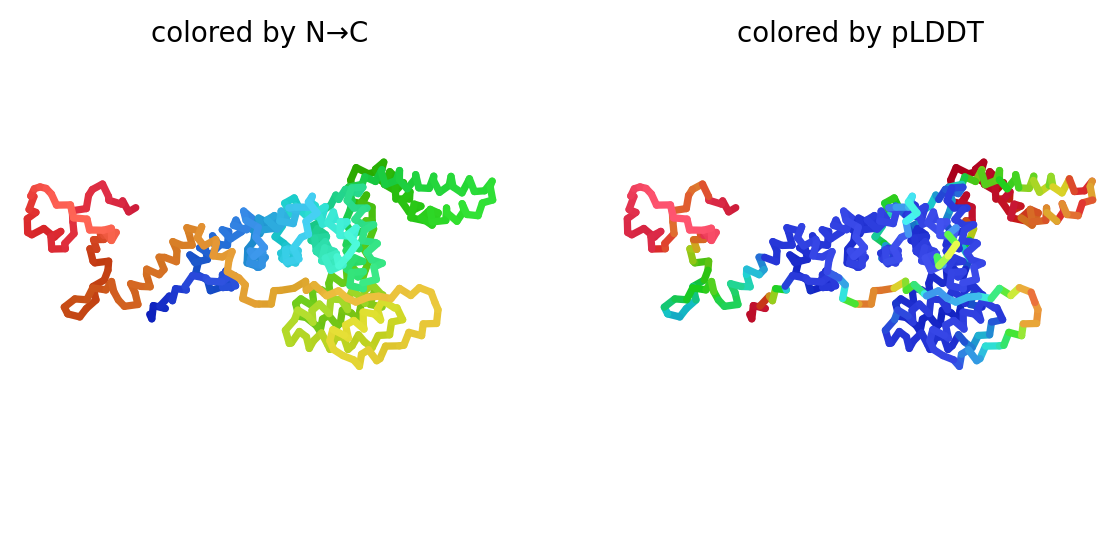

### rank_2_model_4_ptm_seed_0.png

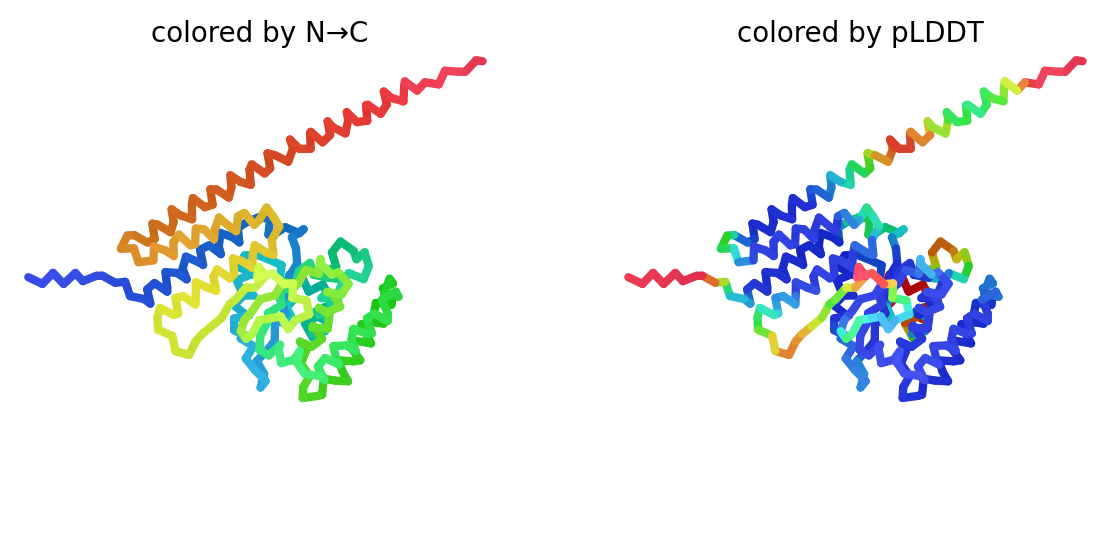

### rank_2_model_5_ptm_seed_0.png

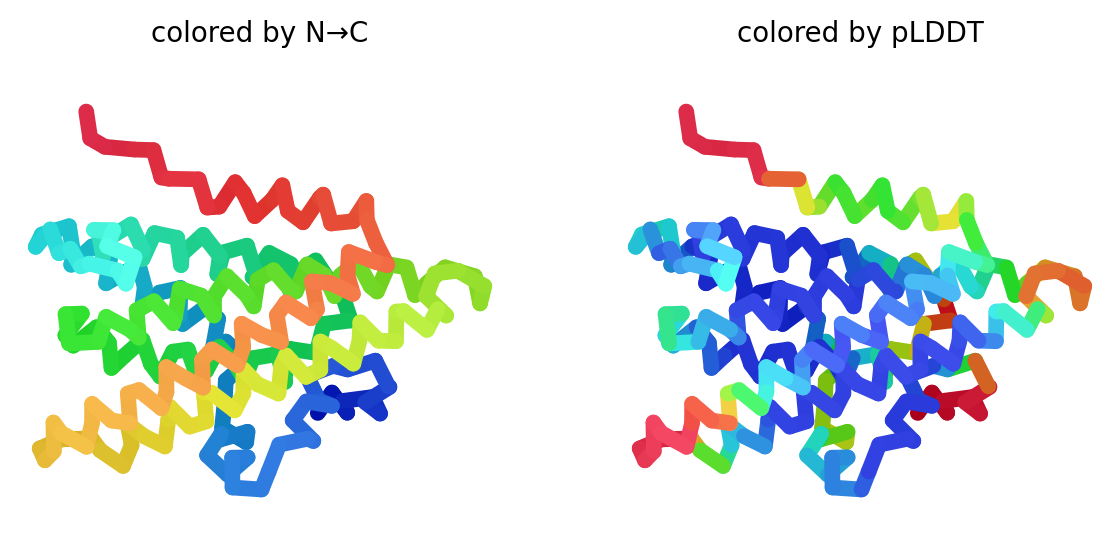

### rank_2_model_5_ptm_seed_0.png

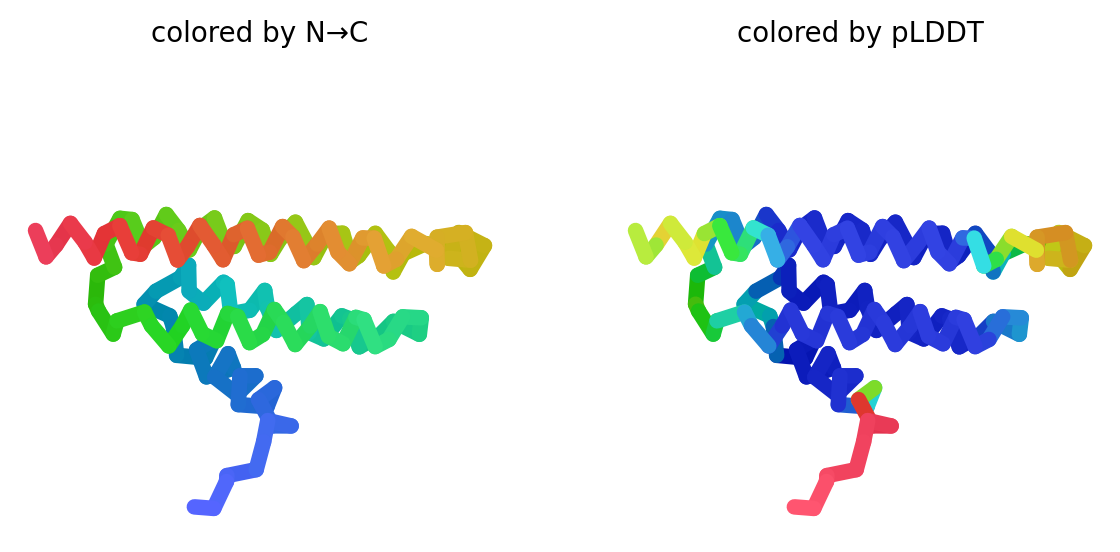

### rank_2_model_5_ptm_seed_0.png

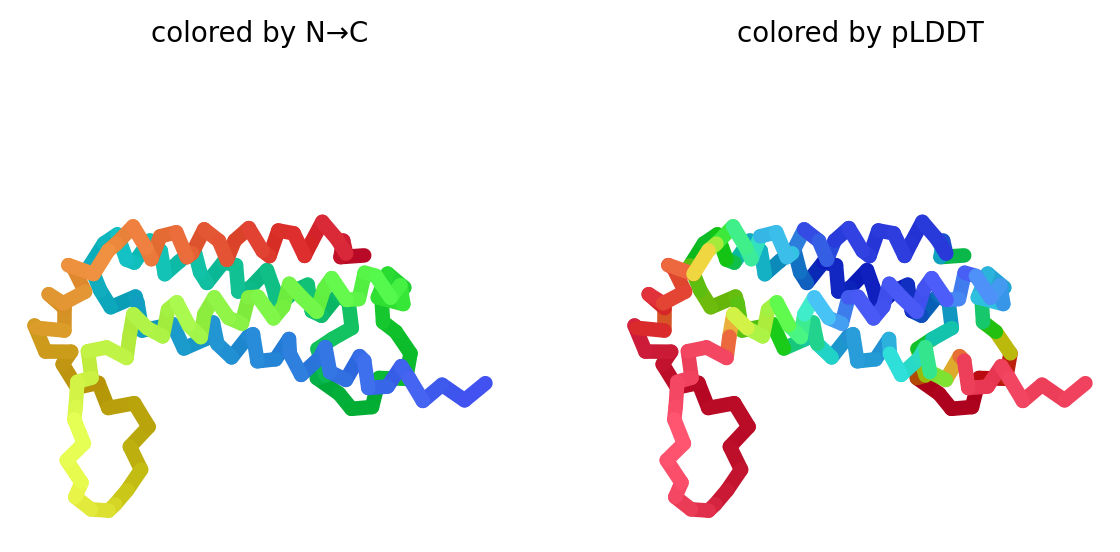

### rank_2_model_5_ptm_seed_0.png

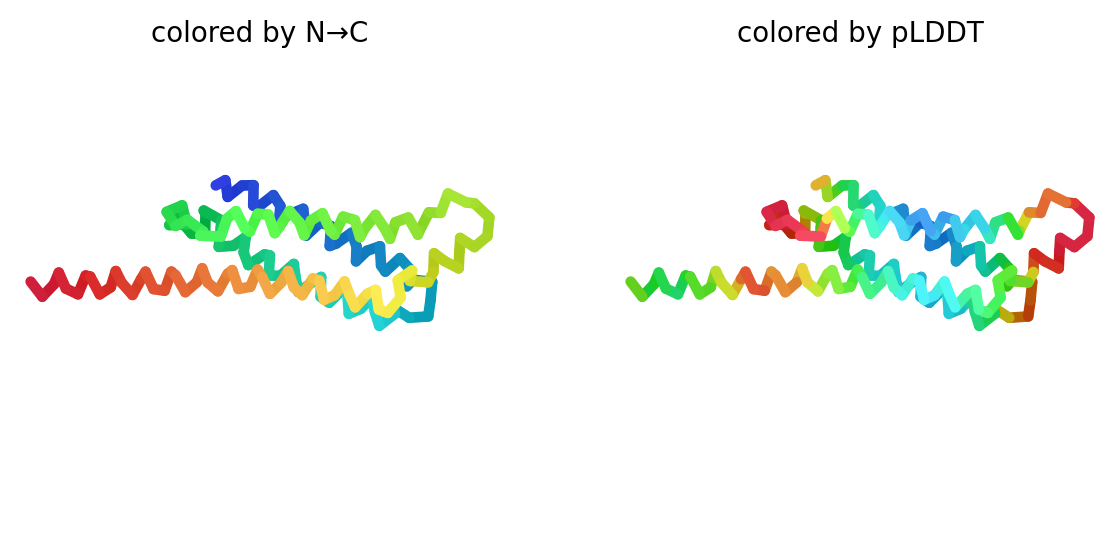

### rank_2_model_5_ptm_seed_0.png

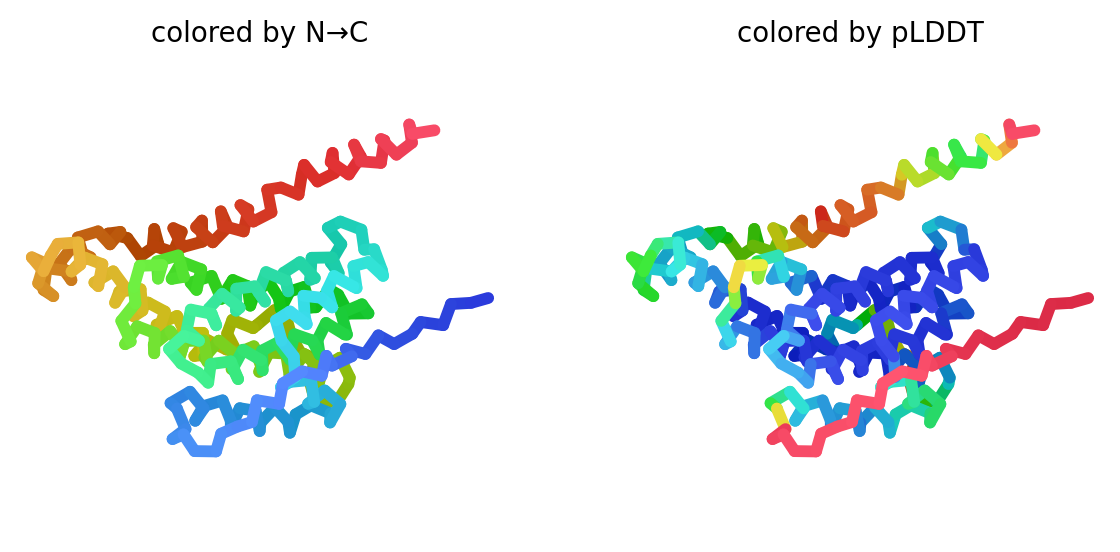

### rank_3_model_1_ptm_seed_0.png

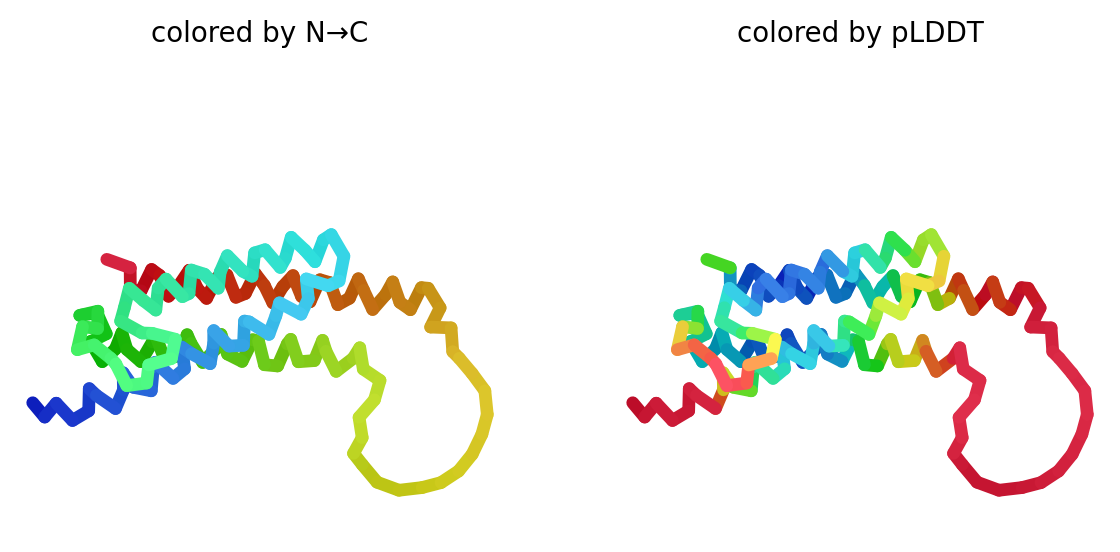

### rank_3_model_1_ptm_seed_0.png

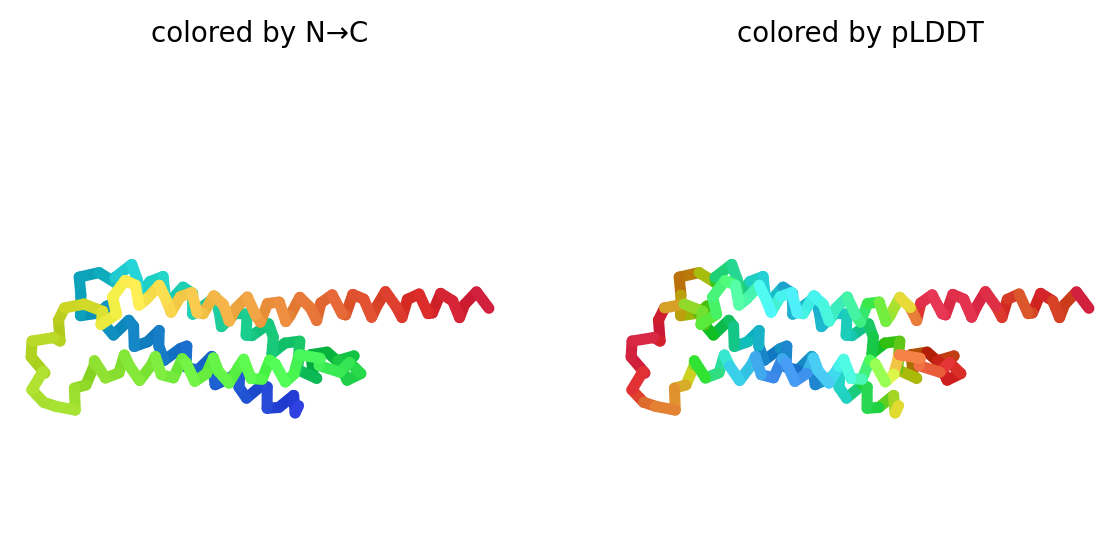

### rank_3_model_1_ptm_seed_0.png

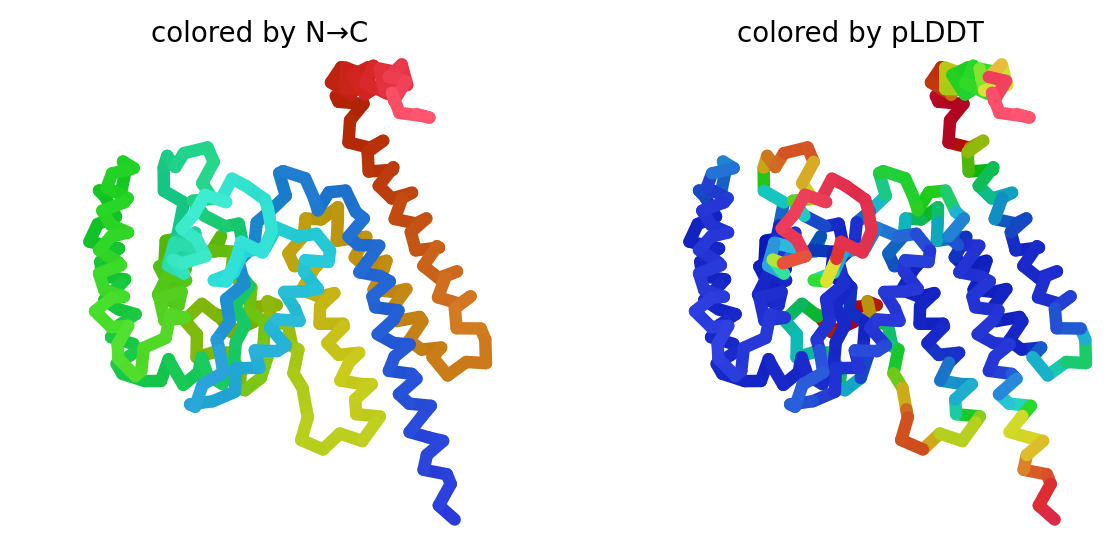

### rank_3_model_2_ptm_seed_0.png

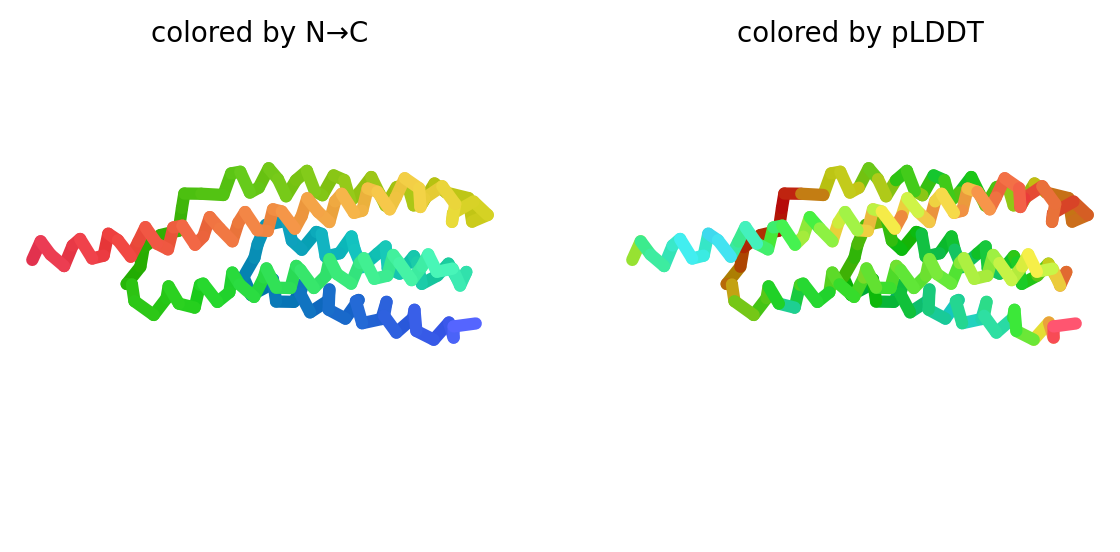

### rank_3_model_2_ptm_seed_0.png

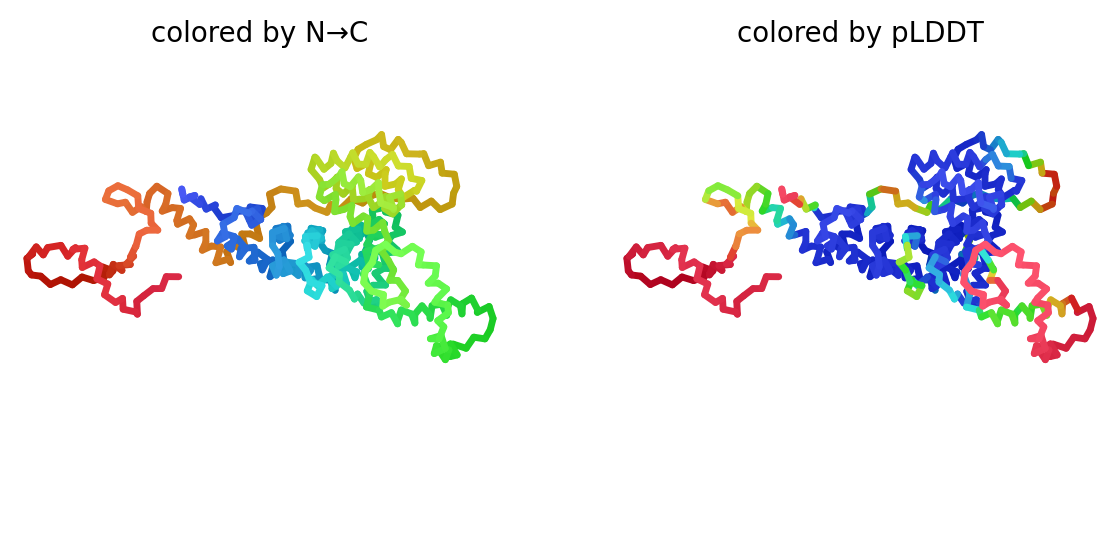

### rank_3_model_4_ptm_seed_0.png

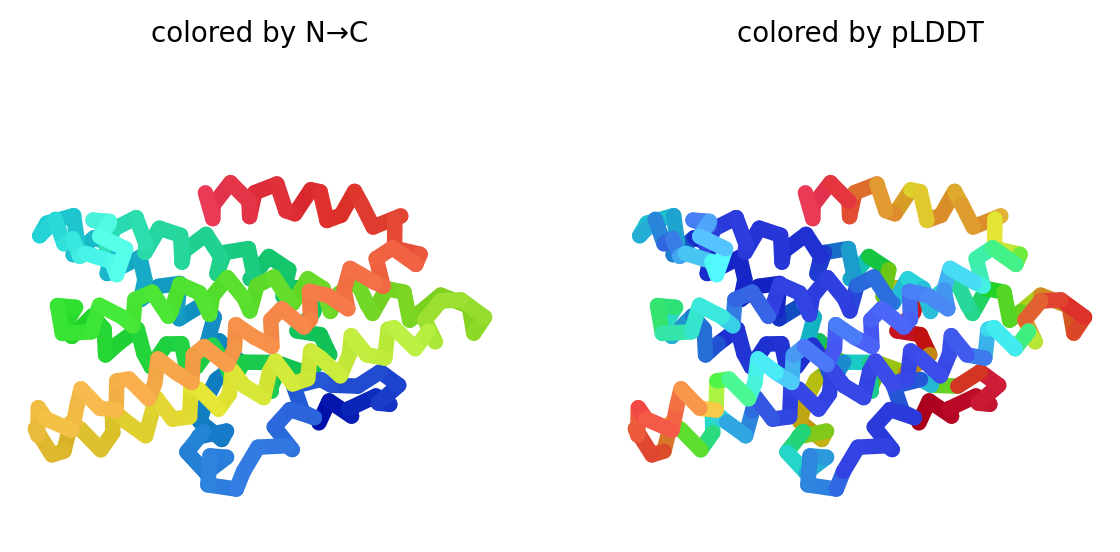

### rank_3_model_4_ptm_seed_0.png

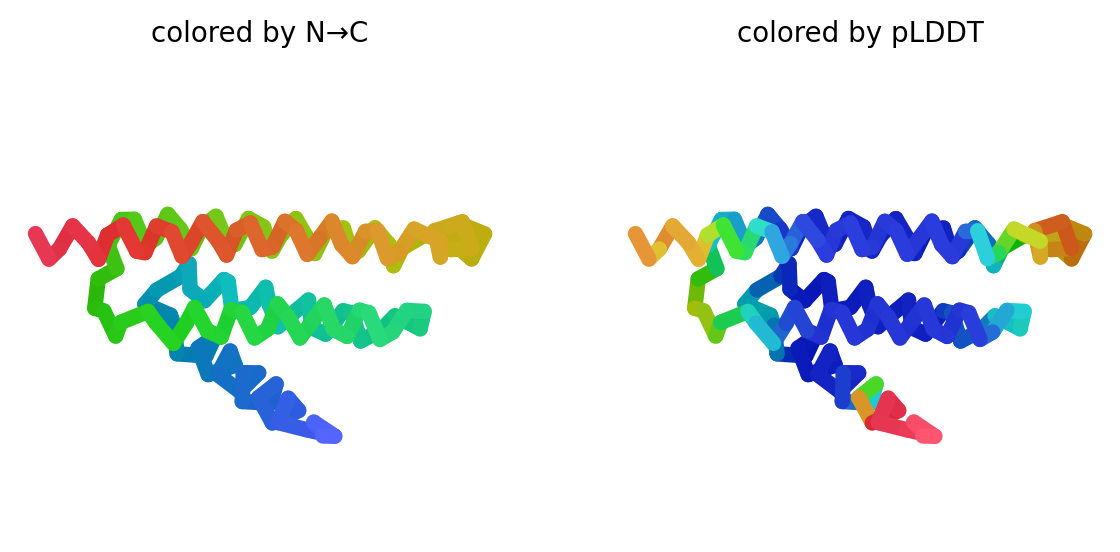

### rank_3_model_4_ptm_seed_0.png

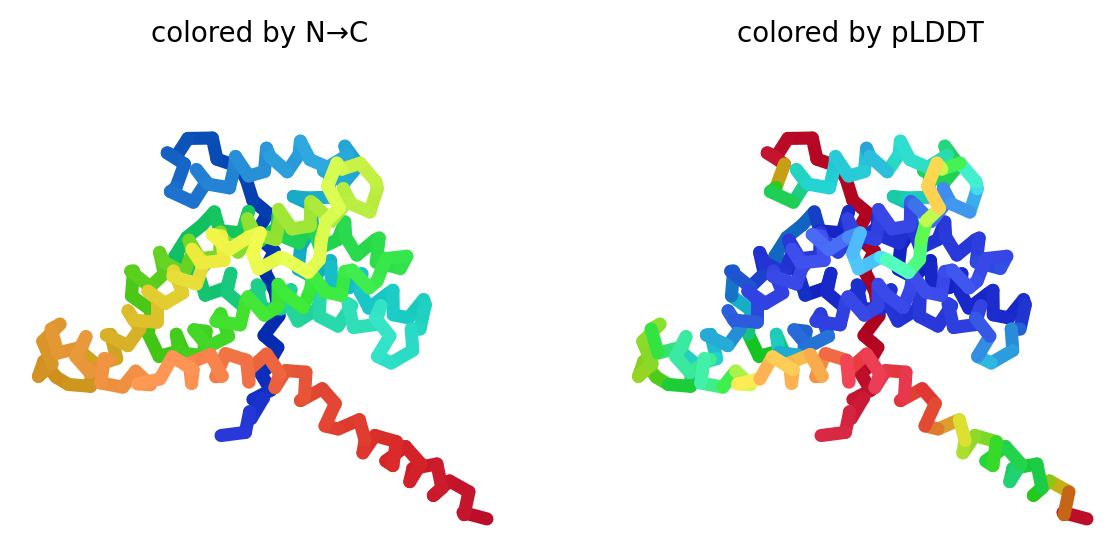

### rank_3_model_4_ptm_seed_0.png

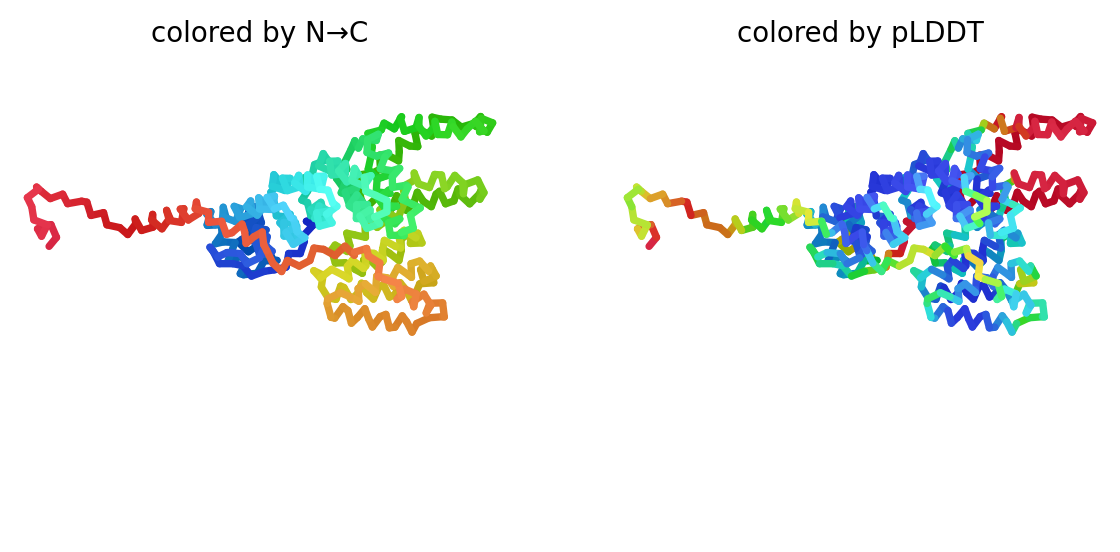

### rank_4_model_1_ptm_seed_0.png

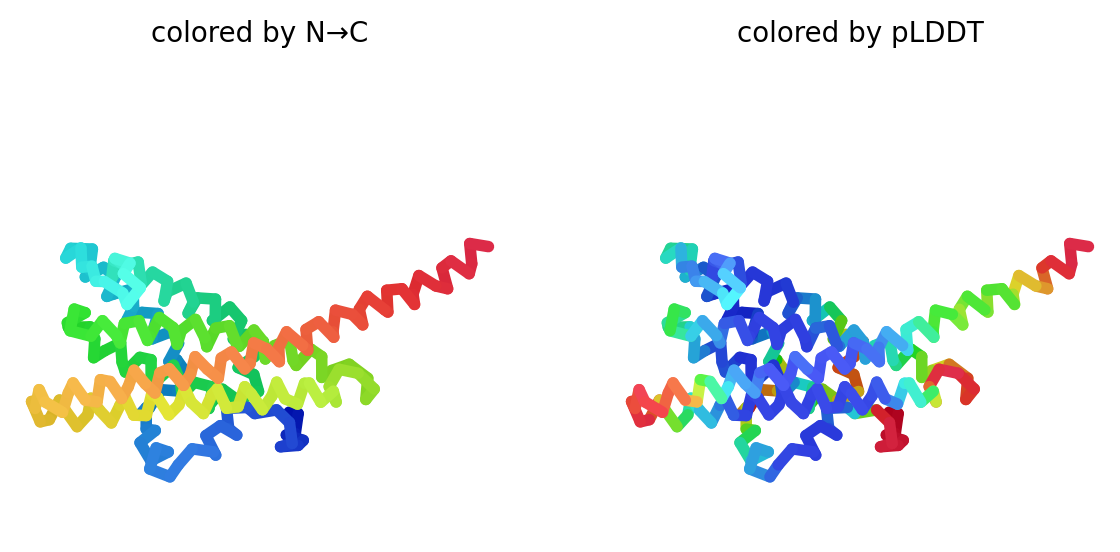

### rank_4_model_1_ptm_seed_0.png

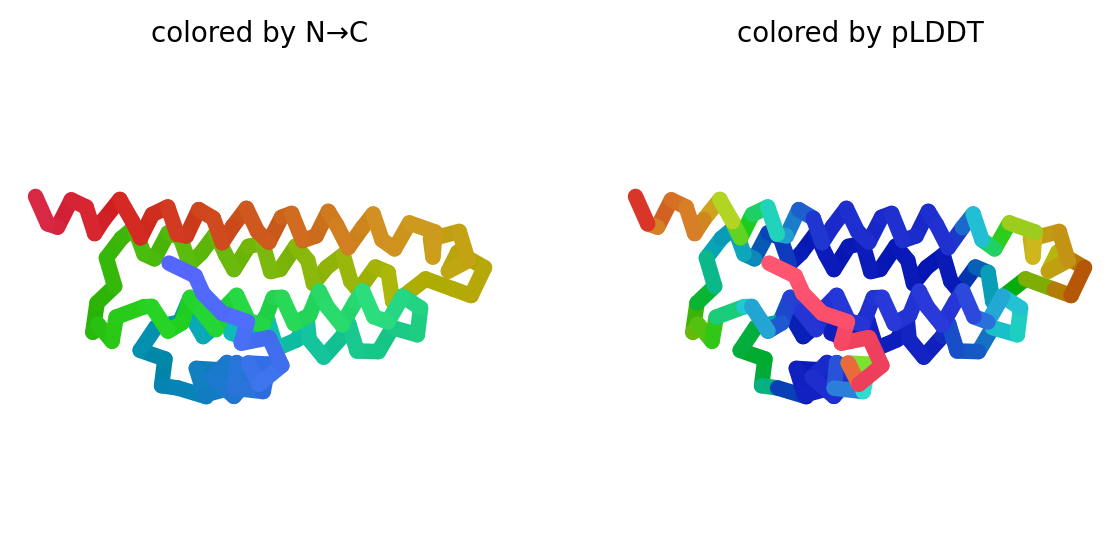

### rank_4_model_1_ptm_seed_0.png

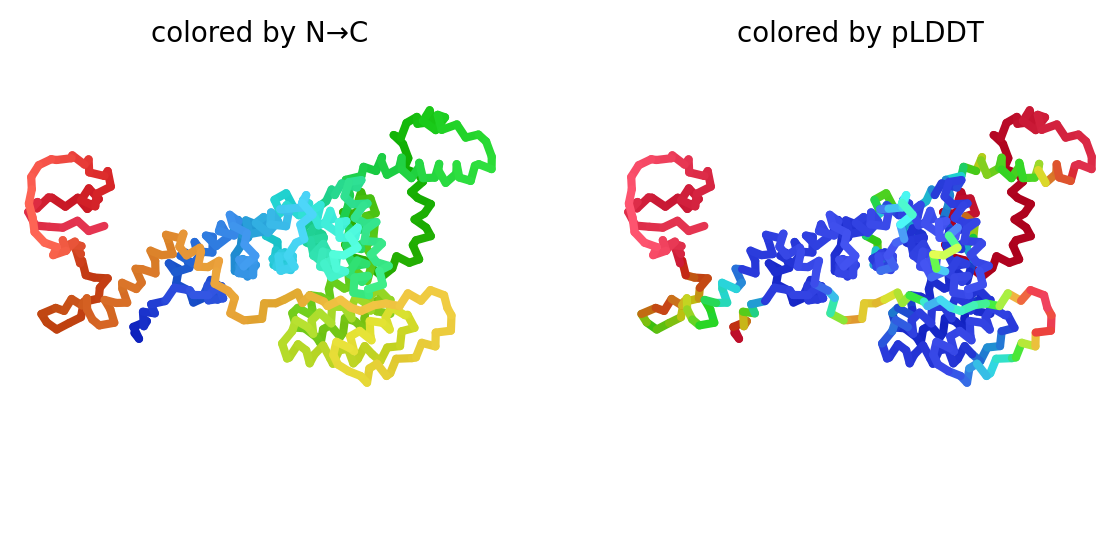

### rank_4_model_1_ptm_seed_0.png

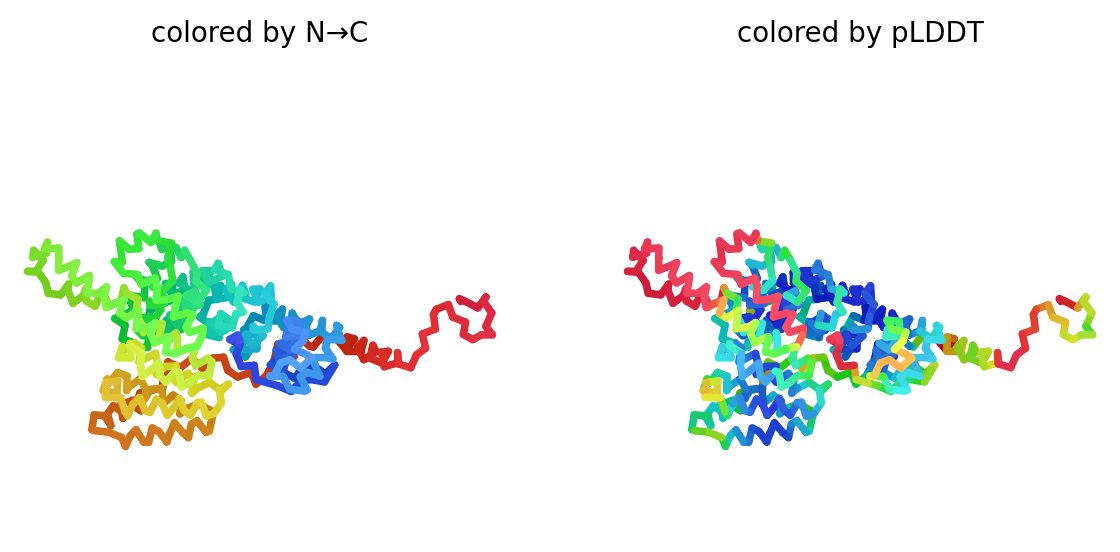

### rank_4_model_2_ptm_seed_0.png

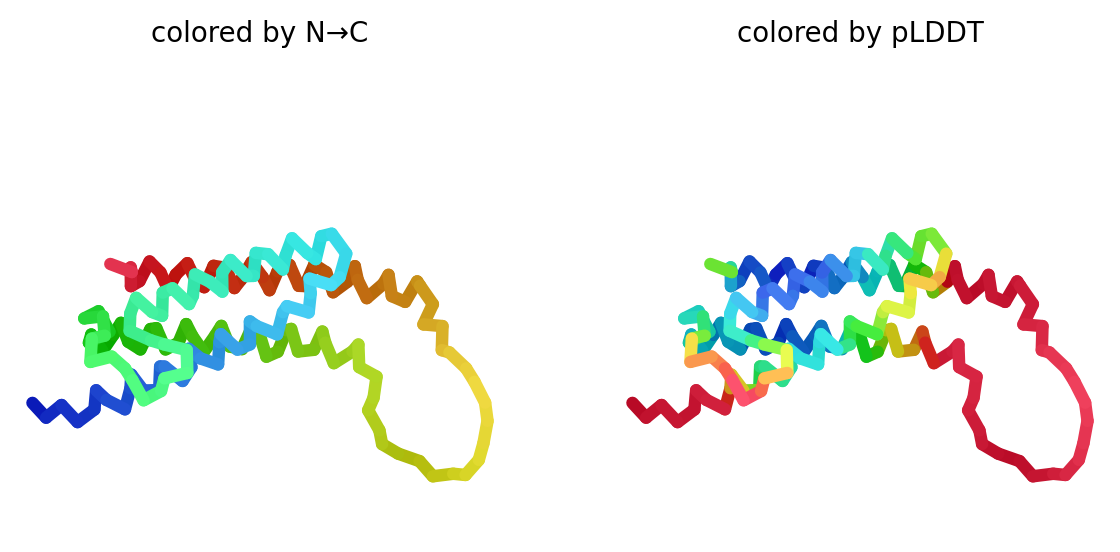

### rank_4_model_2_ptm_seed_0.png

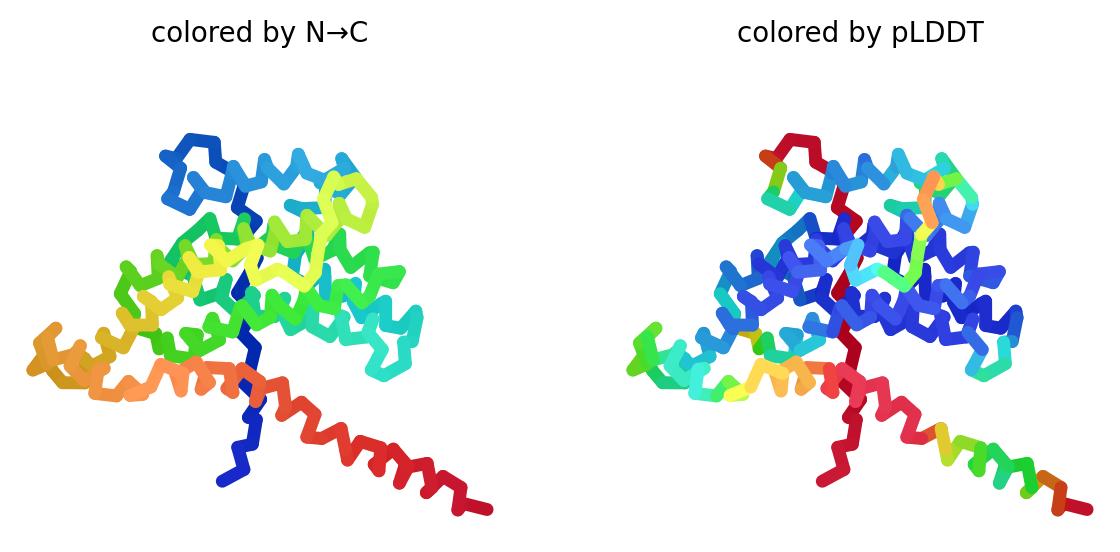

### rank_4_model_2_ptm_seed_0.png

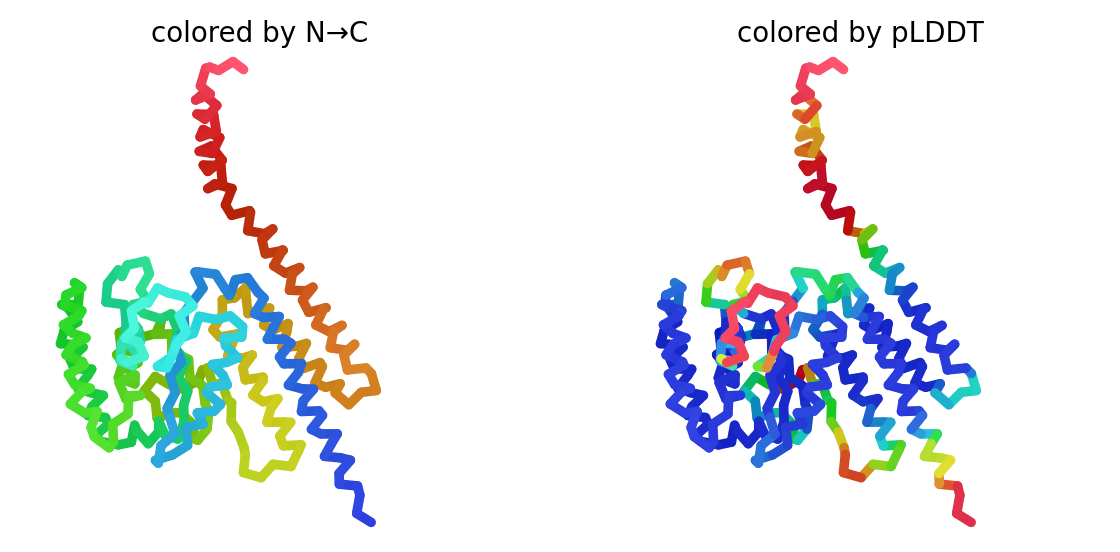

### rank_4_model_4_ptm_seed_0.png

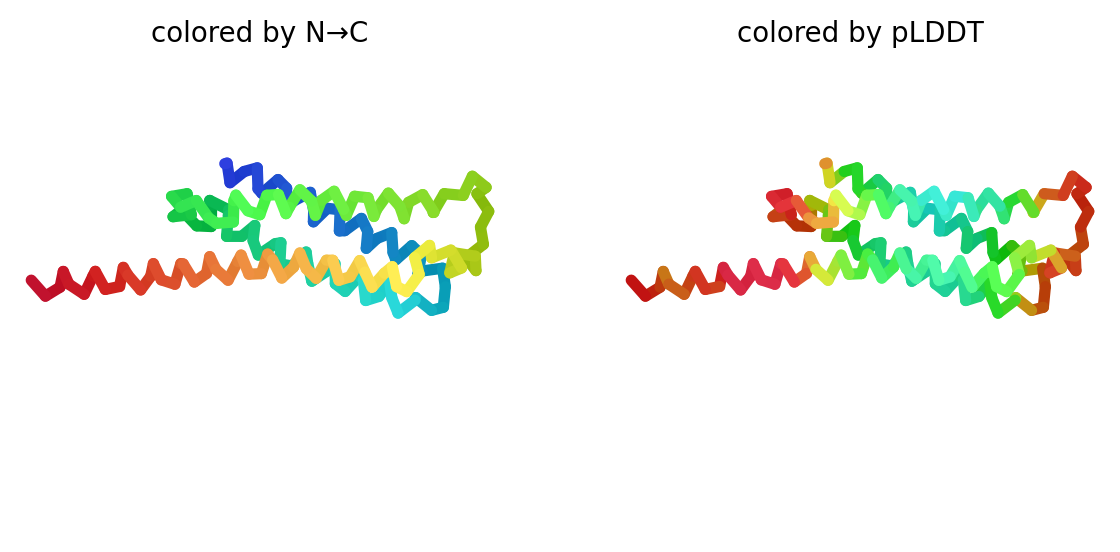

### rank_4_model_5_ptm_seed_0.png

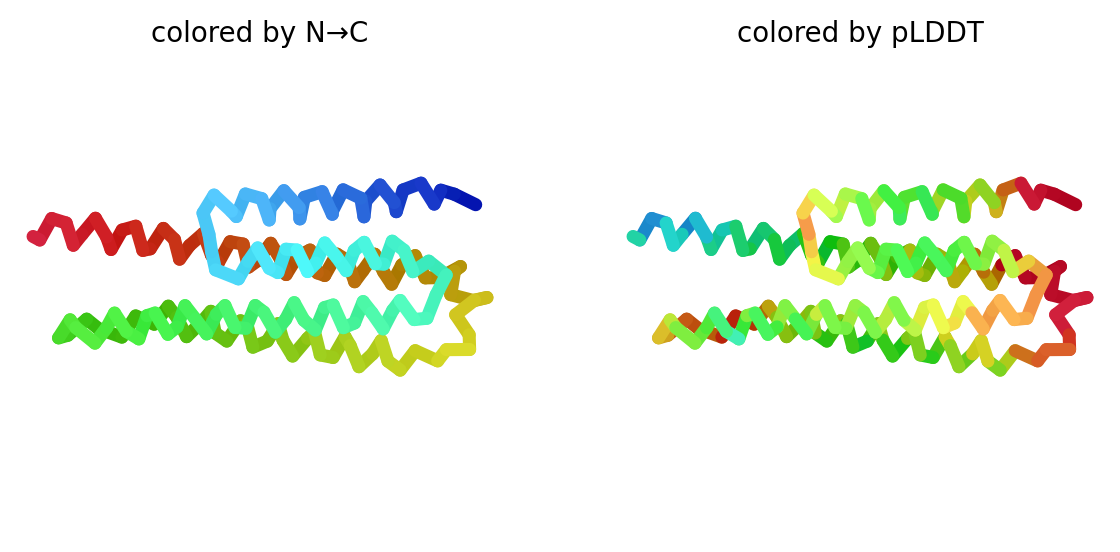

### rank_5_model_1_ptm_seed_0.png

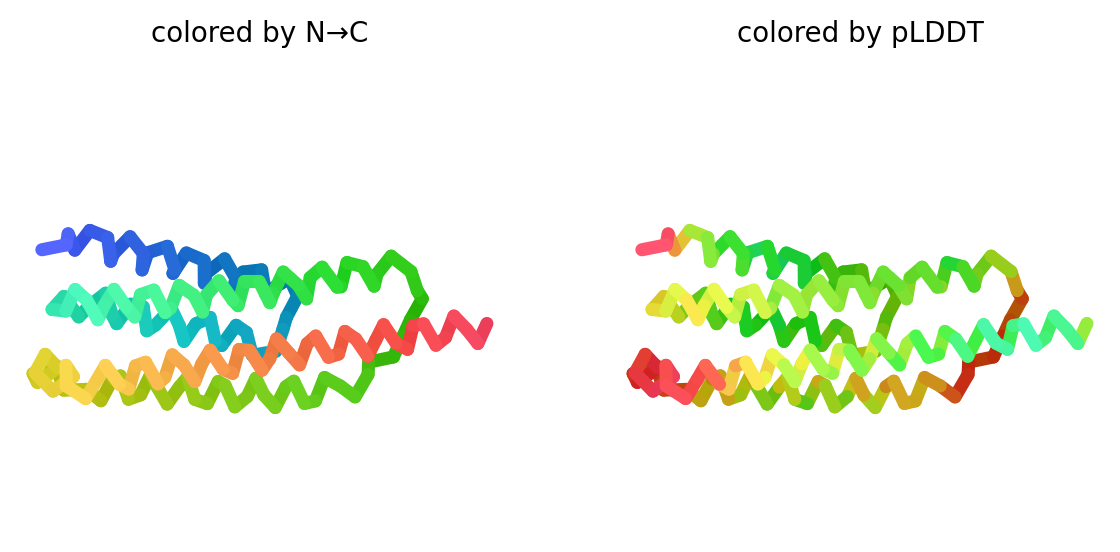

### rank_5_model_1_ptm_seed_0.png

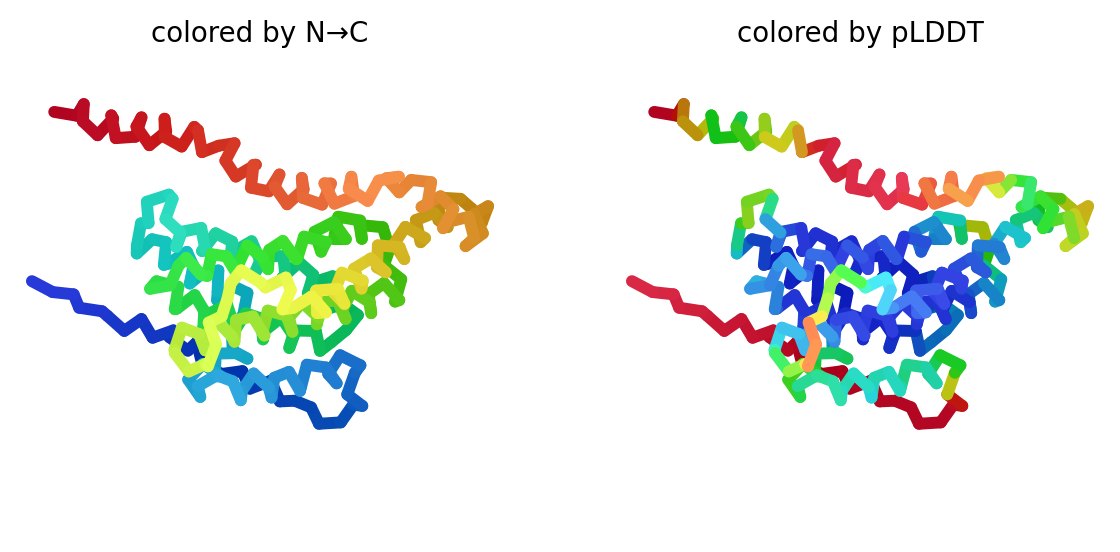

### rank_5_model_2_ptm_seed_0.png

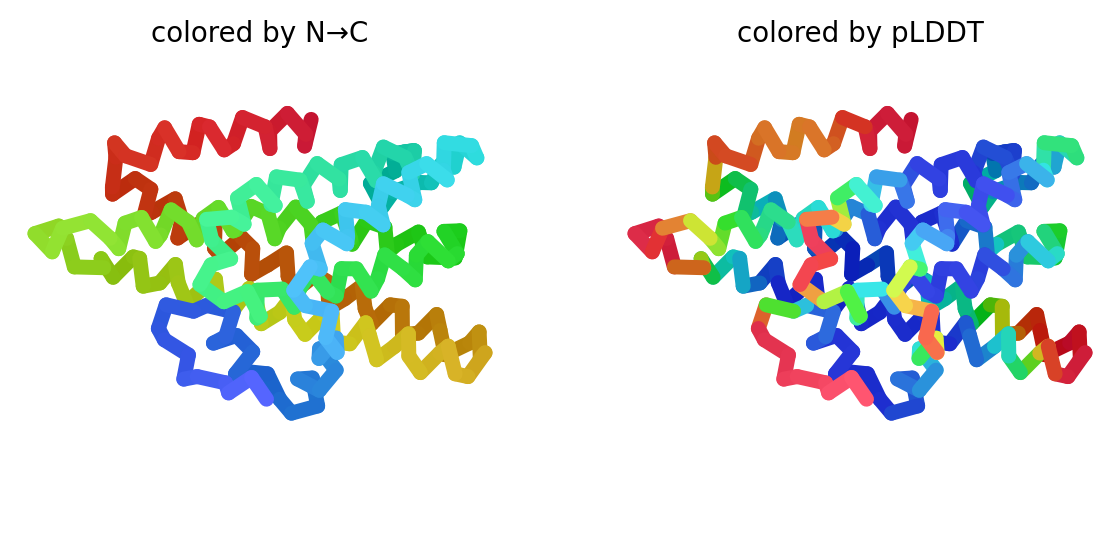

### rank_5_model_2_ptm_seed_0.png

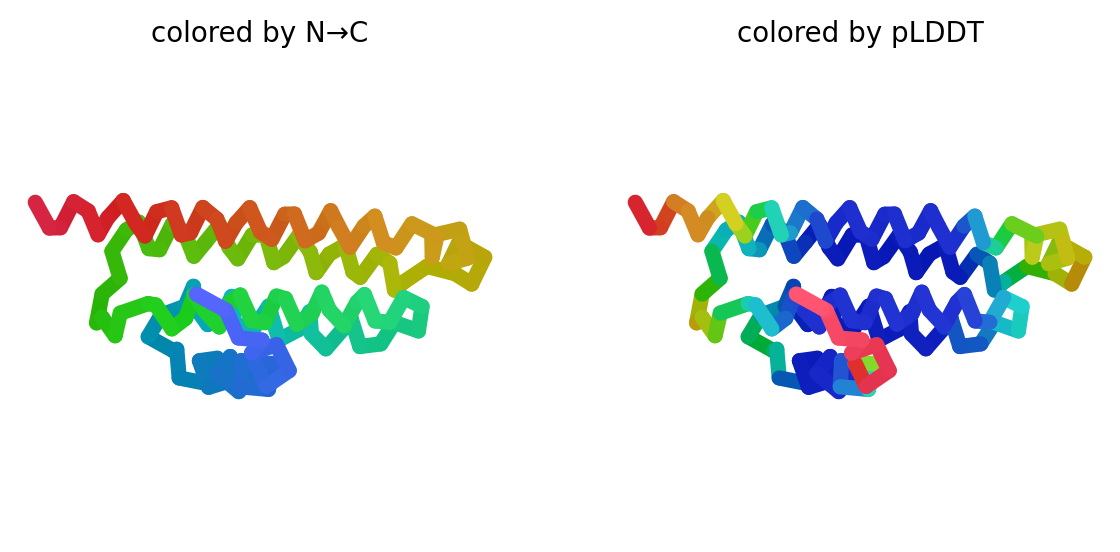

### rank_5_model_2_ptm_seed_0.png

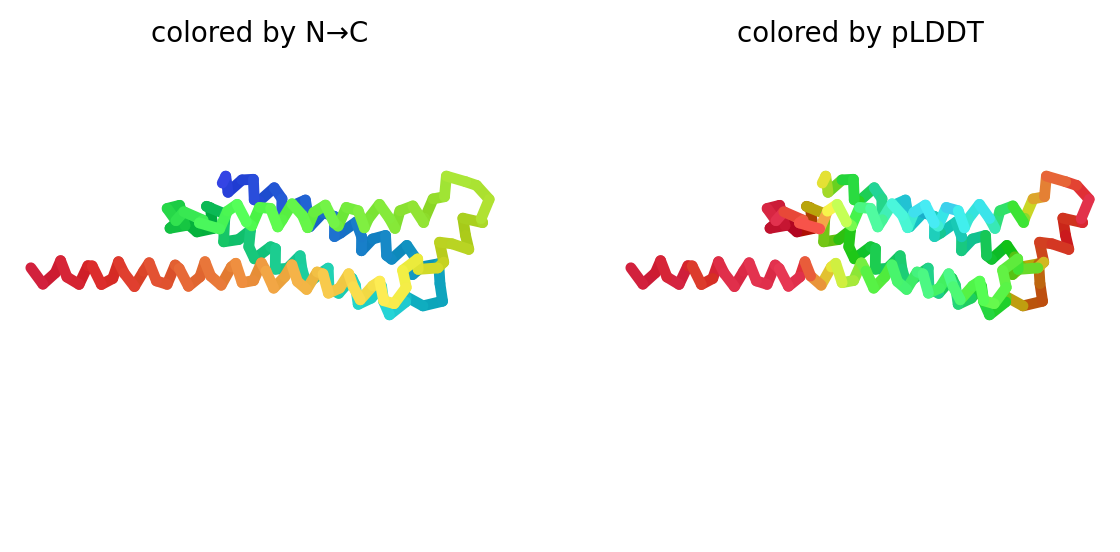

### rank_5_model_4_ptm_seed_0.png

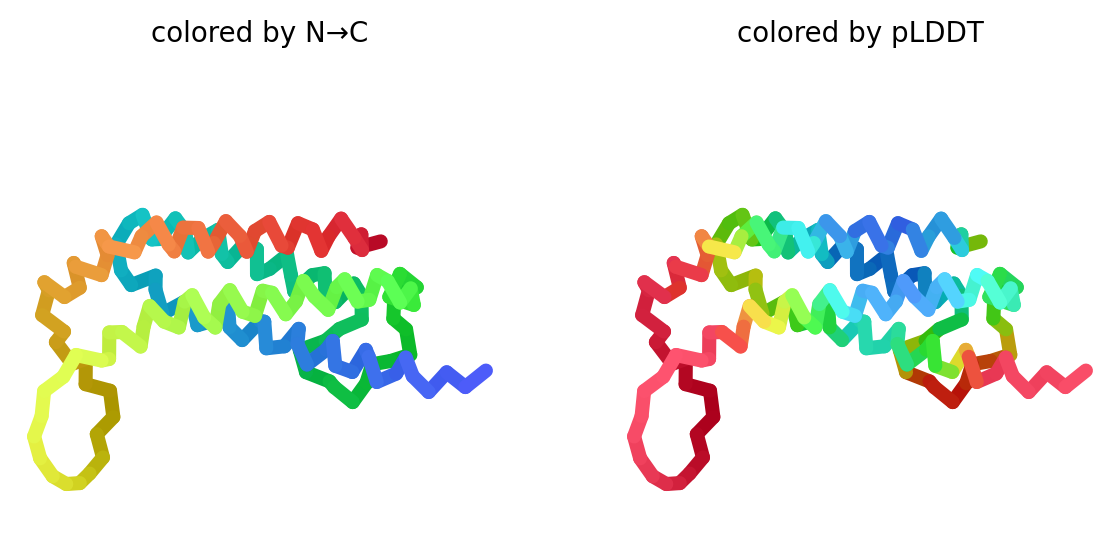

### rank_5_model_5_ptm_seed_0.png

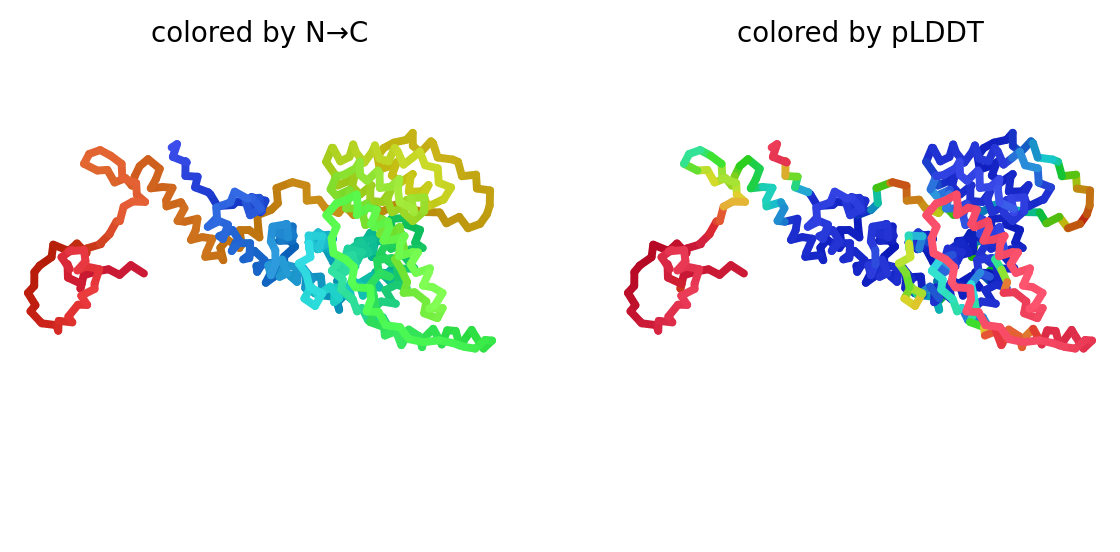

### rank_5_model_5_ptm_seed_0.png

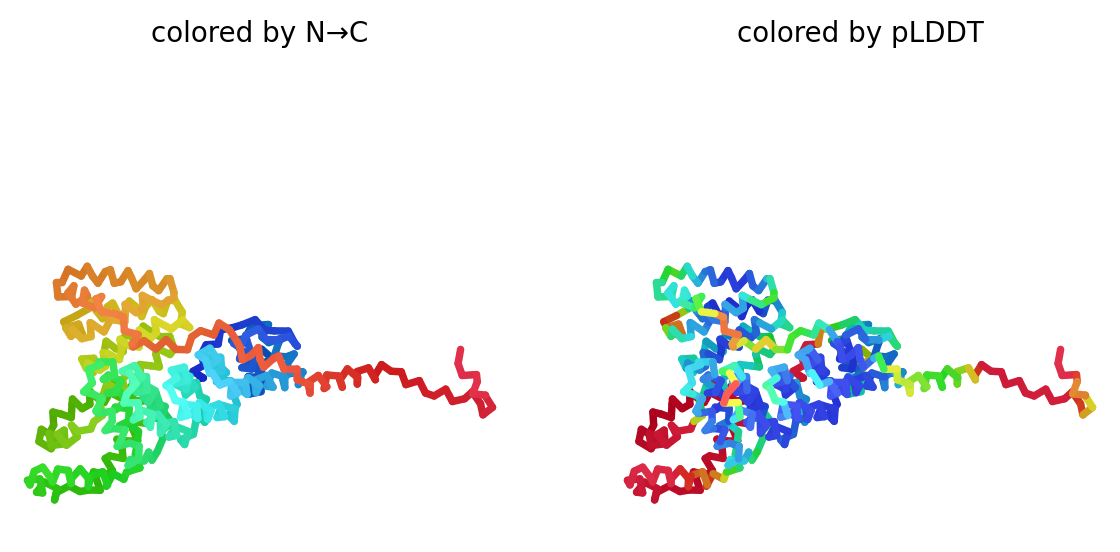

### rank_5_model_5_ptm_seed_0.png

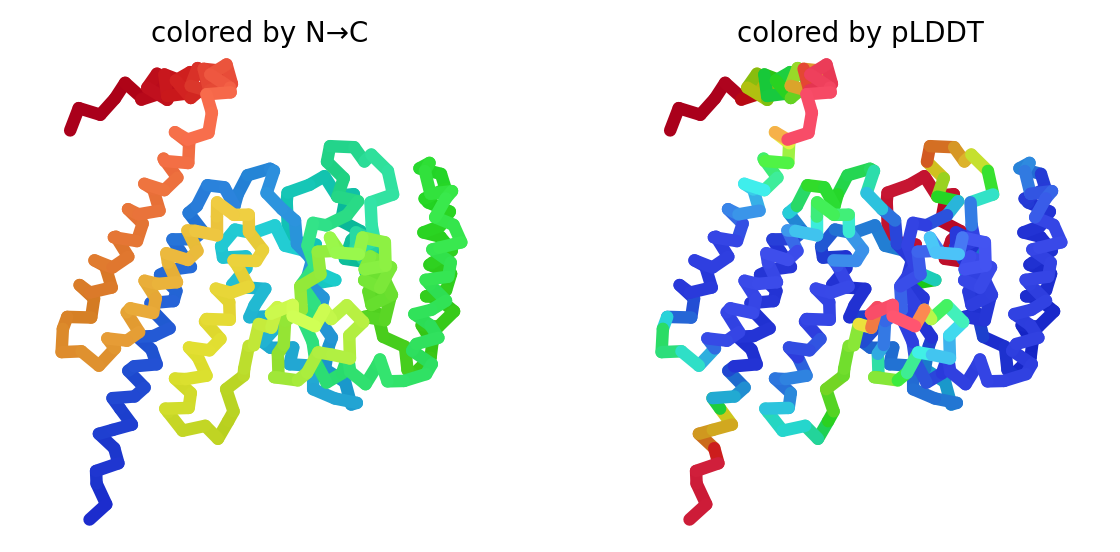
